# A Pan-Cancer Census of Dominant Tumor Immune Archetypes

**DOI:** 10.1101/2021.04.26.441344

**Authors:** Alexis J. Combes, Bushra Samad, Jessica Tsui, Nayvin W. Chew, Peter Yan, Gabriella C. Reeder, Divyashree Kushnoor, Alan Shen, Brittany Davidson, Andrea J. Barczac, Michael Adkisson, Austin Edwards, Mohammad Naser, Kevin C. Barry, Tristan Courau, Taymour Hammoudi, Rafael J Arguëllo, Arjun Arkal Rao, Adam B. Olshen, The Immunoprofiler consortium, Cathy Cai, Jenny Zhan, Katelyn C. Davis, Robin K. Kelley, Jocelyn S. Chapman, Chloe E. Attreya, Amar Patel, Adil I. Daud, Patrick Ha, Aaron A. Diaz, Johannes R. Kratz, Eric A. Collisson, Gabriela K Fragiadakis, David J. Erle, Alexandre Boissonnas, Saurabh Asthana, Vincent Chan, Matthew F. Krummel

**Author notes:** To whom correspondence should be addressed. Mail.

## Abstract

Cancers display significant heterogeneity with respect to tissue of origin, driver mutations and other features of the surrounding tissue. It is likely that persistent tumors differentially engage inherent patterns–here ‘Archetypes’–of the immune system, to both benefit from a tumor immune microenvironment (TIME) and to disengage tumor-targeting. To discover dominant immune system archetypes, the Immunoprofiler Initiative (IPI) processed 364 individual tumors across 12 cancer types using standardized protocols. Computational clustering of flow cytometry and transcriptomic data obtained from cell sub compartments uncovered archetypes that exist across indications. These Immune composition-based archetypes differentiate tumors based upon unique immune and tumor gene-expression patterns. Archetypes discovered this way also tie closely to well-established classifications of tumor biology. The IPI resource provides a template for understanding cancer immunity as a collection of dominant patterns of immune infiltration and provides a rational path forward to learn how to modulate these patterns to improve therapy.

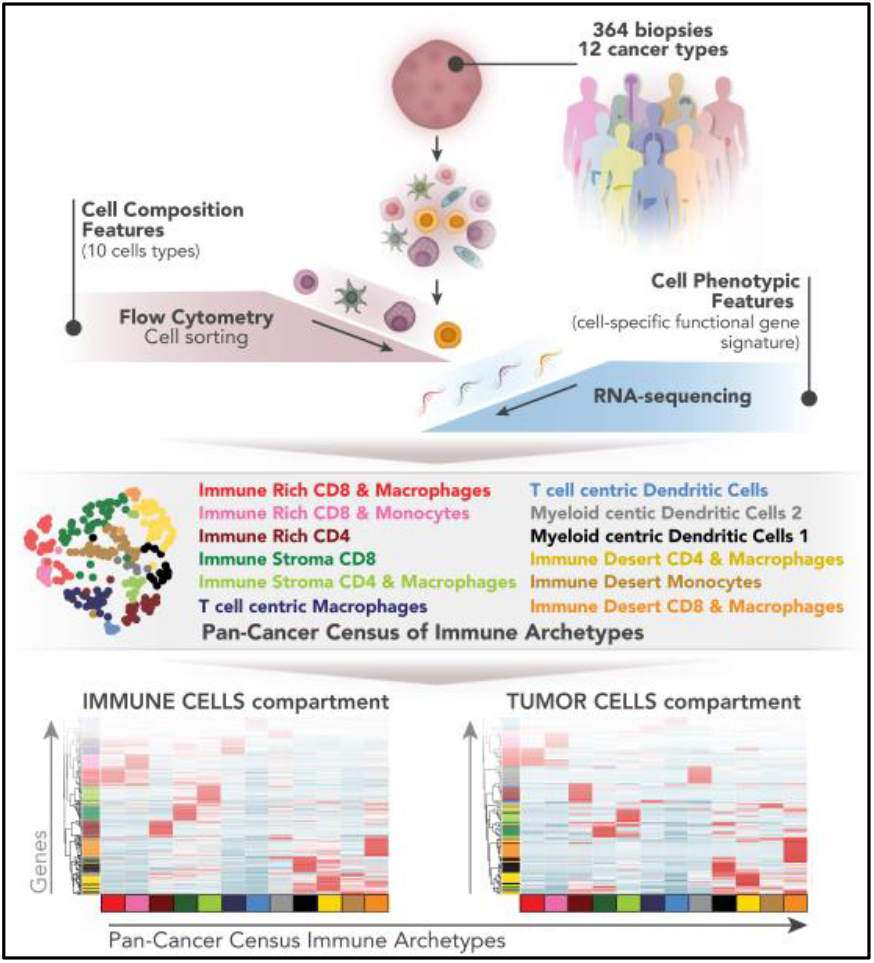

## INTRODUCTION

Pathologists have long recognized that tumors are infiltrated by cells of both the innate and adaptive arms of the immune system and thereby mirror inflammatory conditions arising in non-neoplastic tissues (Dvorak, 1986). Indeed, tumors are complex environments where malignant cells interact with both immune and nonimmune cells to form the complex cellular network of the tumor microenvironment (TME) (Hanahan and Weinberg, 2011). Cancer immunotherapy has revolutionized cancer care by acting directly on the TME and re-engaging the anti-tumor immune response (Iwai et al., 2005; Leach et al., 1996). However, the biology of the immune microenvironment driving these responses is incompletely understood (Hugo et al., 2016; Spranger, 2016) and many patients experienced minimal or no clinical benefit from immunotherapies. A deeper understanding of the diversity of the immune microenvironment across human malignancies is critical to the improvement of immunotherapy treatment strategies (Binnewies et al., 2018; Gajewski et al., 2013; Gotwals et al., 2017; Hegde et al., 2016).

To date, the combination of The Cancer Genome Atlas (TCGA) (Cancer Genome Atlas Network, 2015) and deconvolution technique base on gene expression from bulk tissue such as CIBERSORT (Aran et al., 2017; Chen et al., 2018; Newman et al., 2015) have established a foundational but low resolution landscape of the TME across human tumors (Bindea et al., 2013; Gentles et al., 2015; Mlecnik et al., 2016; Rooney et al., 2015; Thorsson et al., 2018). More recently, single-cell RNA sequencing (scRNA-seq) technologies have been increasingly applied to define the diversity of TME in a single cancer type (Azizi et al., 2018; Goswami et al., 2020; Lavin et al., 2017; Zhang et al., 2019) or focus on a single immune cell compartment (Cheng et al., 2021; Gueguen et al., 2021; Oh et al., 2020; Zhang et al., 2020; Zilionis et al., 2019). Nonetheless, the identification of recurrent motifs in the immune system that spans cancer types is still lacking.

It is now well understood that complex coordination of immune cell states is required to achieve important tissue functions such as wound healing, tissue homeostasis or viral clearance (Mujal and Krummel, 2019).  A given immune response can thus be can be conceived as a collection of cell subsets and specific immune cell pairings linked with function allowing us to define “immune archetypes. For responses to therapy, a strong archetypal relationship between cDC1 and NK cells (Barry et al., 2018; Böttcher et al., 2018) or cDC2 and CD4 conventional and regulatory T cells (Binnewies et al., 2018; Bosteels et al., 2020) have been previously identified in specific tumor types and viral infection. However, the association of dominant tumor-promoting archetypes with tumor biology is still unclear (Chen and Mellman, 2017).

In this study, we leveraged a unique dataset composed of both cell type compositional and transcriptomic data from 364 fresh surgical specimens across 12 tumors types to identify conserved tumor immune archetypes. We used an unsupervised clustering approach based on tumor specific immune gene signatures, benchmarked against ‘ground truth’ cell type composition data from flow cytometry, to identify and validate 12 unique tumor immune archetypes that could then be identified and validated in the TCGA dataset. These archetypes, discovered with only ten measurements, are well correlated to immune and tumor transcriptomic programs that span different tissues of origin and provide an unprecedented resource to classify and study cancer immunity and cancer targets to improve response to immunotherapy.

## RESULTS

### A Pan-Cancer High-Dimensional Study of Dominant Immune Composition

The UCSF Immunoprofiler (IPI) collected fresh surgical specimens from 12 distinct tissues of origin using an unbiased approach, i.e., agnostic of tumor type, stage and grade (Figure 1A, Supplementary Table 1). We performed standardized processing of 364 individual tumor specimens including rapid digestion into single cell suspension for immune phenotyping using multi-parametric flow cytometry (flow panels, see extended method). To identify patterns of gene expression within six broadly defined cell populations, we also performed cell sorting for bulk RNA-sequencing including compartments denoted: 1. “Live”: All viable cells at the time of sorting, 2. “T_conv_”: sorted conventional CD4^+^ and CD8^+^ T cells, 3. “T_reg_”: CD25^+^ CD4^+^ (enriched for regulatory) T cells, 4. “Myeloid”: Lymphocyte-negative HLA-DR^+^ (enriched for myeloid) cells, 5. “Stromal”: CD45^−^ CD44^+^Thy1^+^ cells, and 6. “Tumor”: all other CD45^−^ cells (Figure 1B, and see Extended Methods for full sort descriptions and potential caveats).This approach was chosen over single-cell RNA sequencing (scRNA-seq) because deep sequencing is needed to fully capture transcriptomic differences in sorted cell subsets populations and because scRNA-seq was an immature technology at the start of the program.

**Figure 1:**
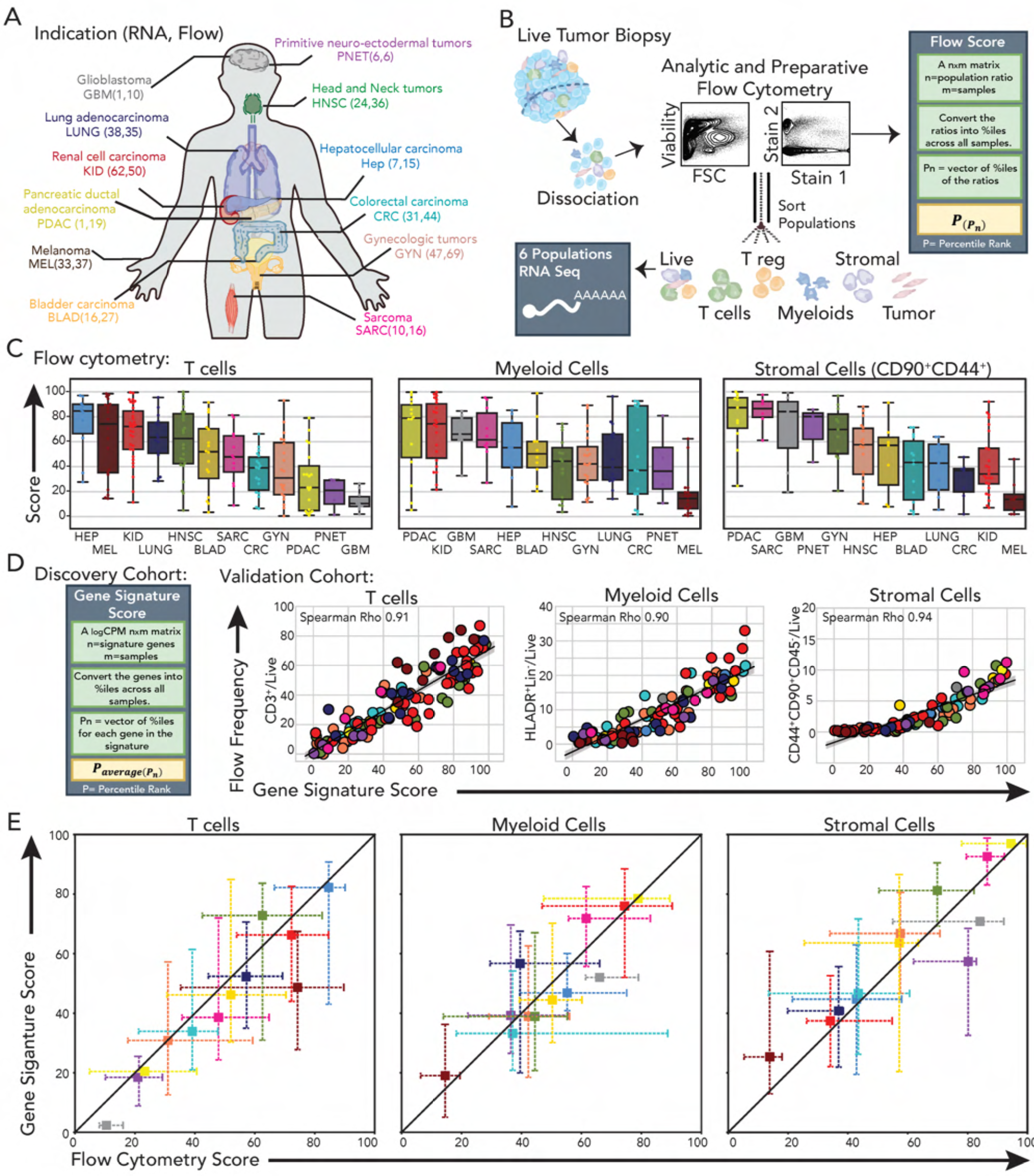
Generation and validation steps of T cells, Myeloid cells and Stromal cells features from solid tumors using flow cytometry and bulk RNA-sequencing. A. Details of the Immunoprofiler initiative (IPI) cohort tumor samples collection, color-coded by anatomical region and annotated with case numbers of bulk-RNA sequencing of viable cells sorted from fresh surgical tumor specimens and total number of samples with flow cytometry data. B. Description of the processing pipeline for digesting fresh tumor specimens into single cell suspension, submitting to multi-parametric flow cytometry for immune phenotyping and cell sorting into six different cell population compartments (live (viable cells), tcell (conventional T cells), treg (T regulatory cells), myeloid (myeloid cells, stroma (CD90+ CD44+ stromal cells), and tumor (tumor cells). C. Box and whisker plots of flow score for T cells (n=200), Monophagocytes (n=159), and Stromal cells (n=121) based on population percent in tumor specimens measured by flow cytometry (see Supplementary table S2 for details by cancer type). D. Correlation plots of Tcell, Myeloid and CD90+ CD44+ Stroma gene signature scores, for each tumor specimens, against their corresponding flow score (See Fig Supp 1 and extended method section). E. Cross-whisker plots comparing median Tcell, Myeloid and CD90+ CD44+ Stroma gene signature scores by tumor indication to the median flow score with the interquartile range on both axes. F. Individual and combined spider plots of median Tcell, Myeloid, CD90+ CD44+ Stroma scores by tumor indication.

We initially used flow cytometry and focused upon the abundance of three major cell types in the TME, namely total α/β T cells (Tcell feature), myeloid cells (Myeloid feature) and non-immune CD45^−^, CD44^+^, Thy1^+^ stromal cells (CD44^+^ CD90^+^ Stroma feature) (supplementary table 2). Consistent with previous descriptions (Mandal et al., 2016; Thorsson et al., 2018; Varn et al., 2017) the cell abundances for these three cell types, plotted as a score (Figure 1B, and extended method) for each tumor indication show tremendous heterogeneity across and within tumor indications suggesting the need for a tumor classification that goes beyond tissue of origin (Figure 1C).

A distinguishing feature of our cohort is that it has linked compositional data via flow cytometry and RNAseq, which we leveraged to discover and then validate a set of gene signatures that would allow us to infer and compare compositional data from external RNAseq-only datasets such as TCGA. We used differential gene expression (DGE) analysis to define unique gene signatures for the Tcell, Myeloid and CD44+ CD90+ Stroma features, comprised of 25, 29 and 21 genes respectively, and used this to generate a compositional score where a high value will correspond to a high cell abundancy in the tumor specimen analyzed. (Figure S1A). Use of this score, when applied to RNAseq data from the ‘Live’ RNAseq compartment in a validation cohort showed a high degree of concordance with the cell type frequencies obtained via flow cytometry, independent of tissue of origin (spearman correlation 0.91,0.90 and 0.94) (T, Myeloid, and Stromal, respectively) (Figure 1D). In addition, rank ordering of tumor indications by these gene scores shows similar trends, and strong correlations across individuals to those obtained by flow cytometric measurements (Figure 1C/S1C and 1E).

Our signatures for intra-tumoral composition significantly outperformed signatures derived from cell lines (e.g.CIBERSORT Figure S1B) when compared to the ‘ground-truth’ of flow cytometric composition data. We presume this improvement results from our data being taken from cells directly collected from fresh tumors tissue as opposed to cell lines or isolated samples (Aran et al., 2017; Newman et al., 2015). When we applied our gene signatures to RNA-seq data from 4341 TCGA tumor specimens RNA-seq data, we found that the median score for each indication between datasets for the majority of the tissues surveyed was similar, suggesting that the abundances we describe by indication (e.g. Figure 1C) extend beyond our sample processing protocol (Figure S1D). The exceptions to this were for lung (LUNG), liver (HEP) and pancreatic (PDAC) tumors and this may be due to sampling or patient-selection variation between the cohorts; thus, in the remainder of this analysis, we did not consider these indications when making additional comparisons.

### Unsupervised Clustering of 3-Features, Independent of Tissue of Origin

We next performed unsupervised clustering, using the Louvain community detection algorithm on a K nearest-neighbor (KNN) weighted graph, on these 3 features for all 260 samples with a “Live” bulk RNAseq compartment (Supplementary Table S2). The clustering was visualized using UMAP, a dimensionality reduction technique (Blondel et al., 2008; Zhu et al., 2020) (see extended method section), both on the IPI and TCGA cohorts (Figure 2A/E). The optimal clustering parameters were evaluated by minimizing the Davies Bouldin Index (DBI) (Davies and Bouldin, 1979), a metric that assesses the ratio of the intra-cluster distance to inter-cluster distance (Figure S2A-B; see Feature Clustering in Extended Methods section) and in both cohorts this resulted in six clusters. In the IPI cohort, this separated two immune rich clusters (termed immune rich (IR) and immune stromal rich (IS)) defined by high expression of Tcell and Myeloid features and differentiated by Stromal enrichment in IS. This similarly differentiated two Immune desert (ID) clusters (Immune Desert and Immune Stromal Desert) defined by low expression of Tcell and Myeloid features, and again differentiated by enrichment of CD44^+^ CD90^+^ stroma within the Immune Desert cluster. Finally, we also identified clusters of surgical specimens with enrichment in only one immune feature, namely the Tcell Centric (TC) and Myeloid Centric (MC) features (Figure 2B-D and Figure S2C left). Application of our gene scores to the TCGA cohort also produced six clusters when minimizing the DBI and these corresponded to the six clusters found in the IPI cohort (Figure 2E-F and Figure S2C-D).

**Figure 2:**
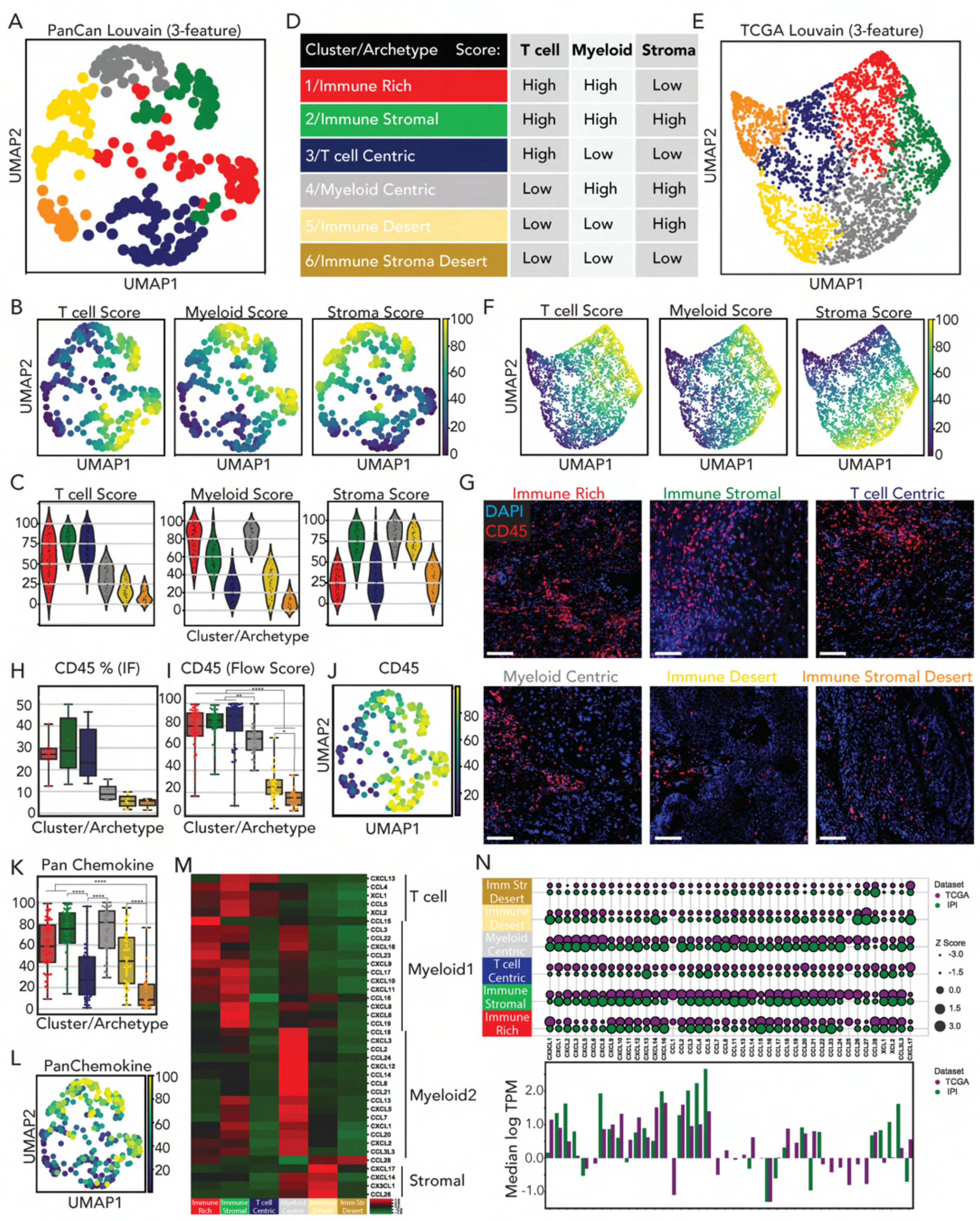
Identification of coarse immune archetypes in solid tumors using Louvain clustering on two independent datasets. A,E. UMAP display using KNN and Louvain clustering (a graph-based community detection algorithm) of tumor immune archetypes using Tcell, Myeloid and CD90+ CD44+ Stroma features to cluster patients in the IPI (A) and TCGA (E) cohorts,. Each dot represents a single patient summarized by the 3 features. B,F. UMAP overlays of the Tcell, Myeloid and CD90+ CD44+ Stroma features in the IPI (B) and TCGA (F) cohorts. C. Violin plots of Tcell, Myeloid and CD90+ CD44+ Stroma features for each cluster/archetype in IPI cohort. D. Table summarizing the six cluster/archetypes with descriptions based on the abundance of the Tcell, Myeloid and CD90+ CD44+ Stroma features. G,H. Representative Immunofluorescence of tumor specimens using CD45 (red) and DAPI (blue) staining for each cluster/archetype (G) and respective quantification of immune cell frequency (H). I,J. Box and whisker plot (I) and UMAP overlay (J) of immune cell frequency using flow cytometry. K,L. Box and whisker plot (K) and UMAP overlay (L) of a pan chemokine phenotype gene signature score. M. Heatmap and hierarchical clustering of median chemokine gene expression per cluster/archetype identified in IPI cohort. N. (top) Bubble plot of median chemokine gene expression by cluster/archetype identified in the IPI (green) and TCGA (violet) cohorts. (bottom) Bar plots of median Log TPM gene expression of each chemokine in the IPI (green) and TCGA (violet) cohorts.

To both evaluate the histological descriptions in our archetypes and provide validation of the flow cytometry data, we ran immunohistochemistry assays on tissues collected from a subset of the IPI tumor surgical specimens. Consistent with expectation, we observed that samples taken from immune rich clusters (1-3) had the highest CD45^+^ infiltration, while immune desert samples had the lowest (Figure 2G-H and Figure S2G-L). Myeloid Centric clusters were more similar to the immune desert clusters for overall CD45+ infiltration as assessed by imaging (Figure 2G-H and Figure S2I-J). However, MC clusters were intermediate when considering flow cytometry percentages which may represent a modest preferential recovery of immune cells by dissociation when compared to non-immune (Figure 2I-J). Future analyses of spatial dimensions may reveal paired infiltration patterns (i.e. T cell plus or minus myeloid or ‘excluded’ versus ‘infiltrated’ (Galon et al., 2006; Mlecnik et al., 2016), although this will involve considerable orthogonal analyses due to the intrinsic complexity of diverse tissue morphologies (e.g. in lung versus colon).

Given the importance of chemokines in recruitment of immune cells, we sought to determine whether these archetypes had unique chemokine gene-expression signatures that might corroborate their classification. We first derived a single gene signature score applied to the “Live” bulk RNAseq compartment, based on 39 chemokines (Nagarsheth et al., 2017), and found a high score in immune rich clusters and a low score in immune desert cluster in both the IPI and TCGA cohorts (Figure 2K-L and Figure S2E). Two exceptions stood out using the pan-chemokine measure: Tcell and Myeloid Centric clusters showed decreased and increased pan-chemokine scores, respectively (Figure 2K-L). We thus examined chemokines individually, using hierarchical clustering of the median chemokine expression amongst the tumors of each archetype, and identified sets of chemokines associated with each cluster (Figure 2M-N). For instance, the three T cell enriched clusters are enriched in either chemokines expressed by T cell (XCL1, XCL2, CCL4) or T cell attracting chemokines (CXCL13, CCL5) (Nagarsheth et al., 2017). Other archetypes had their own pattern of differentially expressed chemokines and these were broadly consistent with their composition. Immune desert clusters typically had a few unique and specific chemokines expressed. The patterns observed were similar on a chemokine-by-chemokine basis for archetypes as defined by composition in TCGA (Figure 2N), using the gene-signature archetype delineation (Figure 2E). Taken together, these results demonstrate that a three-feature scoring of tumor tissue identifies six unique clusters strongly correlated with immune infiltration and expression of distinct sets of chemokines.

### Distribution of 3-feature Archetypes by Indication and outcome

Assessing archetypes based on just these three features demonstrated a heterogenous distribution of tumor indications among the different clusters, although many indications, such as Kidney and Melanoma had significant biases (Figure 3A and Figure S3A). Taking advantage of the large clinical dataset available in the TCGA cohort, we sought to broadly assess whether these simple archetypal classifications had a relationship to prognosis. Agnostic to the tissue of origin, the overall survival at 5 years is significantly better for the immune rich archetype compared to all the others (p-value 3.9E^−11^) with a general trend in outcome that tracks with overall immune infiltration (Figure 3B). To extend this analysis and focus within indications, we analyzed distributions of archetypes in both IPI and TCGA and also assessed the outcome in individual indications (Figure 3C and Figure S3B-D). While the relative composition of archetypes by indication were broadly similar in both cohorts, there was some variation e.g. heavily ‘3 Tcell Centric’ bias in Melanoma in IPI cohort manifests as a combination of these and ‘1:IR’ or ‘IS’ in TCGA. Furthermore, while some indications such as colorectal cancer appears to have slightly better prognosis for ‘IR’ archetypes, trends were weaker within most individual indications. This variability prompted us to consider that these coarse-grained three-feature archetypes may not capture the full range of immune cell heterogeneity in tumors. Heterogeneity was especially evident in features such as Neutrophil density, Treg and CD4^+^/CD8^+^ T cell ratio which all showed high variability within 3-feature archetypes (Figure 3D-F).

**Figure 3:**
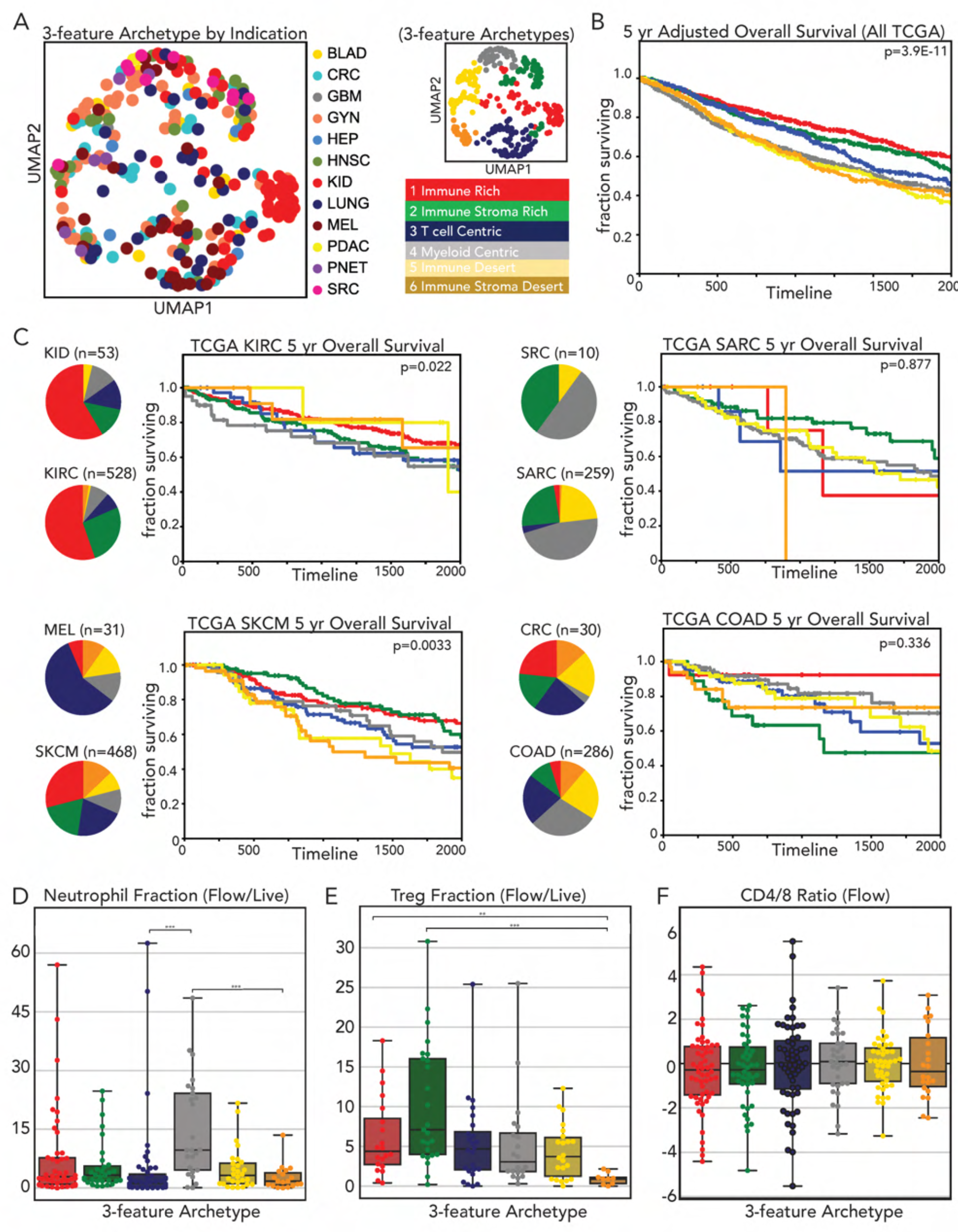
Coarse immune archetypes are independent of tissue origin and associated to overall survival. A. UMAP display and graph-based clustering of tumor immune archetypes using Tcell, Myeloid and CD90+ CD44+ Stroma features color-coded by tumor indication. B. Kaplan-Meier overall survival curves for each immune tumor archetype identified in the TCGA cohort. C. (Left) Pie charts representing distributions of each archetype by indication in the IPI (top) and TCGA (bottom) cohorts. (Right) Kaplan-Meier overall survival curve for each immune tumor archetype identified in the TCGA cohort for Kidney renal clear cell carcinoma (KIRC), Skin cutaneous melanoma (SKCM), Sarcoma (SARC) and Colon adenocarcinoma (COAD). D. Box and whisker plot of Neutrophils frequency in tumor measured by flow cytometry for each cluster/archetype identified. E. Box and whisker plot of CD4+ regulatory T cells frequency in tumor measured by flow cytometry for each cluster/archetype identified. F. Box and whisker plot of Ln CD4+ to CD8+ conventional T cell frequency ratio in tumor measured by flow cytometry for each cluster/archetype identified using three features.

### Developing A 6-Feature Archetype Definition

Using flow cytometry data, we again grouped tumors by indication to assess variation in regulatory (Treg), CD4^+^ and CD8^+^ conventional T cell frequencies amongst T cells, both between and within indications (Figure 4A) (supplementary table 3). Repeating our previous methodology (Figure S1), we used DGE between the Treg RNAseq compartment and the other cell sorted RNAseq compartments to generate Treg gene signature composed of 9 genes, (Figure S4A) that was highly similar to other published signatures (Arce Vargas et al., 2018; Plitas et al., 2016; Zemmour et al., 2018). To isolate a signature for CD8 versus CD4 within our T_conv_ RNAseq compartment, we used the flow data to identify samples rich in CD4 or CD8 conventional T cells and performed DGE between them (Figure S4B, and extended method). Notably, most of the identified genes have been previously associated with CD4^+^/CD8+ conventional T cell identity (e.g., CD8A, IL7R) or tissue residency (e.g., VCAM1, BACH2) (Richer et al., 2016; Savas et al., 2018) (Figure S4A-B). The Treg, CD4, and CD8 feature scores, assessed in a selected validation cohort, showed very high correlation (Spearman correlation 0.86,0.97 and 0.98, respectively), with their respective population abundances measured by flow cytometry. Again, these correlations were significantly better than correlations of cell-line derived signatures on the same tumor specimens (Figure S4C). Unsupervised clustering of all samples using the CD4 and CD8 feature signature genes within our T_conv_ RNAseq revealed the existence of at least 3 distinct groups of tumors across indications: CD8-biased tumors, CD4-biased tumors and a large population that was mixed for both these signatures (Figure 4B).

**Figure 4:**
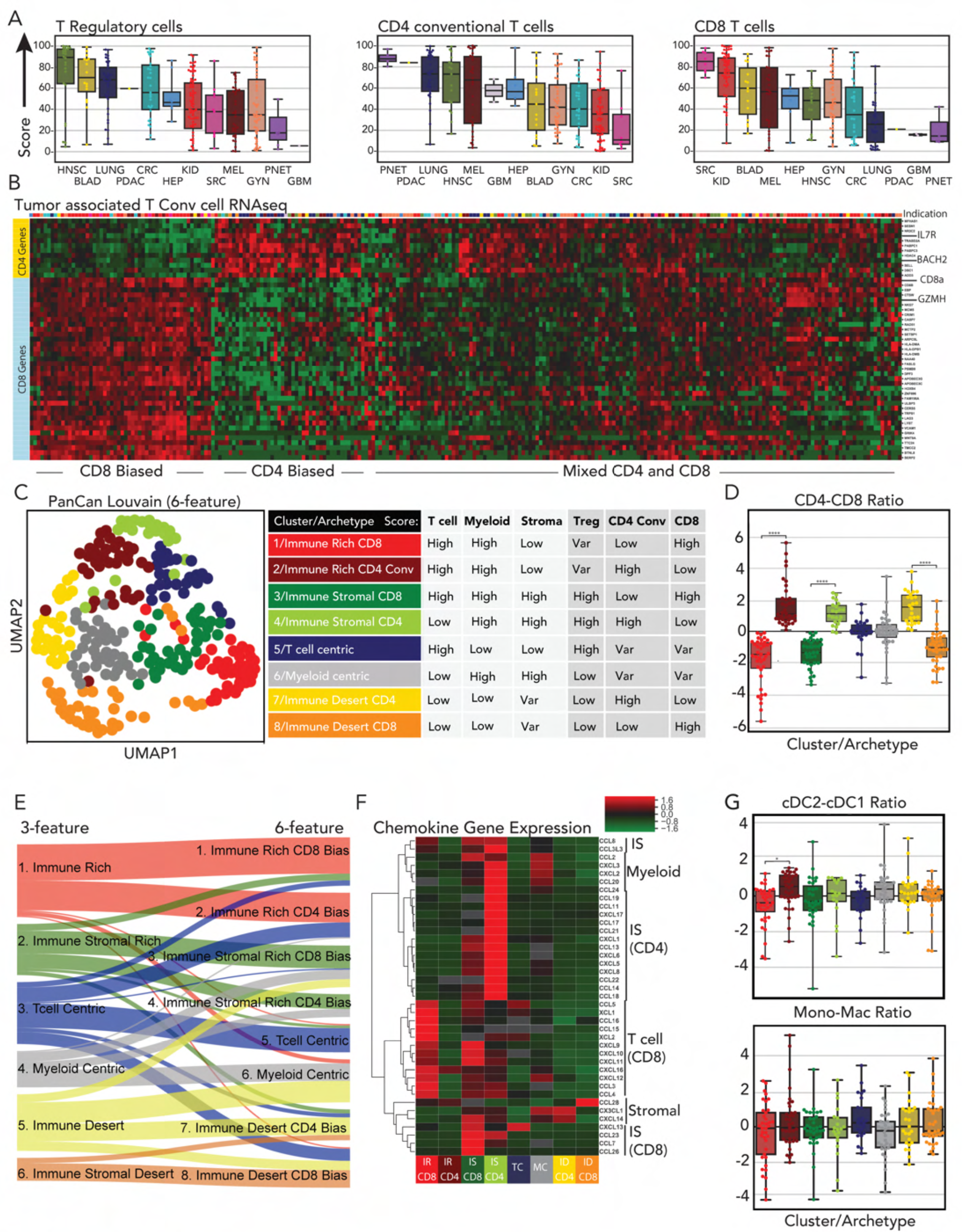
Inclusion of T cell subset features subdivide immune archetypes by CD4 to CD8 ratio. A. Box and whisker plots of feature gene signature scores for CD4+ regulatory T cells (Treg feature) out of the live compartment, CD4+ and CD8+ conventional T cells (CD4 and CD8 features) out of the tcell compartment of patients in the IPI cohort. B. Heatmap and hierarchical clustering of CD4 (yellow) and CD8 (blue) feature genes’ normalized expression, for patients in the tcell compartment. C. (Left) UMAP display and graph-based clustering of tumor immune archetypes using Tcell, Myeloid, CD90+ CD44+ Stroma, CD4, CD8 and Treg features to cluster patients in the IPI cohort. Each dot represents a single patient summarized by the 6 features. (Right) Table summarizing the eight cluster/archetypes with descriptions based on the abundance of the Tcell, Myeloid CD90+ CD44+ Stroma, CD4, CD8 and Treg features. D. Box and whisker plot of Ln CD4+ to CD8+ conventional T cell feature gene signature score ratio in tumor measured each of the cluster/archetype identified with 6 features clustering. E. Alluvial plot depicting how cluster/archetype membership perpetuates or subdivides from 3 to 6 feature clustering. F. Heatmap and hierarchical clustering of the median chemokine gene expression for each cluster/archetype identified in the IPI cohort using six features. G. Box and whisker plot of Ln cDC2 to cDC1 ratio (top) and Mono to Macs ratio (bottom) measured by flow cytometry for each cluster/archetype identified using 6 features.

We next combined the CD4, CD8 and Treg gene signature scores with the Tcell, Myeloid and CD44+ CD90+ Stroma gene signature scores to perform 6-feature clustering (Figure 4C and Figure S4D-F). DBI optimization yielded eight clusters and analysis by alluvial plot revealed that the two new clusters formed are partly a subdivision of the previous coarse Immune Rich and Immune Stromal Rich archetypes, now delineated by CD4 to CD8 ratio (Figure 4D-E and Figure S4E). This increase in feature granularity also resulted in some specimens shifting within the classification. For example, some Tcell Centric samples now shifted to being considered “Immune-rich:CD4” because of their profound high CD4 score, Furthermore, this analysis revealed that the 3-feature cluster of ‘Immune Stromal Desert’ was relatively CD8 rich whereas the ‘Immune Desert’ favored CD4 cells. CD45 densities remain high for the immune rich tumors and neutrophils remain generally variable within and between archetypes (Figure S4G-H). Assessing chemokine gene expression showed that this re-clustering dramatically segregated chemokine gene expression in the Immune Rich archetype and also further refined the chemokine expression found amongst the other archetypes (Figure 4F and Figure S4I)

### Mapping T Cell Exhaustion and Myeloid Subset Heterogeneity In 6-Feature Archetypes

Exhaustion in T cells (T_ex_) represents a transcriptional state for T cells that arises in cancers and chronic viral infections and is characterized by progressive loss of effector functions, high and sustained inhibitory receptor expression, and acquisition of a distinct metabolic and transcriptional program (Blank et al., 2019). We used the T_conv_ RNA-seq compartment to identify genes that showed the highest correlation with CTLA4, PDCD1, HAVCR2, CD38 and LAG3, previously identified exhaustion markers. We identified 11 such genes, which included TOX, a transcription factor recently described as key driver of T cell exhaustion (Beltra et al., 2020; Khan et al., 2019; Scott et al., 2019) (Figure S4J and Extended Method). We then benchmarked this gene signature score against the abundance of CD4^+^, CD8^+^ and the sum of both CD4+ and CD8+ T cells co expressing CTLA4, PD-1 and CD38 as markers of T_ex_ (Figure S4K). This T_ex_ gene score best mirrored the flow-cytometry-based abundance of CTLA-4+/PD-1+ within the combined CD8+ and CD4+ compartments (Figure S4L). Using this T_ex_ gene signature score we observed enrichment of exhaustion in CD8 biased archetypes including the immune desert (ID) archetype (Figure S4M-N) and with low MHC I expression by the tumor cells in these archetypes (Figure S4O).

In addition to T cell exhaustion, we also sought to assess myeloid heterogeneity in our tumor landscape, widely known to be variable in tissues. We thus probed the abundance of mononuclear phagocytic cell (MPC) subsets including monocytes (Mo), macrophages (Mp), classical dendritic cells (cDC2, cDC1) and plasmacytoid dendritic cells (pDCs) for each archetype using flow cytometry (Figure S4P). This revealed that, despite, slight enrichment between archetypes e.g. of cDC2 over cDC1 in IR CD4 Bias, the frequencies of cDC1/cDC2, monocytes, and macrophages showed variability within archetypes, indicating that MPC frequencies are segregating the 6-features archetypes. (Figure 4G and Figure S4Q).

### 10-Feature Archetype Definition

The complexity of the MPC identity and plasticity has complicated efforts to determine which populations are beneficial or subversive to the anti-tumor response (Broz et al., 2014; Etzerodt et al., 2020; Gubin et al., 2018). To assess myeloid diversity across all samples, we generated a scRNAseq sub-study of sorted tumor associated MPC to identify specific gene signatures for the principal MPC subsets (Figure S5A). Following removal of non-MPC cellular contaminants, we found 5 unique clusters from 3,880 input cells (Figure S5B). Using DGE between each cluster, we identified unique 5 gene signatures for each subset (Figure S5C). Most of the genes identified have been previously associated to their respective MPC subsets such as VCAN, TREM2, CD1C, CLEC9A and LAMP5 in monocytes, macrophages, cDC2, cDC1 and pDCs, respectively (Binnewies et al., 2018; Cheng et al., 2021; Combes et al., 2017; Molgora et al., 2020; Sancho et al., 2009). When validated against cellular abundance by flow cytometry, the correlation of each MPC subset feature score across all indications were highly correlated (Spearman correlation of 0.86, 0.91, 0.93, 0.94 and 0.94 for Monocytes, Macrophages, cDC2, cDC1 and pDC, respectively) (Figure S5D and extended method). Again, when we analyzed the distributions of these five MPC subset features scores by indication, a high heterogeneity by indication was apparent (Figure 5A and Figure S5E).

**Figure 5:**
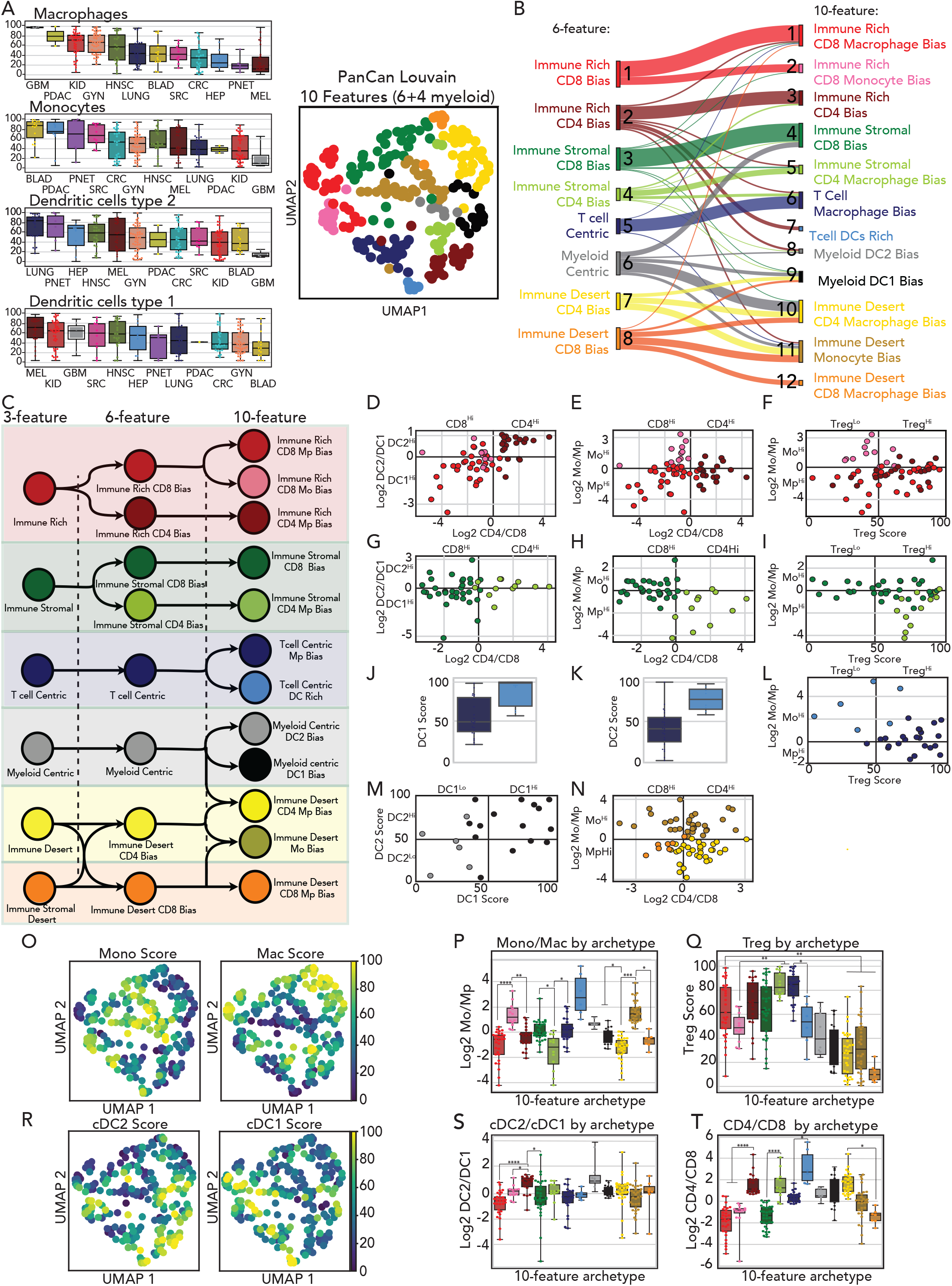
Single-cell RNA sequencing-derived myeloid signatures refines immune archetypes. A. (Left)Box and whisker plots for the Macrophages, Monocytes, cDC1 and cDC2 feature scores, calculated in the myeloid compartment, in the IPI cohort. (Right) UMAP display and graph-based clustering of tumor immune archetypes using Tcell, Myeloid, CD90+ CD44+ Stroma, CD4, CD8, Macrophages, Monocytes, cDC1 and cDC2 feature scores to cluster patients in the IPI cohort. B. Alluvial plot depicting how cluster/archetype membership perpetuates or subdivides from 6 to 10 feature clustering. C. (Left) Schematic of a “phylogeny” of the cluster/archetypes as they progressed from 3-feature to 6-feature to 10-feature clustering. D-N. Scatter plots of different features defining the twelve clusters/archetypes identified in the IPI cohort using 10 features. O, R. UMAP overlay of Macrophages (Macs) and Monocytes (Mono) and classical dendritic cell type 1 (cDC1) and 2 (cDC2) feature scores (R) (See Fig S5H) for each cluster/archetype identified using 10 features. P, Q, S, T. Box and whisker plot of Ln Mono to Macs ratio (P), Treg feature gene score (Q) Ln cDC2 to cDC1 ratio (S) Ln CD4 to CD8 conventional T cells ratio (T) for each cluster/archetype identified using 10 features.

Thus, we repeated the unsupervised clustering after adding four MPC subset features including macrophages, monocytes and the two types of classical dendritic cells to the previous six features (Supplementary Table 3). The 10-feature clustering produced 12 unique tumor immune archetypes after DBI minimization, with no bias toward specific archetypes found between scores calculated from flow and scores directly from RNAseq (Figure 5A and Figure S5F-H). As previously observed when adding T cell subsets, the addition of the four MPC measures subdivided preexisting immune archetypes but did not increase those by 16-fold as could occur if these were randomly assorted (Figure 5B-C). MPC inclusion also resulted in some specimens shifting between MC and ID archetypes, driven by strong monocytes or macrophages enrichment.

We found that generally both IR and ID sub-archetypes were distinguished by specific pairings of T cell and myeloid subsets but that the specific pairing varied. Specifically, the monocyte/macrophage ratio demarcates two different IR CD8 bias archetypes where Treg abundance is generally higher in macrophage-enriched tumor specimen (Figure 5D-F and 5O-Q). One archetype, IR CD4 biased, is differentiated from its IR CD8 counterparts by enrichment in cDC2 vs cDC1 (Figure 5D and 5R-T). However, such correlation between CD4/CD8 and cDC2/cDC1 ratios is only observed among IR archetypes and not in archetypes containing high stromal densities (Figure 5G-I and 5R-T).

The relationship between monocytes/macrophages and T cell subsets composition also differed significantly depending on archetypes (Figure 5D-N). In the IS archetypes (Figure 5H) CD8 abundance is opposed to Macrophage abundance, but in ID, both CD4 and CD8 rich sub-archetypes are equivalently enriched in macrophages and monocytes (Figure 5N, 5P and 5T). TC (Tcell centric) archetypes are characterized by a positive correlation between Treg and Macrophage abundance (Figure 5J-L and 5P-Q). This discordance between the abundances of macrophages and other immune cells in different archetypes could suggest that a generalized Macrophage score does not capture critical heterogeneity in macrophage phenotypes or that macrophages and T cells may engage in additional as-yet-undiscovered relationships that regulate their numbers.

### Defining Immune Gene Expression Pattern Across The 10-Feature Archetypes

Focusing on 10-feature archetypes, we next surveyed the association of the archetypes that emerged, with cell fractions and gene sets that were not used for clustering. Significant archetype-specific enrichment was found for gene sets that define intra-tumor NK cells in IR based on published signatures (Barry et al., 2018) (Figure 6A). Plasma cells, defined by enrichments of IgG genes, were notably also enriched in T cell Centric Macrophage bias tumors whereas B cell genes (CD20, CD79) were more prevalent in T cell Centric DC rich (Chen et al., 2020; van Galen et al., 2019) (Figure 6A). These may correspond to a prevalence for tertiary lymph nodes in those archetypes although we did not observe higher DC frequencies in those tumor specimens (Figure S5H). An archetype-specific immune composition pattern again correlated with specific enrichment of groups of chemokine transcripts. (Nagarsheth et al., 2017; Tokunaga et al., 2018) (Figure 6B). Notably, chemokines with specificity for families of chemokine receptors now frequently co-clustered within 10-feature archetypes.

**Figure 6:**
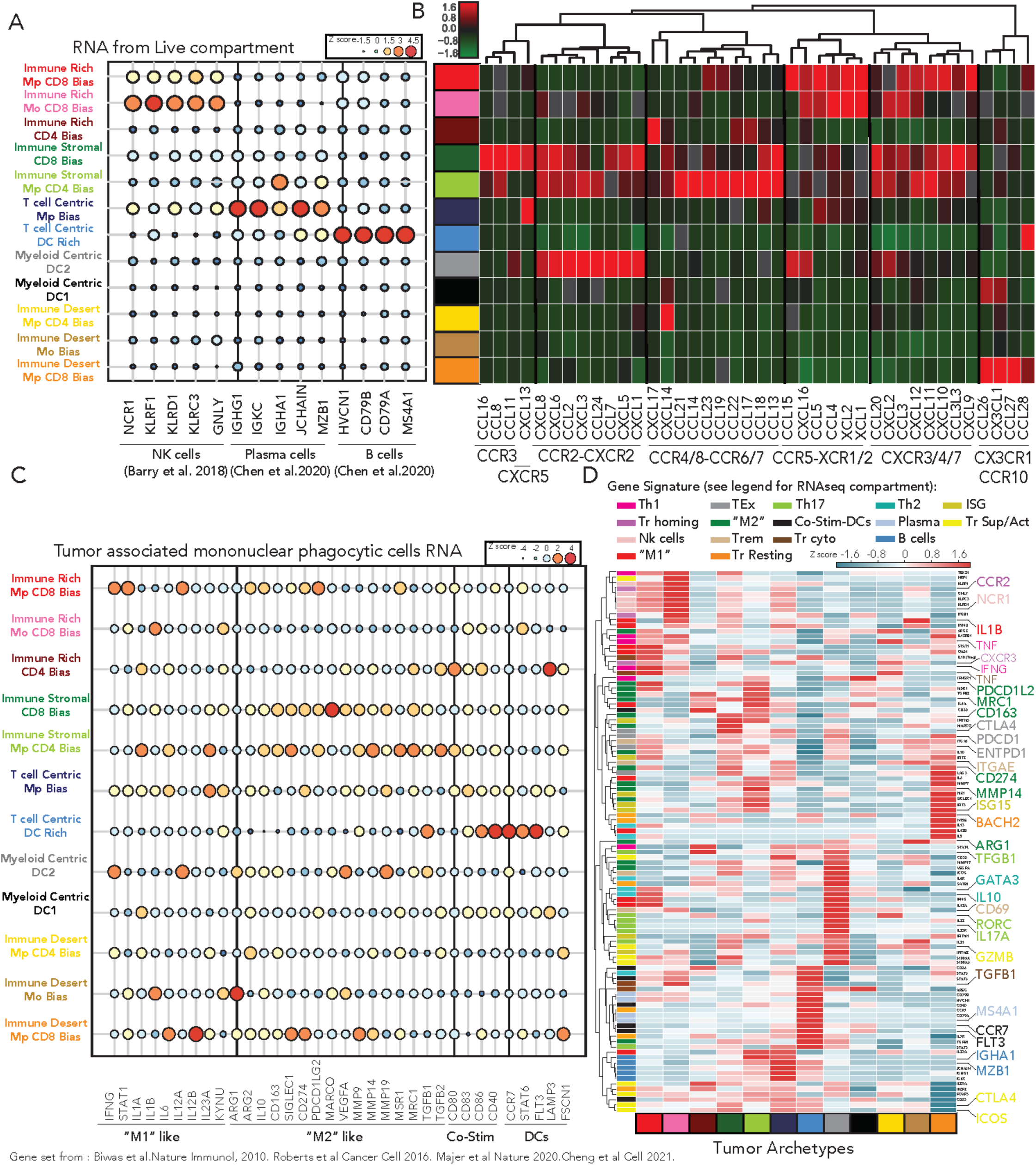
immune gene expression pattern specific to each tumor archetype. A. Bubble plot of NK cells (natural killer cells), B cells and plasma cells associated gene expression in the live compartment grouped by cluster/archetypes in the IPI cohort using 10 features. B. Heatmap and hierarchical clustering of the median chemokine gene expression of all chemokines in the Chemokine phenotype signature, grouped by cluster/archetype in IPI cohort using 10 features. C. Bubble plot of gene expression in Macrophages (M1, M2) and Dendritic cells (Co-Stim, DC) function in the myeloid compartment, grouped by cluster/archetypes in the IPI cohort using 10 features. D. Heatmap and hierarchical clustering of the median gene expression of B cells (all viable RNAseq compartment), NK cells ((all viable RNAseq compartment), plasma cells (all viable RNAseq compartment) T cells phenotypes (Tconv RNAseq compartment, T regs(Treg RNAseq compartment), macrophages and dendritic cells function (Myeloid RNAseq compartment) grouped by cluster/archetype in IPI cohort using 10 features.

We next used the myeloid RNAseq compartment to explore the level of expression of genes previously associated to tumor-infiltrating MPC, namely ‘M1’,’M2’, ‘DC’ and ‘Co-stimulatory molecules’ (Biswas et al., 2013; Cassetta et al., 2019; Cheng et al., 2021; Maier et al., 2020; Roberts et al., 2016) across all 12 archetypes. Despite the heterogeneity in individual gene expression associated to ‘M1’ and ‘M2’ macrophage phenotype among the different archetypes we observed a general enrichment in ‘M2’ genes in IS archetypes (Figure 6C). On the other hand, ‘DC’ and ‘Co-Stim’ associated genes were highly expressed in the Tcell Centric DC rich archetype with the exception of LAMP3 which was significantly enriched in the IR CD4 archetype (Figure 6C). LAMP3 expression has been recently associated to a distinct tumor-associated cDCs subset characterized by high expression of regulatory molecules such as PD-L1 and correlating with Treg abundance in tumor (Cheng et al., 2021; Maier et al., 2020; Zhang et al., 2019).

To further elucidate the heterogeneity found in myeloid function we combined the myeloid ‘signature’ gene sets with a selection of 12 other gene sets highly linked to subsets of Treg, T_conv_, B cell, plasma cells, and NK cells and performed hierarchical clustering of these genes sets based on their median expression in the archetypes. The gene sets were evaluated in different RNAseq compartments corresponding to their appropriate cell type, namely ‘Th1’ genes in the T_conv_ RNAseq compartment, ‘M1’ genes in the myeloid RNAseq compartment and ‘Treg homing’ in the Treg RNAseq compartment (Figure 6D and Supplementary table 3). Combining the gene sets in this way revealed distinct immune signatures for each archetype (Figure 6D). For instance, while the IR CD8 macrophage bias archetype was enriched in genes associated with the type 1 response (IFNG, TNF, IL1B) both in T_conv_ and MPC compartments, both IS and ID CD8 archetypes were characterized by their unique combination of upregulated gene expression associated to T cell exhaustion (PDCD1, CTLA4, ENTPD1) together with ‘M2’ like macrophages (PDCD1L2, CD163, CD274, and MRC1). This analysis also identified putative gene expression interactions present in the same archetype across cell types, such as high CCR2 expression by Tregs in an archetype rich in monocytes, which are known producers of the CCR2 ligand, CCL2 (Loyher et al., 2018; Mondini et al., 2019) (Figure 6D). Taken together, this demonstrated that the 12 tumor immune archetypes represent a collection of distinct immune phenotypes made up of a unique combination of cell composition and transcriptomic profiles.

However, certain immune populations remained variable across the archetypes. For instance, neutrophil infiltration, presents only a slight and statistically insignificant rise in archetypes with low immune infiltration (Figure S6A-B). Conversely, a gene signature previously associated to stimulatory dendritic cells abundance and better outcome in Melanoma patient are slightly enriched in archetypes with high total immune infiltration (Barry et al., 2018) (Figure S6A and S6C). Altogether, those data suggest that while the 12 immune archetypes identified may represent the dominant combination of cell types and transcriptomic profiles present in tumors, they may independently be populated with other cell types or signatures.

### 10-Features Tumor Archetype Ties Closely to Tumor Biology And Disease Outcome

Finally, we sought to examine the relationship of these immune archetypes to the phenotype of the tumor cells themselves. As previously shown for the coarse archetypes, the 10-feature archetypes are also diverse in their tissue of origin; however, some tumor indications remain highly represented within particular archetypes (e.g., Kidney: Immune Rich CD8, Melanoma: T cell centric archetypes) (Figure 7A and Figure S7). Regardless of the frequent intra-indication heterogeneity, we observed a strong relationship between tumor proliferative capacity, measured by the fraction of tumor cells which were Ki67^+^ as measured by flow, and immune desert and myeloid centric archetypes (Figure 7B).

**Figure 7:**
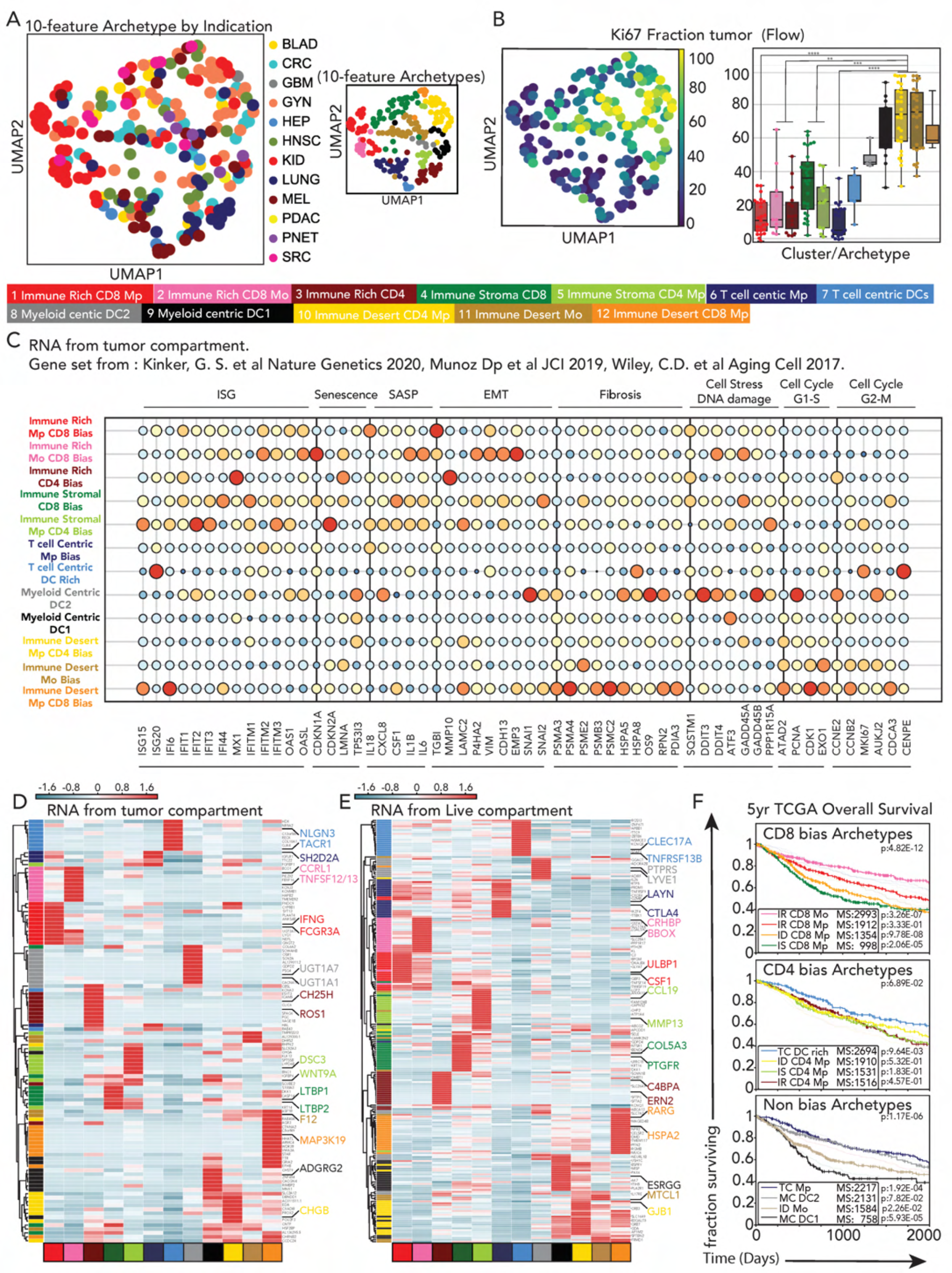
Immune archetypes tie closely to tumor biology and disease outcome. A. UMAP display and graph-based clustering of tumor immune archetypes using Tcell, Myeloid, CD90+ CD44+ Stroma, CD4, CD8, Macrophages, Monocytes, cDC1 and cDC2 feature scores color-coded by tumor indication. Each dot represents a single patient summarized by the 10 features. B. UMAP overlay (Left) and box and whisker plot (Right) of tumor proliferation measured by frequency of Ki67+ CD45− cells by flow cytometry in the IPI cohort. C. Bubble plot of the median gene expression, in the tumor compartment, of gene sets associated with previously identified tumor transcriptional programs grouped by cluster/archetypes in the IPI cohort using 10 features. D,E. Heatmap and hierarchical clustering of immune archetype gene signatures median expression in the tumor compartment (D) and the live compartment (E) in the IPI cohort. F. Kaplan-Meier overall survival curve from the TCGA cohort for each immune archetype using gene signatures in (E), split by T conventional subset enrichment). Median survival (MS) and p-value associated to each survival curve are noted.

Based on this we assembled gene signatures for various aspects of tumor biology to determine whether immune features mapped to tumor biology. Expression of cell cycle associated genes indicated enhanced tumor cell proliferative capacity in ID archetypes, consistent with the Ki67 data (Figure 7C). Conversely, IR and ISR tumors were enriched in other transcriptomic programs such as ISG, Senescence or EMT (Kinker et al., 2020; Muñoz et al., 2019; Wiley et al., 2017) and only some immune deserts were highly enriched in fibrosis associated genes (Figure 7C).

Without pre-selection of genes from previous literature, we performed DGE and then hierarchical clustering using the most differently expressed genes, between archetypes on the sorted tumor RNAseq compartment (Figure 7D). Within the 12 sets of differential genes, we noted some discernable patterns, for example increased IFNG expression in tumor cells from the IR CD8 Macrophage archetype, concordant with increased type 1 response in immune cells from the same archetype.

In order to define gene signatures that we could assess in TCGA, we queried our sorted live compartment to find genes that correlated with each archetype (Figure 7E). This identified a set of 12 unique gene signatures from which we observed both immune (e.g. LAYN, CTLA4, CSF1, CCL19) and non-immune (CRHBP, PTGFR, HSPA2, MTCL1) related genes. Using these minimal 12 signatures on the TCGA dataset, we demonstrated that the tumor archetypes identified in our IPI dataset were retrieved with a similar relative composition of archetypes by indications in KIRC, CRC, LUAD, HNSC, BLCA or with a slight shift between archetypes from the same coarse archetype (i.e. : ID CD4 Bias to IR CD8 bias) in GYN and SKCM (Figure S7). This suggests the predictive potential of these signatures for rapid classification of patients into archetypes.

Finally, survival analysis of the different tumor archetypes identified in the TCGA dataset for each cancer type separately showed that better outcome may be indication-specific and each indication may have a different type of TIME promoting the best immune response (Figure S7). However, when survival analysis was performed across indications, we detected significant outcome differences between archetypes that have similar T cell subset enrichment (Figure 7F). For instance, in IR CD8 archetypes the apparent enrichment in monocytes over macrophages (Pink archetype versus Red) is associated to better survival (Median survival:2993 days) and more so when compared to ID and IS CD8 biased archetypes (p-value 4.82E-12) (Figure 7F top). Conversely, in archetypes with no significant bias between CD4 and CD8 T cell, TC archetypes characterized by an enrichment of Macrophages over monocytes displayed the higher median survival (22217 days, p-value 1.17E-6) while in archetypes with CD4 T cell bias no significant outcome difference between archetypes was detected regardless of myeloid biases (p-value 6.89E-2) (Figure 7F middle and bottom). This indicates that tumor archetypes provide a template to study different subtypes of anti-tumor immune response in a variety of primary human cancer and to understand how to best modulate these archetypes depending on specific immune context.

## DISCUSSION

In this study, we present a holistic survey of dominant immune archetypes across 12 cancer indications using fresh surgical tumor specimen from 364 patients. Empowered by complementary profiling assays that assembled corresponding information from matched compositional, transcriptomic and imaging data on a variety of tumor-infiltrating cells (Figure 1, http://immunoprofiler.org/), we were able to discover 12 distinct immune archetypes that span cancer types. Each archetype is made up of a unique combination of cell composition— some used to derive the cluster, but many that are learned from it—and immune and tumoral transcriptomic phenotypes.

Starting with just 10 independent cell compositional features, unsupervised clustering revealed only 12 distinct clusters. While more work will need to be done to ensure that this is not due to a lack of sampling, this implies that some combinations of cell densities may not exist in the TIME of solid tumors. This is partly in line with previous work which identified six immune subtypes spanning multiple tumor types in TCGA using mainly immune cell fractions from deconvolution analysis of all tissue bulk RNA-seq data and immune gene expression signatures (Thorsson et al., 2018). Notably, similar dominant immune pathways are also revealed in our analysis such as the IFN pathway or the imbalance between T cell and MPC subsets. However, our analysis leveraged our linked compositional data via flow cytometry and RNAseq data from sorted cell populations to identify further the origin of those different immune networks which may partly explain our higher number of archetypes. Future work using single-cell omics across tumor types will be critical to determine if the 12 immune archetypes identified in our study can be further subdivided or if cell subsets identified using these technologies generally align with our current landscape (Cheng et al., 2021; Slyper et al., 2020; Zhang et al., 2020).

As previously shown by others, in a single cancer type or in tumor mouse models, the TME can be coarsely categorized into immune rich and immune desert areas (Duan et al., 2020; Galon and Bruni, 2019; Mariathasan et al., 2018). Our analysis revealed that the tumor immune archetypes subdivide this broad classification and identify distinct immune networks for each archetype and with unique relationships amongst cell densities and chemokine networks. While IR archetypes are characterized by patterns previously described in Melanoma, Head and Neck or Breast malignancies—such as a correlation between CD8+ T cells, cDC1 ratio and NK cell abundance (Barry et al., 2018; Salmon et al., 2016) or between CD4+ T cell and cDC2(Binnewies et al., 2019; Michea et al., 2018)—other archetypes implicate different immune cell networks such as the apparent enrichment of plasma cells and Tregs in TC archetype, a pattern previously identified as supporting plasma cell residency in bone marrow (Glatman Zaretsky et al., 2017). It is tempting to propose that conserved relationships of cell densities found in such a non-cancerous setting might be because some tumors hijack a particular immune archetype that may normally be used in a completely different setting. This idea is perhaps reinforced by the distinct chemokine expression profile identified for each tumor immune archetype. While IR CD8 archetypes are defined by a CXCL9, CXCL10, CXCL11/CXCR3 axis already described in the TME, ID CD8 Macrophage bias archetype is defined by a strong expression of chemokines binding CX3CR1, an axis essential in immune surveillance and homeostasis (Gerlach et al., 2016).

The association between chemokine expression programs and the tumor immune archetypes also suggests an association with a specific spatial organization. While we confirmed the overall correspondence between flow cytometry, transcriptomic data and immunohistochemistry assays, the spatial landscape of the tumor immune archetypes would need further investigation. In future, the combination of multiplexed imaging technologies such ion beam imaging (MIBI) (Angelo et al., 2014; Keren et al., 2018) and single-cell spatial transcriptomics (Asp et al., 2019; Hu et al., 2020) will enable this analysis, using the knowledge that only some archetypes have the relationship to begin with. In this way, archetype-discovery plays an important role in selecting tissues containing similar biology for concerted study.

Tumor-specific genetics and mutational burden have been proposed to be key for anti-tumor immunity in multiple indications (Brown et al., 2014; Ghosh et al., 2021; Goodman et al., 2017). However, further investigation on extrapolating immune gene signature across TCGA cohort showed no correlation between gene expression and mutational burden in any cancer type (Spranger et al., 2016). Conversely, we demonstrated that immune tumor archetypes are associated to tumor proliferation, diverse tumor transcriptomic programs and overall survival. Studying the synergy between immune archetypes and tumor mutational profile, may elucidate further the importance of tumor mutational burden in anti-tumor immune response (Quigley et al., 2018).

In summary, our comprehensive characterization of the TME across many human solid cancer types reveals the existence of common and reproducible immune archetypes defined by distinct cell networks. To facilitate usage of our data for the wide research community, we will provide access to transcriptomic and compositional data for each archetype (datalibrary.ucsf.edu/public-resources). It is still unclear whether improving tumor cure rates will be based on the enhancement of a specific archetype, and the expansion of this classification in metastatic tumors as well as in animal model may help to test this (Maynard et al., 2020). Moreover, it will be important to expand this analysis to the peripheral immune system as well as biopsies before and after immunotherapies in order to define the relationship between ‘dominant’ and ‘reactive’ tumor immune archetype across tumor types (Bi et al., 2021; Sade-Feldman et al., 2018). Furthermore, there will undoubtedly be many additional ways to explore this dataset and discover patterns of immune and cancer biology. In this sense, the IPI dataset and the accompanying curation by these archetypal assignments can now serve as a rich resource to gain deeper understanding of cancer immunity in patients and therefore serve as a framework to direct immunotherapies to the most relevant biology.

## ACKNOWLEDGEMENTS

We thank all members of the Krummel Lab, UCSF ImmunoX, UCSF CoLabs and the UCSF Immunoprofiler Consortium for discussion and guidance while developing this study. We would like to thank Dr Nicholas Kuhn for scientific discussion. We would like to thank Dr Kenneth Hu for editing the manuscript. We would like to particularly thank Isabelle Tingin, Garry Shumakher, Elizabeth Edmiston and Meghan Zubradt for their constant support and help during this study. We would like to thank Dr Ana Catharina Silva for her help in designing the graphical abstract. Acquisition and analysis of certain human samples described in this study was partially funded by contributions from AbbVie, Amgen, Bristol-Myers Squibb, and Pfizer as part of the UCSF Immunoprofiler Initiative. Further support came from the NIH (R01 U01) for MFK and P30 : P30CA082103 for ABO. Finally, we thank all patients and their families, for placing their trust in us.

## AUTHOR CONTRIBUTIONS

AJC, BS, AB and MFK performed and/or provided supervision of experiments, generated and analyzed data, and contributed to the manuscript by providing figures, tables and important intellectual contributions. AJC, BS and MFK had full access to all of the data in the study and take responsibility for the integrity of the data and the accuracy of the analyses. AJC, BS and MFK wrote and edited the manuscript. AJC, BS, SA, AAR, DA, performed computational analyses of the data. AJB and MA performed the library RNA extraction and libraries construction for all the bulk RNA sequencing present in this manuscript. GKF, ABO and SA provided important intellectual input in genomics and statistical analysis for this study. AJC, JT, NWC, PY, GCR, DK, AS, BD, AB, KT, ES, PJ, JS, JH, JL, TC, KCB, RJA, TH, VC were part of the UCSF Immunoprofiler team who performed digestion, staining, sorting and initial quality control analysis on the different tumor specimens included in the IPI dataset. RKK, JSC, CEA, AID, PH, AAD, JRK, and EAC, are part of the UCSF Immunoprofiler clinical team who was actively involved facilitation the access to the tumor specimens included in the IPI dataset. AJC, VC, KCD, DJE, GKF, SA and MFK are leadership members of the UCSF Immunoprofiler consortium and were actively involved in the establishment of the pipeline and patient consent and the direction of projects. BS, AP, CC, JZ and KCD were actively participated in patient enrollment and clinical data collection. The UCSF Immunoprofiler initiative Consortium included all other scientist involved in the generation of data included in this study. All authors edited and critically revised the manuscript for important intellectual content and gave final approval for the version to be published.

## Supplementary Materials For

This PDF file includes:

Supplementary Figure Legends

Supplementary Tables 1 to 5

Extended STAR Methods and Associated References

### SUPPLEMENTARY FIGURE LEGENDS

**Figure S1:**
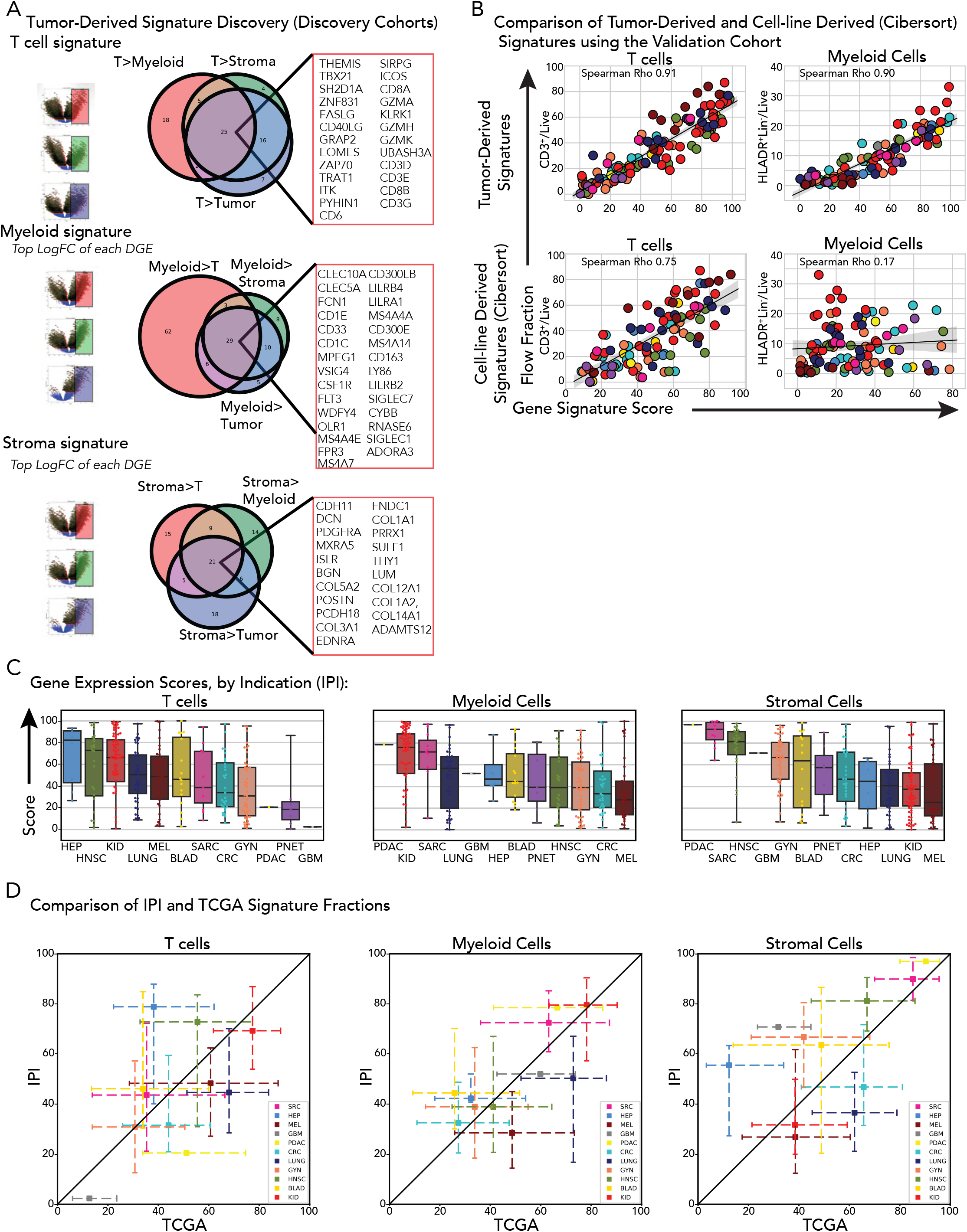
Generation and validation steps of T cells, Myeloid cells and Stromal cells features from solid tumors using flow cytometry and bulk RNA-sequencing, related to Figure 1. A. Volcano plots, Venn diagrams and gene names showing the method of feature gene signature discovery for the Tcell, Myeloid and CD90+ cd 44+ Stroma features using differential gene expression in tumor associated sorted compartments (see extended method section). B. Correlation plots of Tcell and Myeloid gene signature score against their corresponding flow population fraction (top) and CIBERSORT derived fraction (bottom) color-coded by tumor indication. C. Box and whisker plots of Tcell, Myeloid and CD90+ CD44 Stroma features gene scores in the IPI cohort. D. Cross-whisker plots comparing median Tcell, Myeloid and CD90+ CD44+ Stroma gene signature scores by tumor indication between the IPI and TCGA cohorts with interquartile range on both axes.

**Figure S2:**
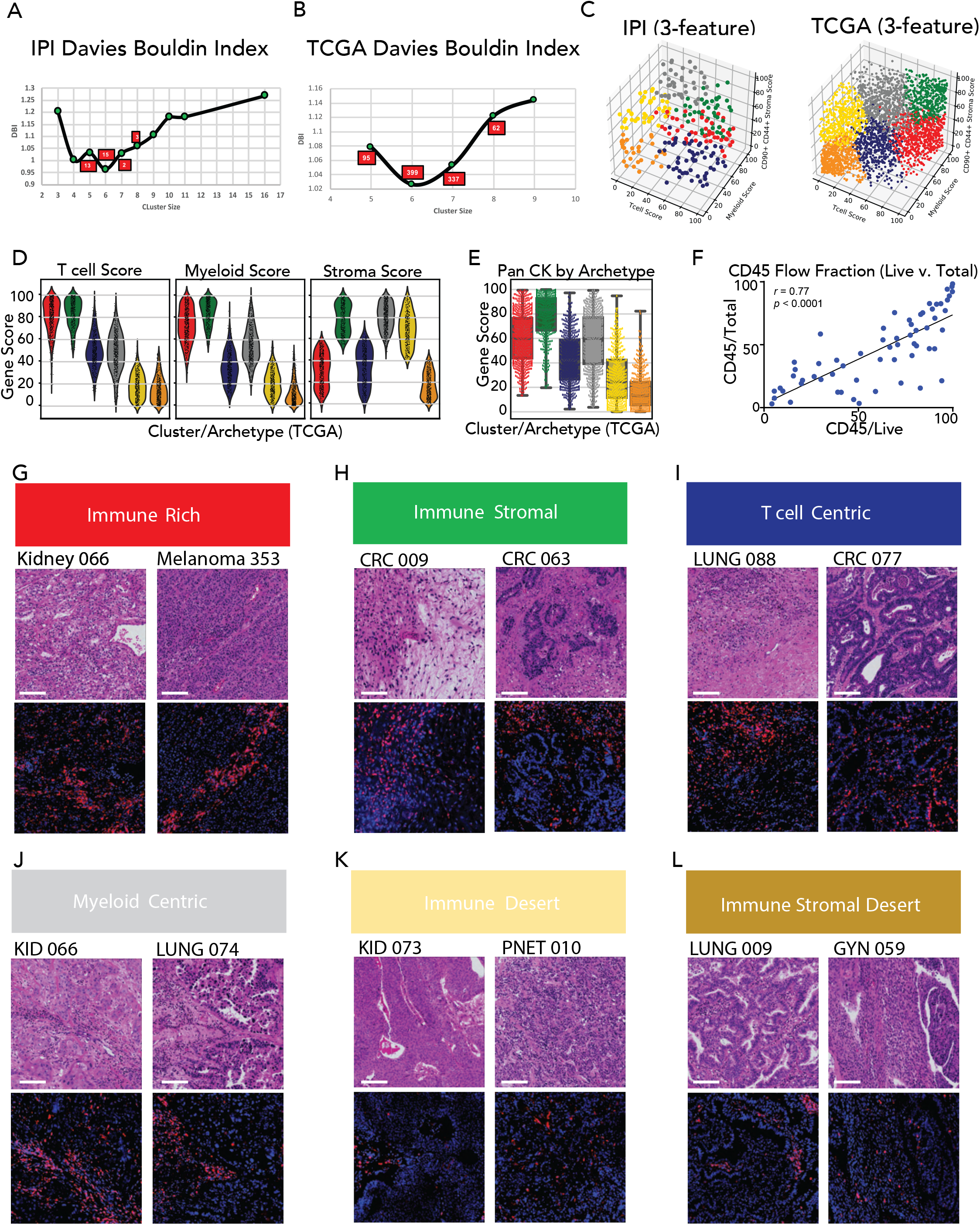
Identification of coarse immune archetypes in solid tumors using Louvain clustering on two independent datasets, related to Figure 2. A, B. Scatter plot of the Davies-Bouldin index and cluster size over multiple iterations of Louvain clustering and varying parameters using 3 features in the IPI (A) or TCGA (B) cohort. C. 3D plot of Tcell, Myeloid and CD90+ CD44+ Stroma scores color-coded by their cluster assignment from Louvain clustering these features in the IPI (left) and TCGA (right) cohorts. D. Violin plots of the Tcell, Myeloid and CD90+ CD44+ Stroma feature score for each cluster/archetype in TCGA cohort. E. Box and whisker plot of a pan chemokine gene score by cluster/archetype identified in TCGA cohort. F. Scatter plot of immune cell population fraction using only viable cells or total cells as denominator. G, H, I, J, K, L. Representative H&E (top) and immunofluorescence (bottom) images of tumor biopsies using CD45 (red) and DAPI (blue) staining for each cluster/archetype identified in IPI cohort.

**Figure S3:**
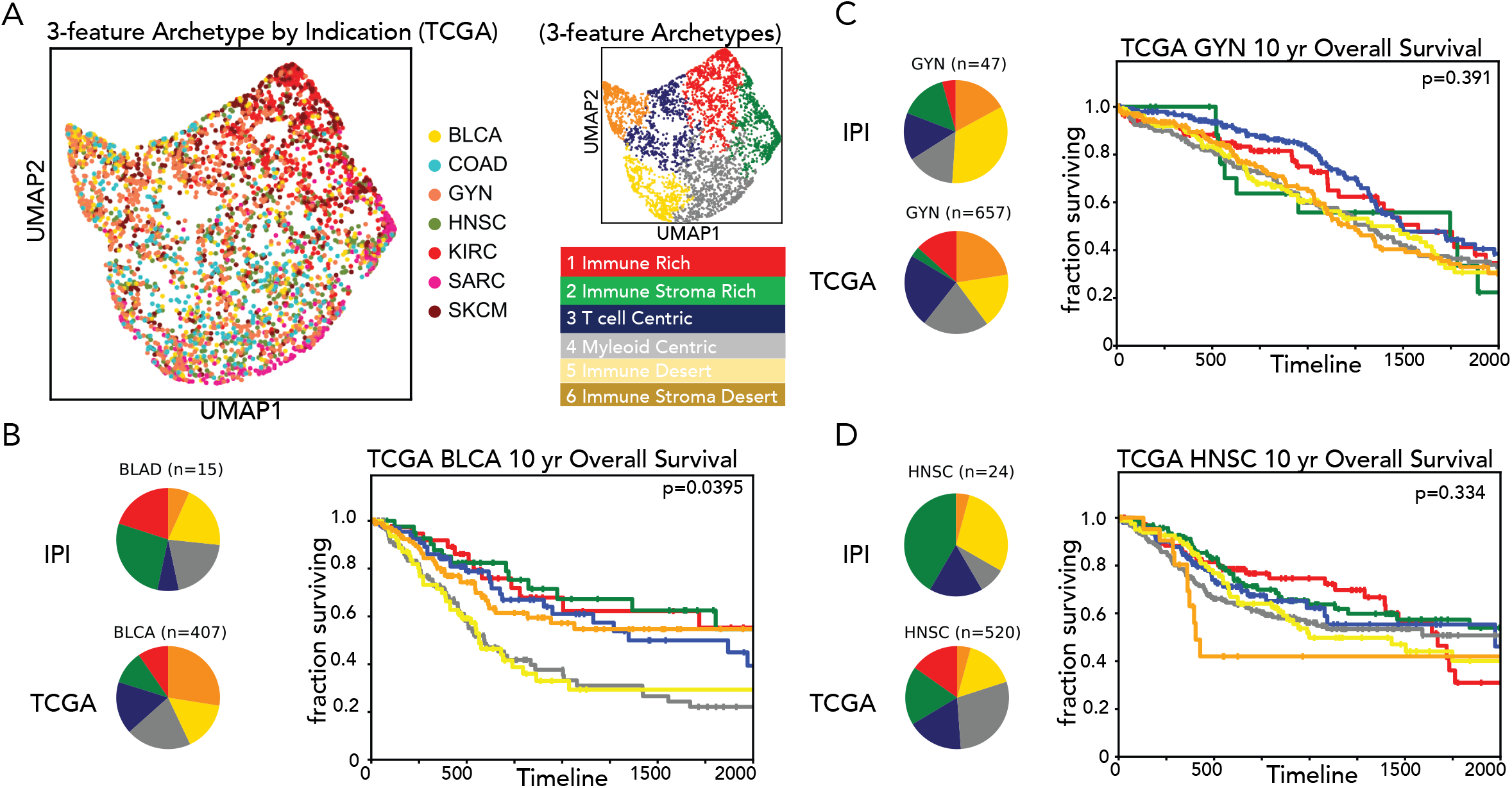
Coarse immune archetypes are independent of tissue origin and associated to overall survival., related to Figure 3. A. UMAP display and graph-based clustering of immune archetypes using Tcell, Myeloid and CD90+ CD44+ Stroma features to cluster patient with cohorts color coded by tumor type. B, C, D. (Left) Pie charts representing distribution of each archetype by indication from IPI (top) and TCGA (bottom) cohorts using 3 features. (Right) Kaplan-Meier overall survival curve for each immune archetype identified on TCGA dataset for Bladder urothelial carcinoma (B, BLCA), Gynecologic tumors (C,UCS + UCEC +OV),) and Head and Neck squamous cell carcinoma (D, HNSC).

**Figure S4:**
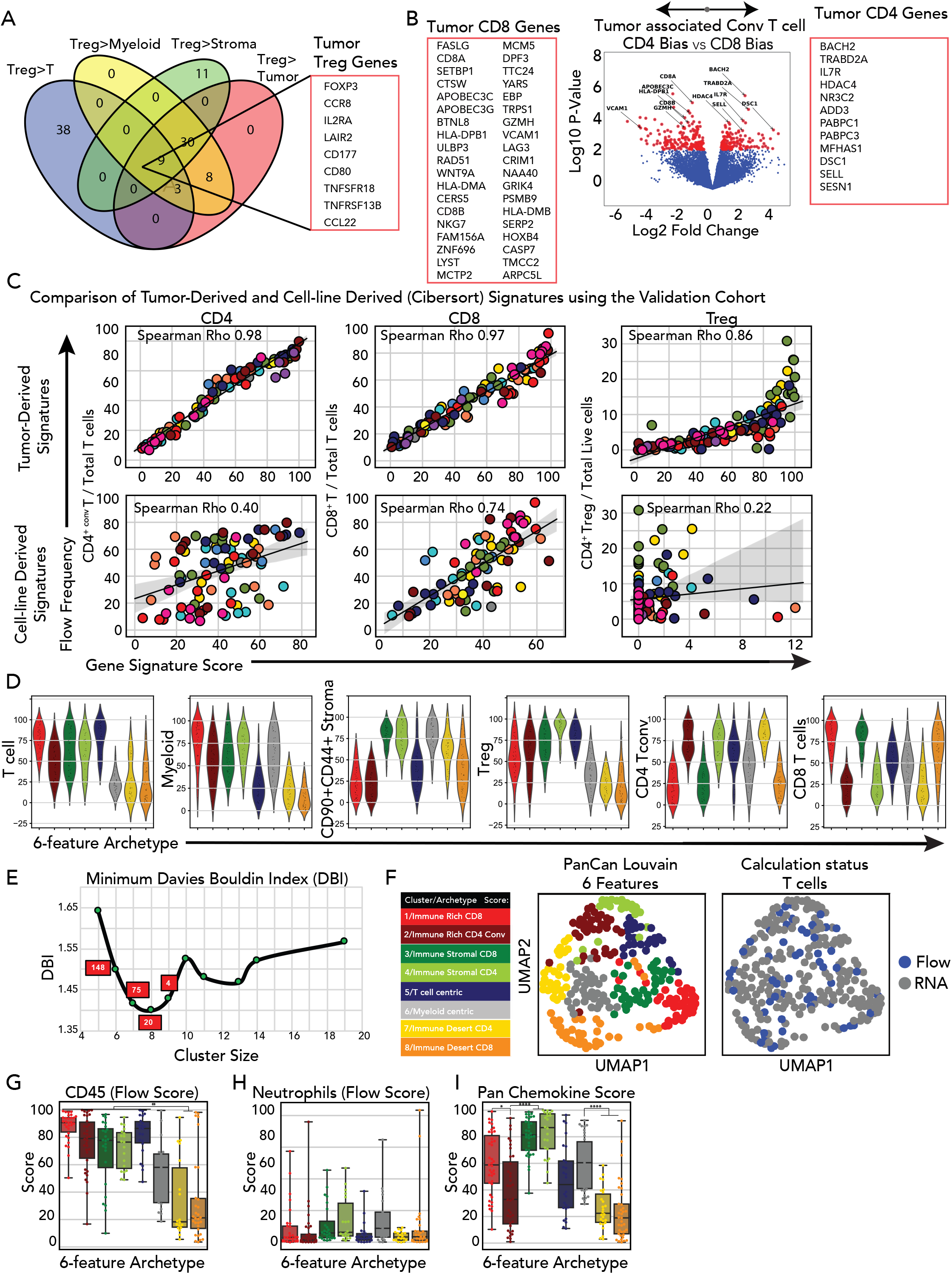
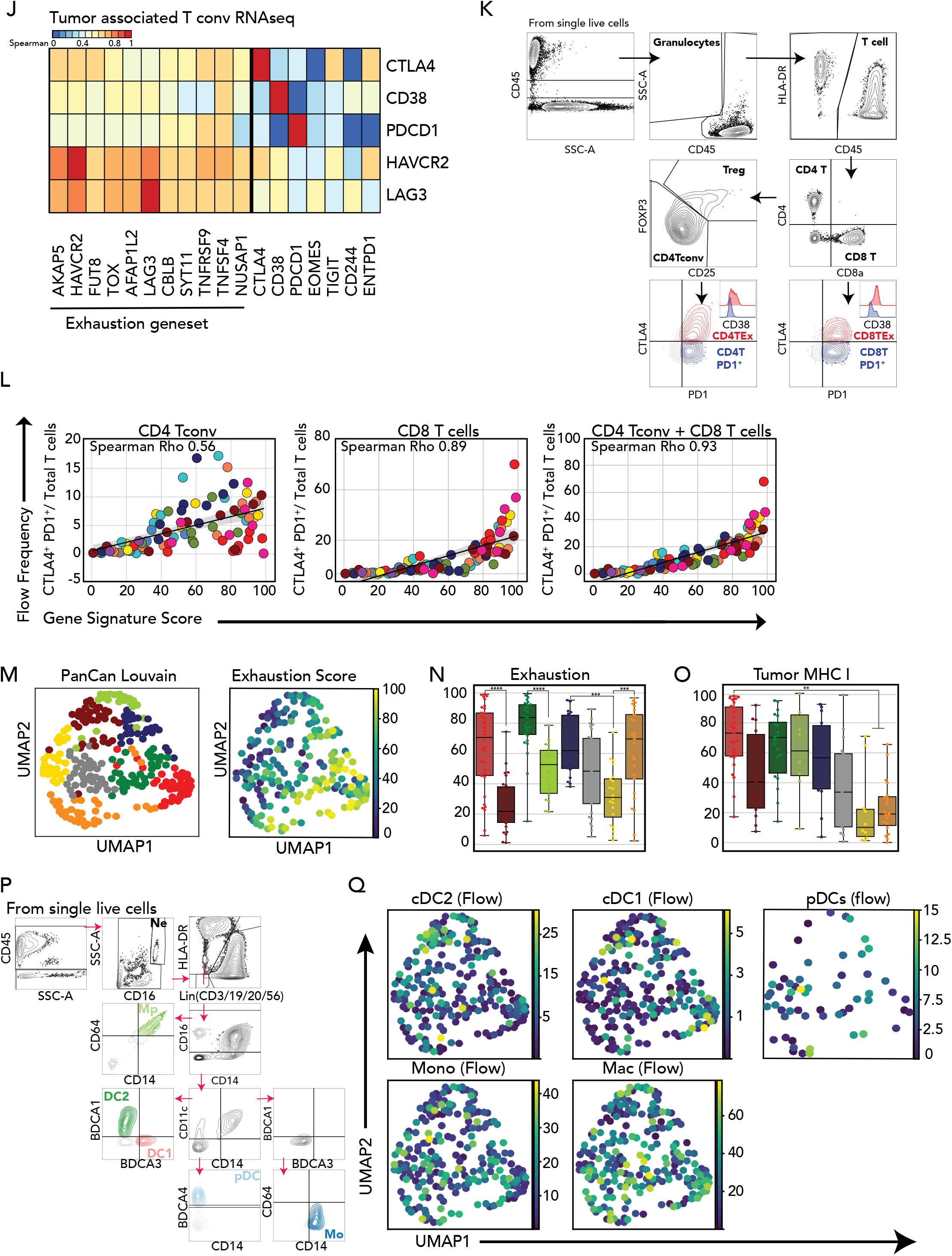
Generation and validation of CD4 CD8 and Treg signatures and further exploration of 6 feature archetypes, related to Figure 4. A. Venn diagram and gene names of the Treg (CD4+ regulatory T cell) feature gene signature score (see extended method section). B. Volcano plot visualizing differential gene expression between CD4+ or CD8+ T cells in the tcell compartment (see extended method section). Feature gene signatures are listed in red squares. C. Correlation plots of CD4, CD8 and Treg feature score versus their corresponding flow population fractions (top) and CIBERSORT derived fractions (bottom) color-coded by tumor indication. D. Violin plots of the 6 features distributions in each cluster/archetype in the IPI cohort. E. Scatter plot of the Davies-Bouldin index and cluster size over multiple iterations of Louvain clustering and varying parameters using 6 features in the IPI cohort. F UMAP of immune archetypes using Tcell, Myeloid, CD90+ CD44+ Stroma, CD4. CD8 and Treg features in the IPI cohort using KNN and Louvain clustering. (Right) UMAP overlay of samples that needed to calculate their CD4 and CD8 feature score flow cytometry population fractions (see extended method section).G, H. Box and whisker plot of total immune cell population fraction (G) and Neutrophils (H) using flow cytometry. I. Box and whisker plot pan chemokine gene score by cluster/archetypes in the IPI cohort using 6 features. J. Heatmap of the Spearman correlation coefficient of gene expression between CTLA4, CD38, PDCD1, HAVCR2 and LAG3 and our Exhaustion phenotype gene signature with additional genes associated with exhaustion used as a control. K. Gating strategy of tumor associated conventional CD4+ and CD8+ T cells and the markers used to define exhaustion. L. Correlation plots of individual and combined CD4+ PD1+ CTLA4+ and CD8+ PD1+ CTLA4+ conventional T cells population fractions against the Exhaustion phenotype gene signature score color-coded by tumor indication. M. (Left) UMAP display and graph-based clustering of tumor immune archetypes using Tcell, Myeloid, CD90+ CD44+ Stroma, CD4, CD8 and Treg features to cluster patients in the IPI cohort. (Right) UMAP overlay of the Exhaustion phenotype score (see J and extended method) calculated in the tcell compartment. N, O. Box and whisker plots of Exhaustion phenotype score (N) and MHC Class I phenotype score calculated in the tumor compartment (O) for each cluster/archetype identified in IPI cohort using 6 features. P. UMAP overlay of classical dendritic cell type 1 (cDC1) and 2 (cDC2), plasmacytoid dendritic cells (pDCs), monocytes (Mono) and macrophages (Mac) frequencies measured by flow cytometry.

**Figure S5.**
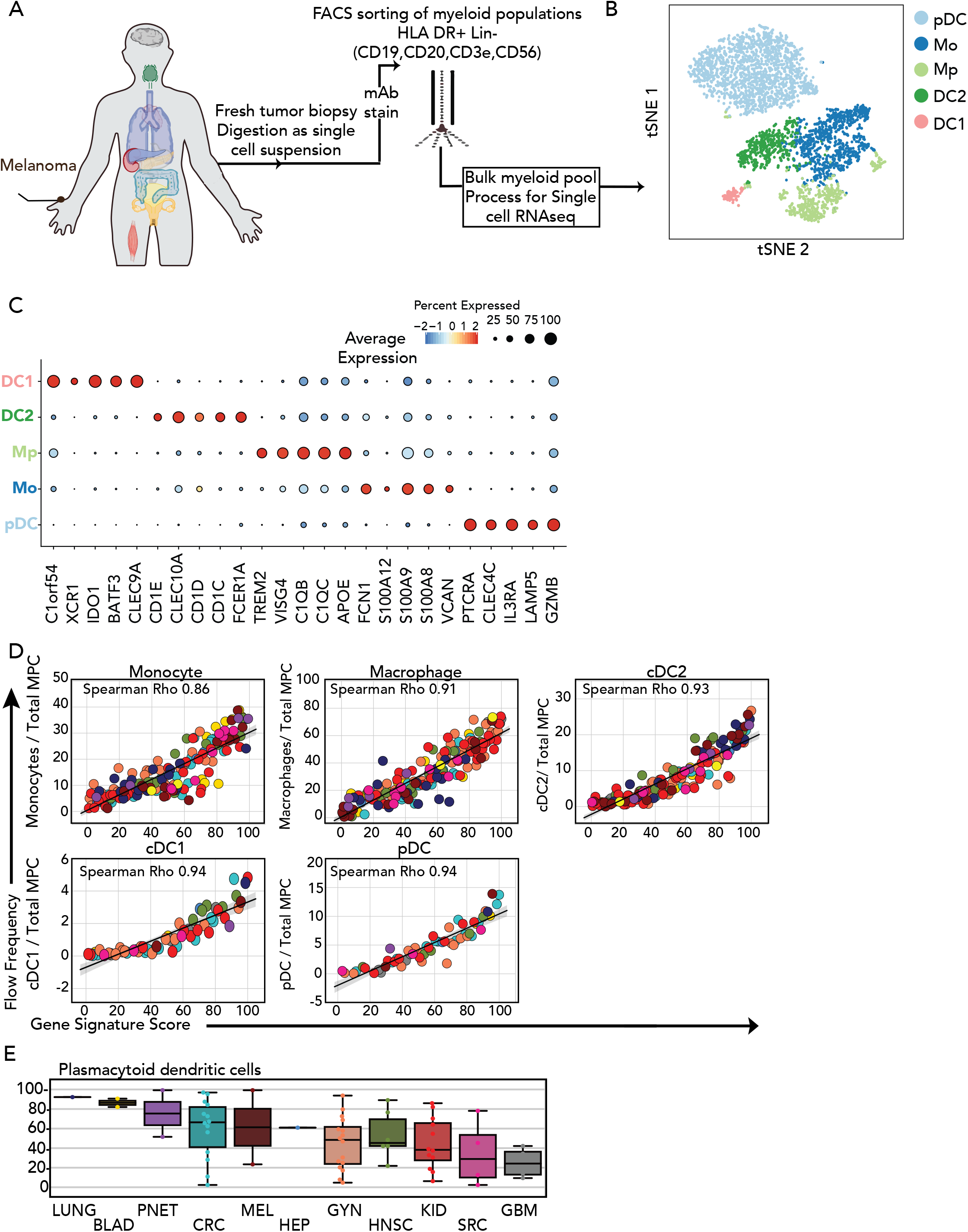
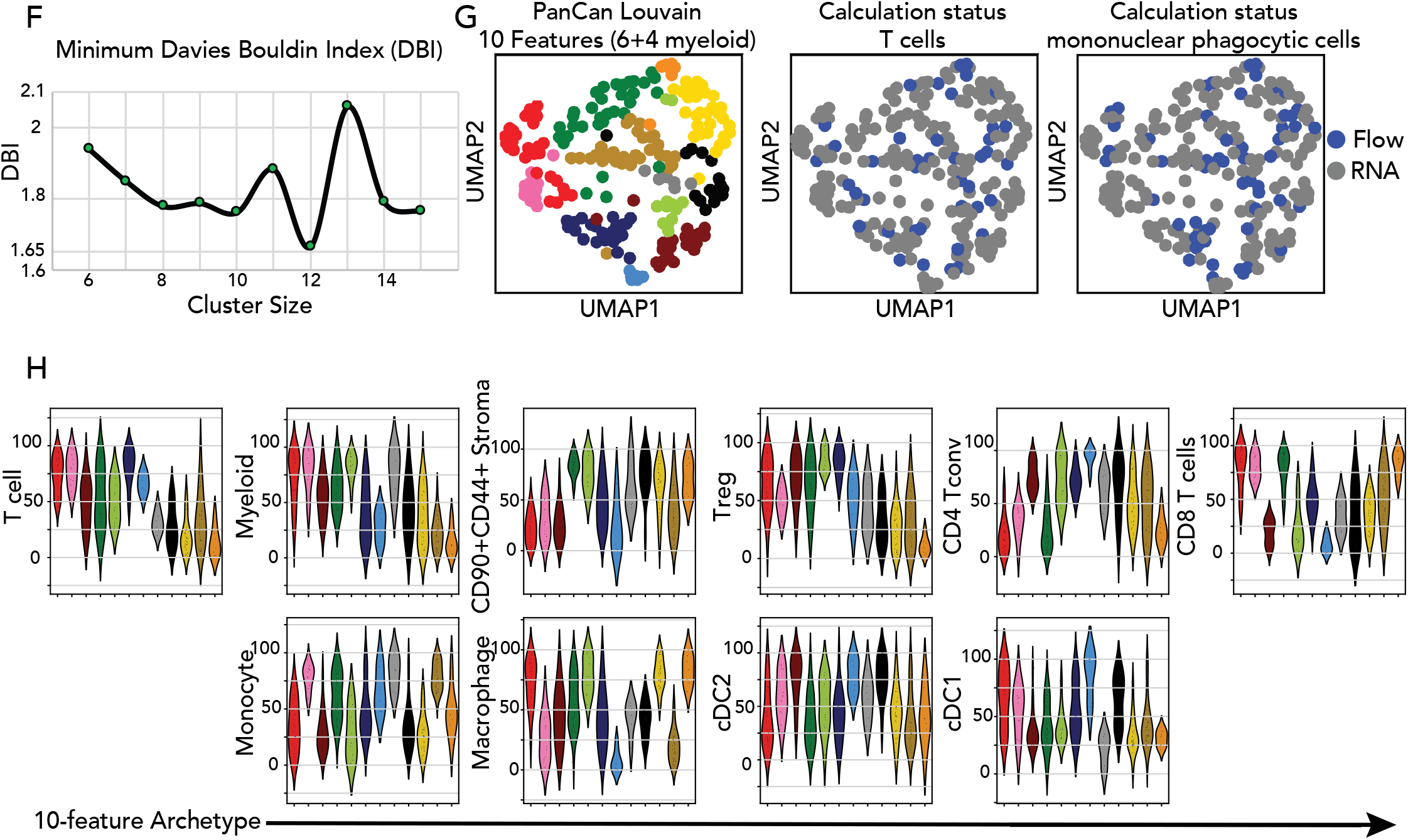
Single-cell RNA sequencing-derived myeloid gene signatures refine immune archetypes, related to Figure 5. A. Details of the processing pipeline for digesting fresh tumor biopsies into single cell suspension, submitting to multi-parametric flow cell sorting of tumor associated myeloid population and encapsulation for single-cell RNA sequencing. B. t-display and graph-based clustering of tumor associated myeloid cells from Melanoma processed for single-cell RNA sequencing. Each dot represents a cell. C. Dot plot of the top differentially-expressed-genes (DEG) between clusters identified in tumor associated mononuclear phagocytic cell (MPC) subsets. D. Gating strategy of tumor associated myeloid cells. E. Correlation plots of Macrophages, Monocytes, cDC1, cDC2 and pDC gene score against their corresponding flow population fractions in the myeloid compartment color-coded by tumor indication. F. Box and whisker plots gene score for pDCs (out of myeloid compartment) in IPI cohort. G. Scatter plot of the Davies-Bouldin index and cluster size over multiple iterations of Louvain clustering and varying parameters using 10 features in the IPI cohort. H. (Right) UMAP of tumor immune archetypes using Tcell, Myeloid, CD90+ CD44+ Stroma, CD4, CD8, Macrophages, Monocytes, cDC1 and cDC2 feature scores to cluster patients in the IPI cohort, using KNN and Louvain clustering. (Center-Left) UMAP overlay of samples that needed to calculate their CD4 and CD8 feature score and their Macrophages, Monocytes, cDC1 and cDC2 feature scores using flow cytometry population fractions (see extended method section). I. Violin plots of the 10 features distributions in each cluster/archetype in the IPI cohort.

**Figure S6:**
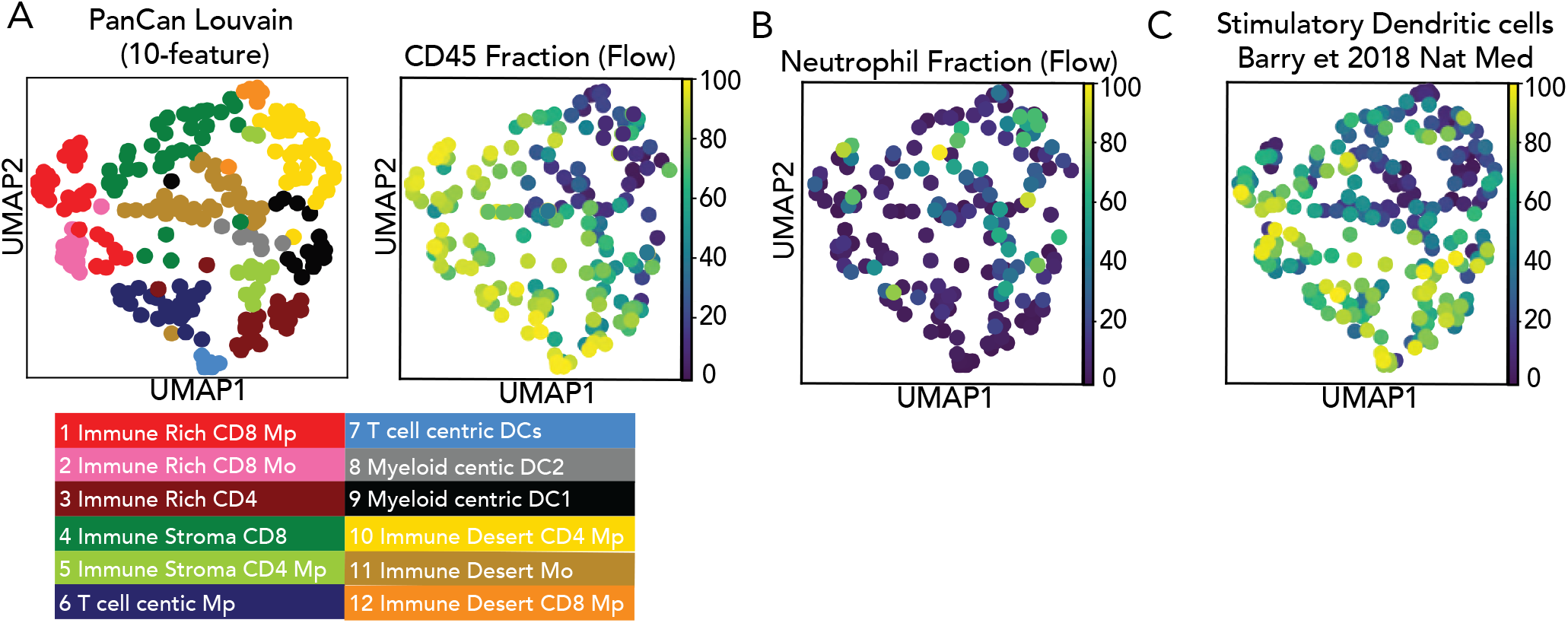
immune cell frequency and transcriptomic profile pattern associated to each tumor archetype, related to Figure 6. A. (Left) UMAP display and graph-based clustering of tumor immune archetypes using Tcell, Myeloid, CD90+ CD44+ Stroma, CD4, CD8, Macrophages, Monocytes, cDC1 and cDC2 feature scores to cluster patients in the IPI cohort. Each dot represents a single patient summarized by the 10 features. (Right) UMAP overlay of CD45+ cell frequency measured by flow cytometry. B. UMAP overlay of neutrophils frequency out the total immune cells measured by flow cytometry. C. UMAP overlay of a Stimulatory DC gene signature score measure on ‘all viable’ RNAseq compartment

**Figure S7:**
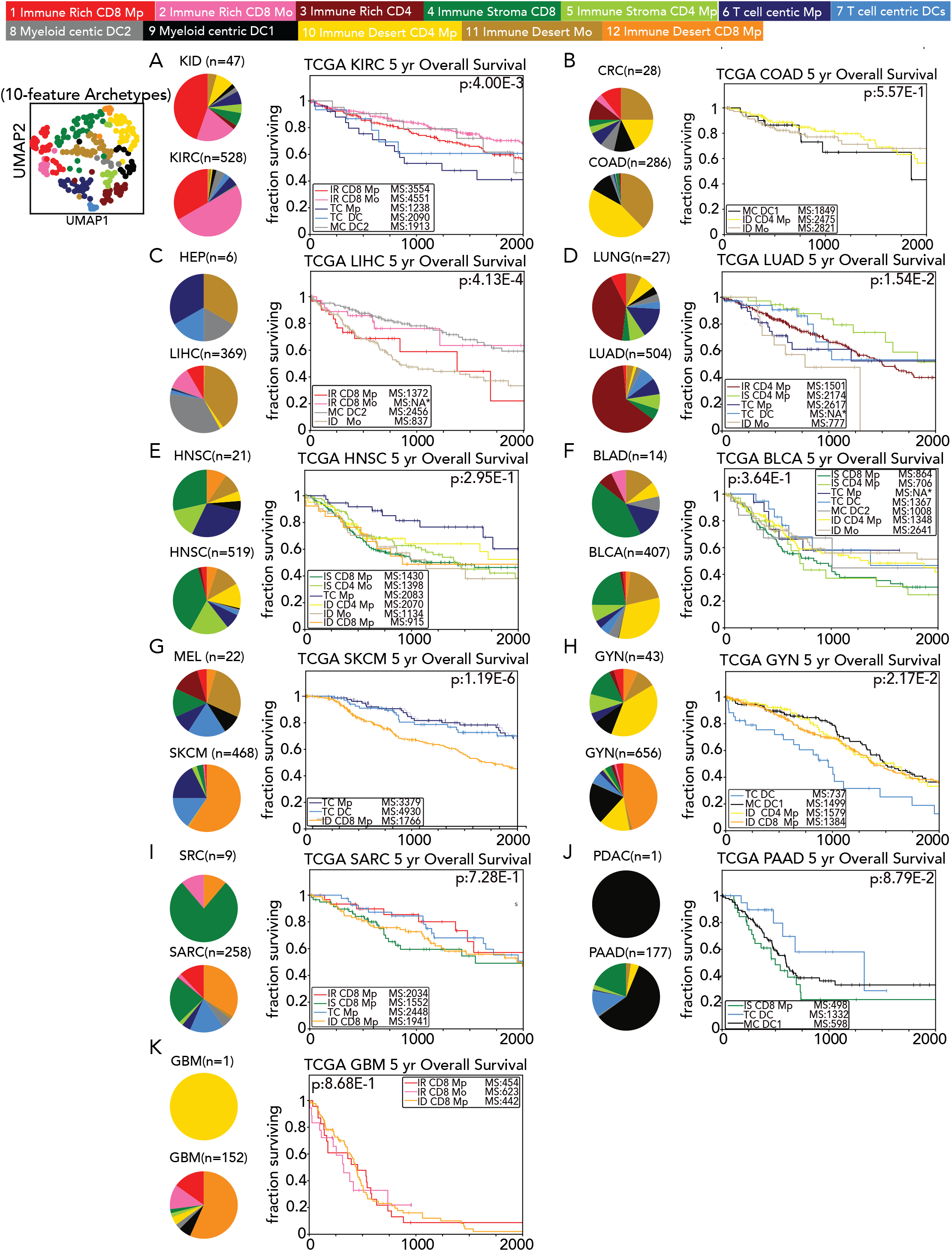
Immune archetypes tie closely to tumor biology and disease outcome, Related to Figure 7. A to J. (Left) Pie charts representing the distribution of each archetype by indication in IPI 10-feature clustering (top) and TCGA (bottom) cohorts. (Right) Kaplan-Meier overall survival curve for the most abundant immune tumor archetype by tumor indication in TCGA cohort (see Extended method).

**Supplementary Table 1:**
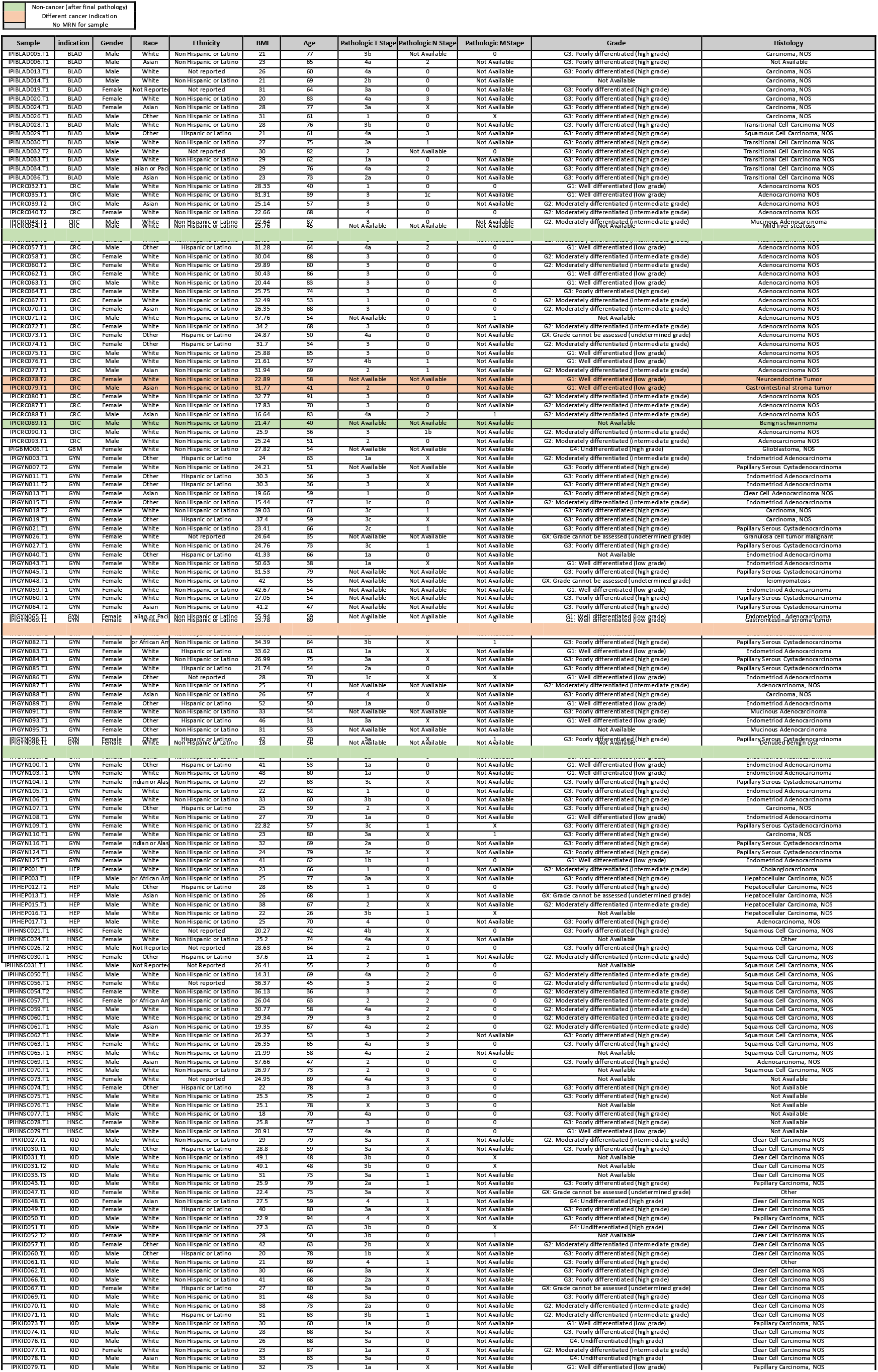
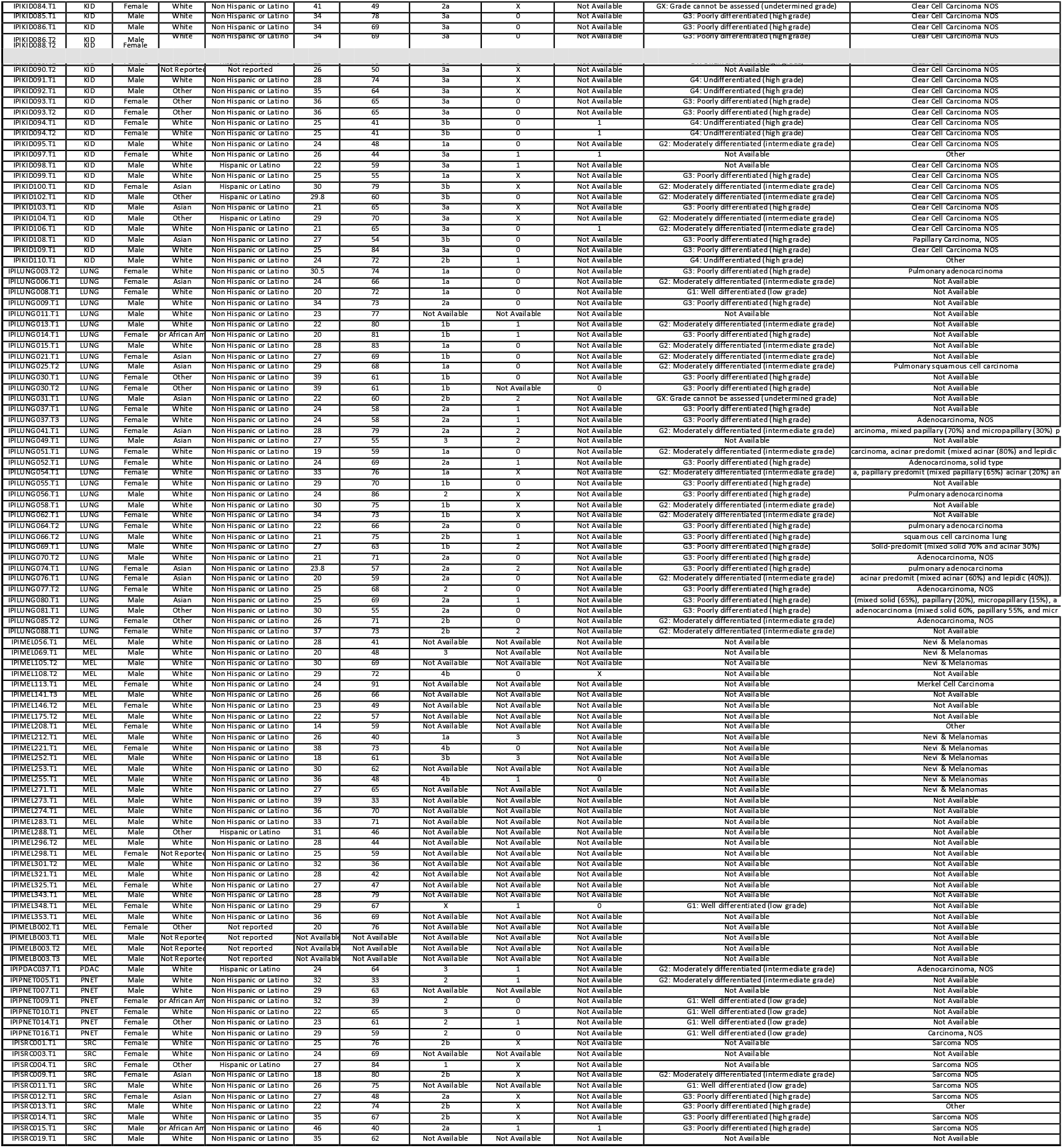
List of UCSF Immunoprofiler Initiative samples and associated clinical data. List of each tumor biopsy included in the first Louvain clustering Figure 2A and associated demographic and Clinical data.

**Supplementary Table 2:**
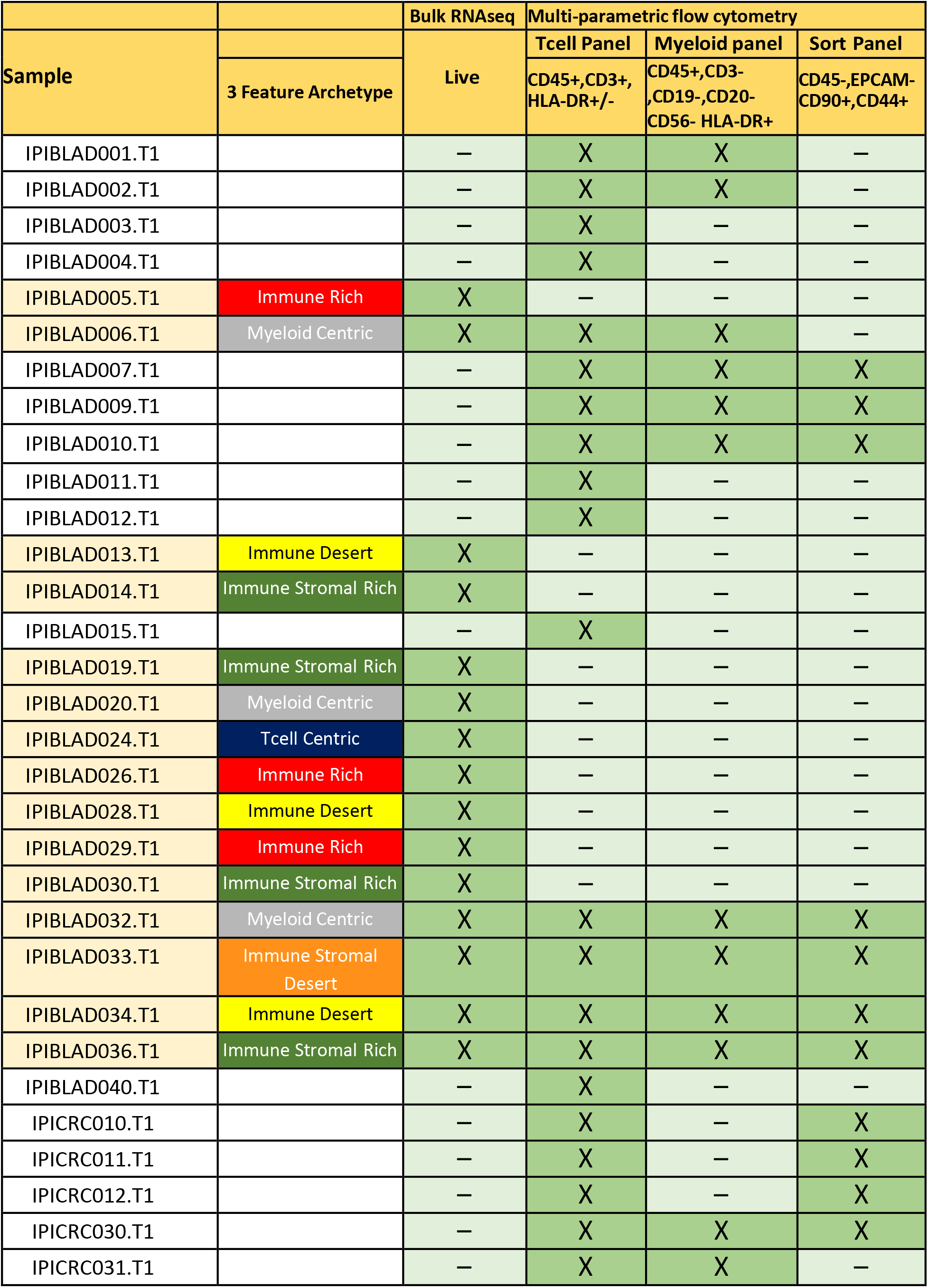
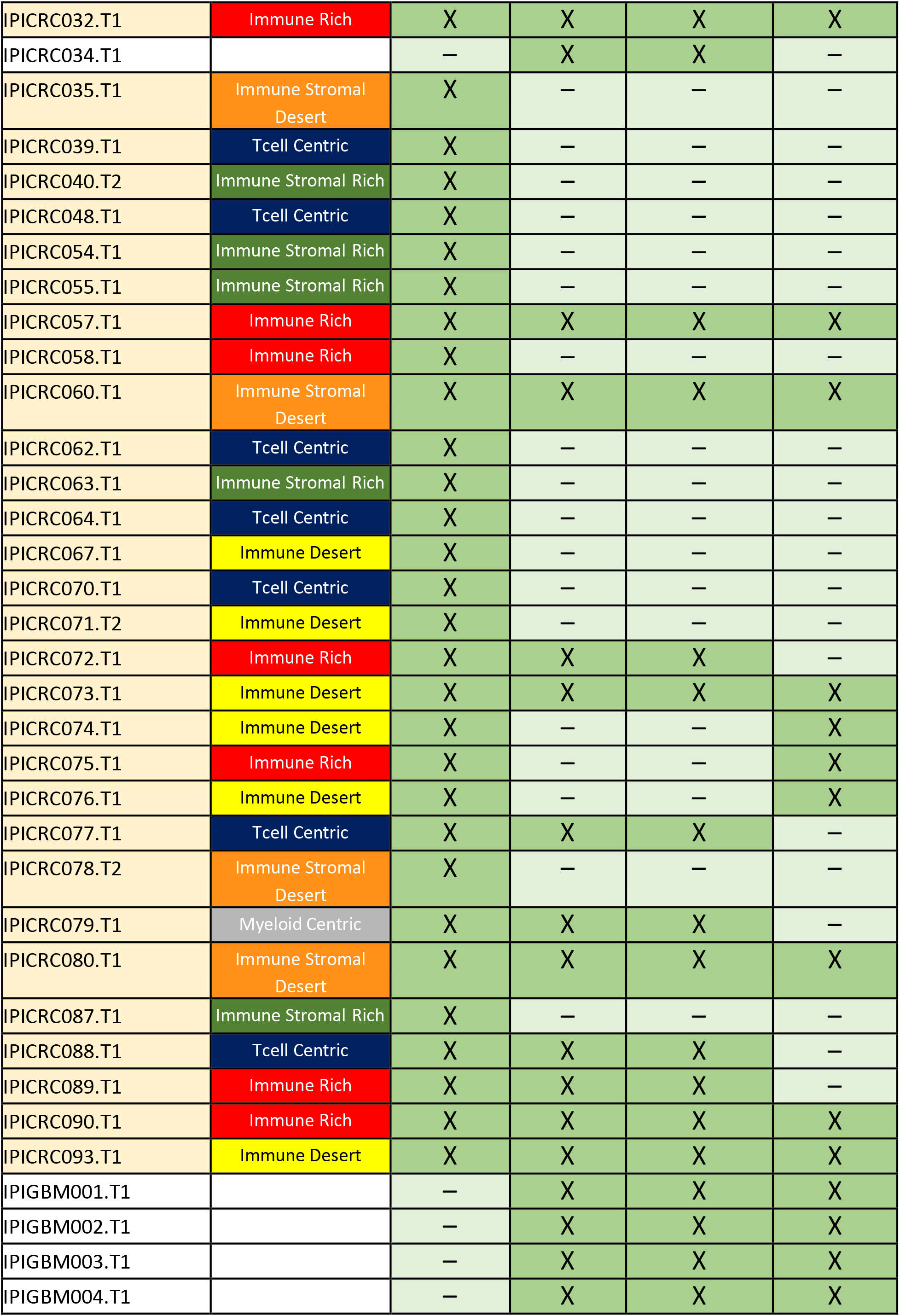
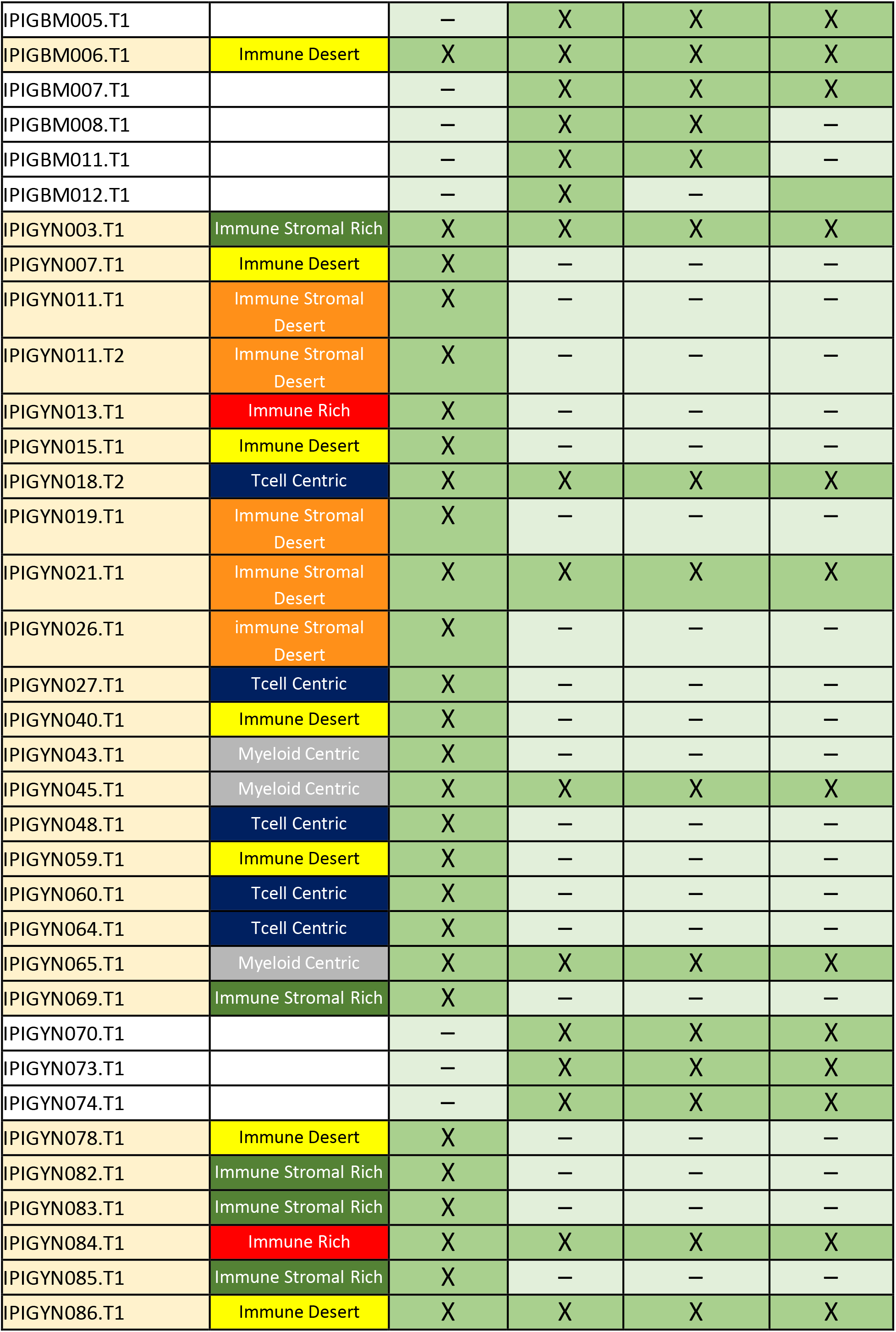
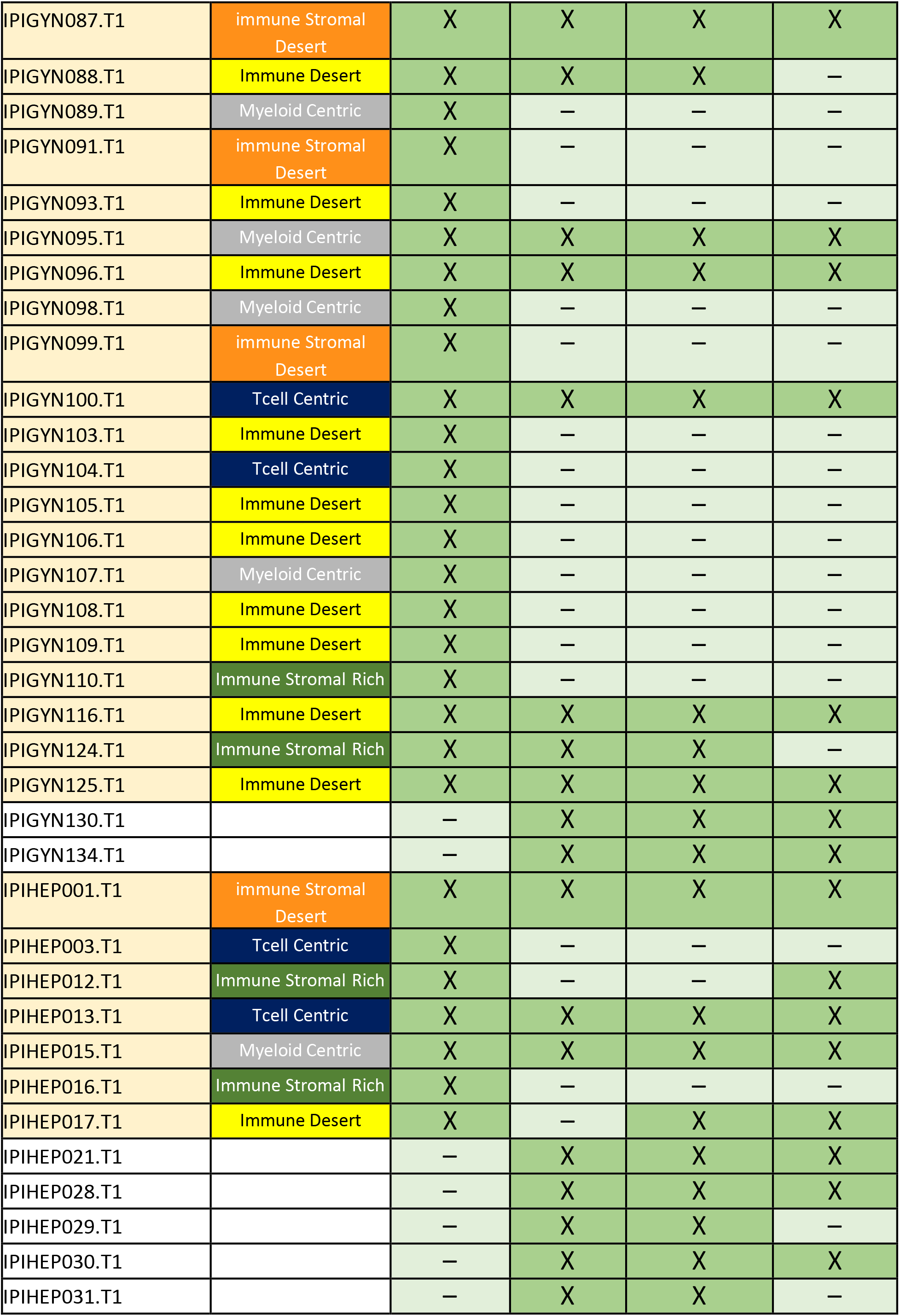
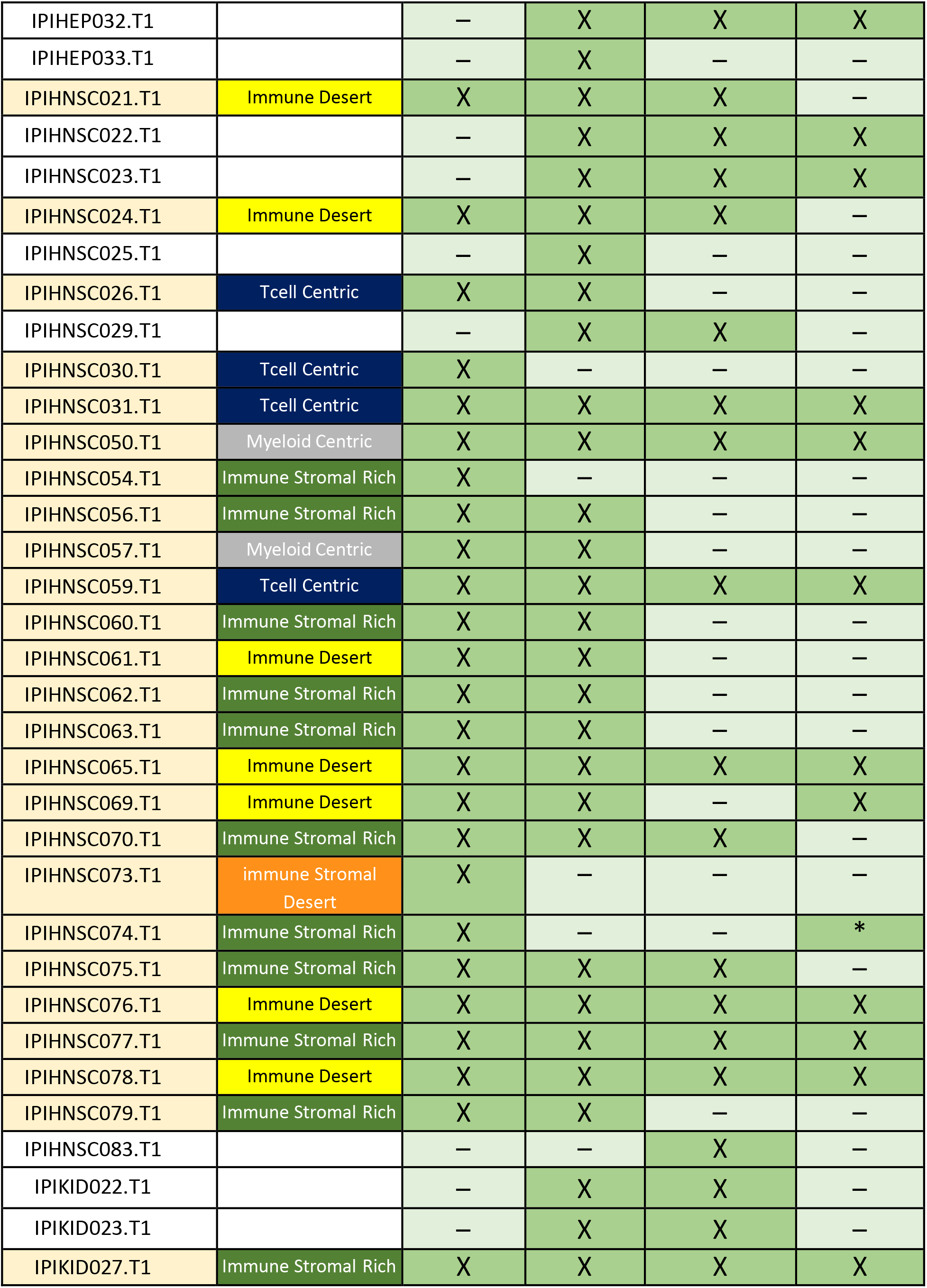
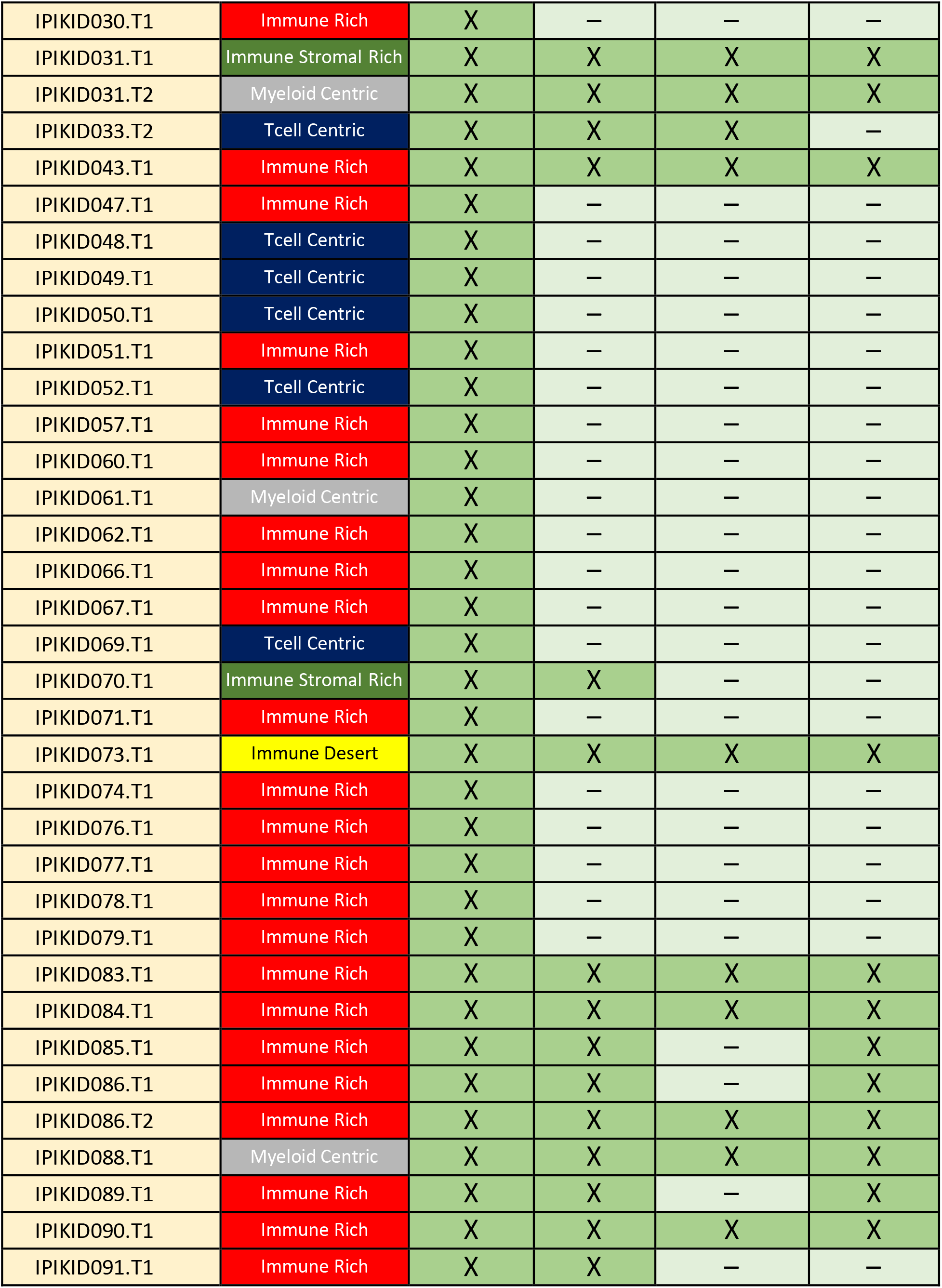
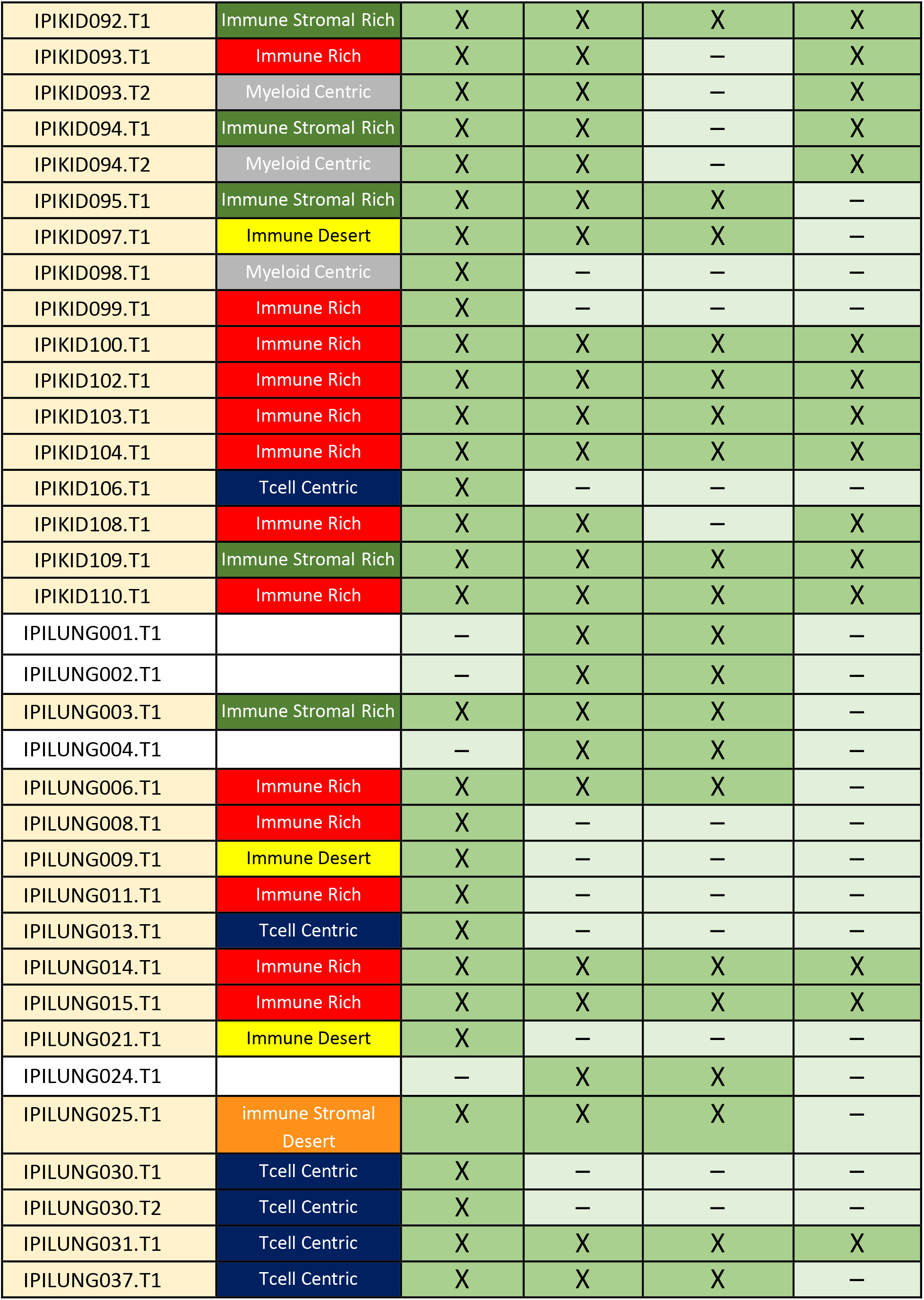
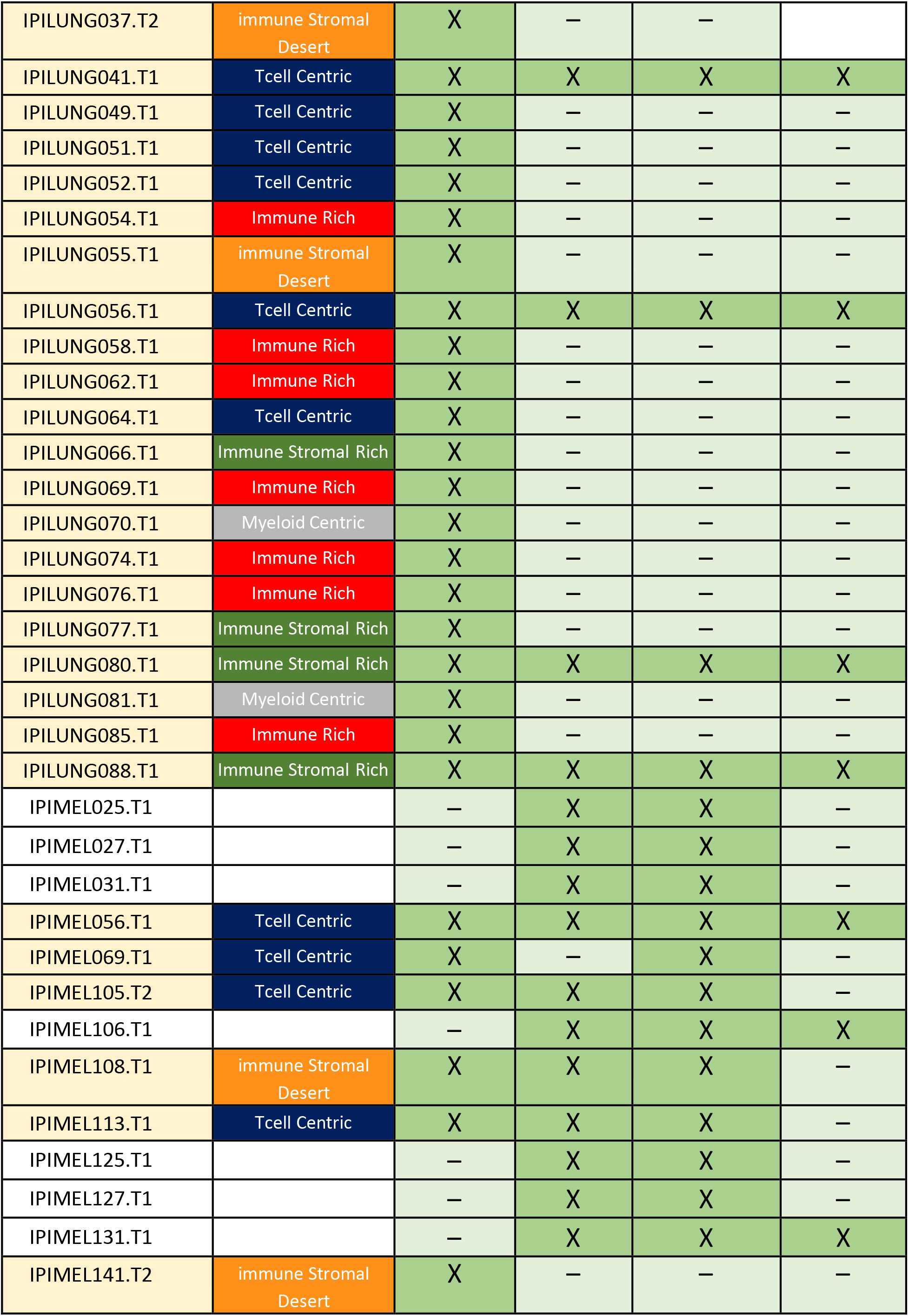
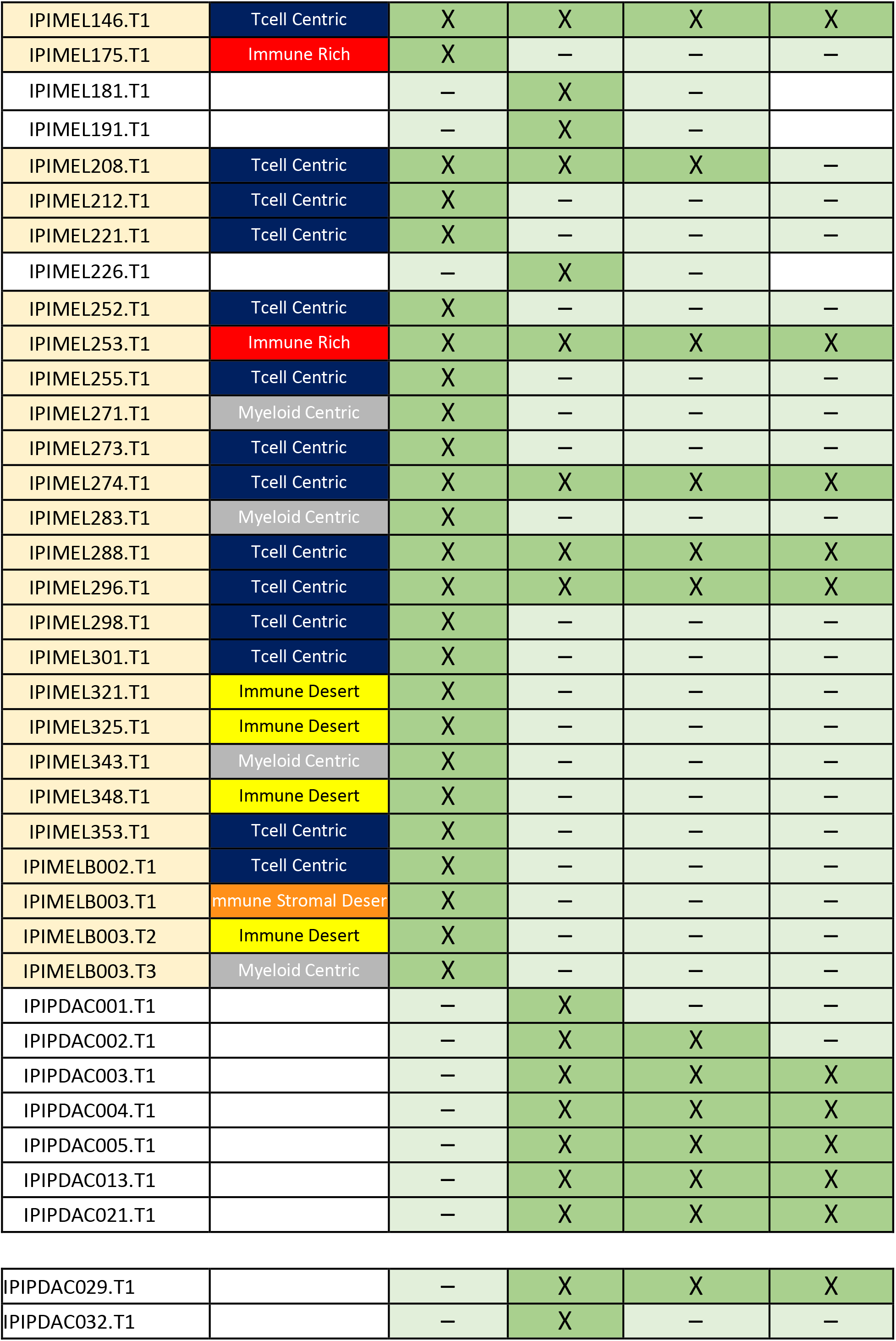
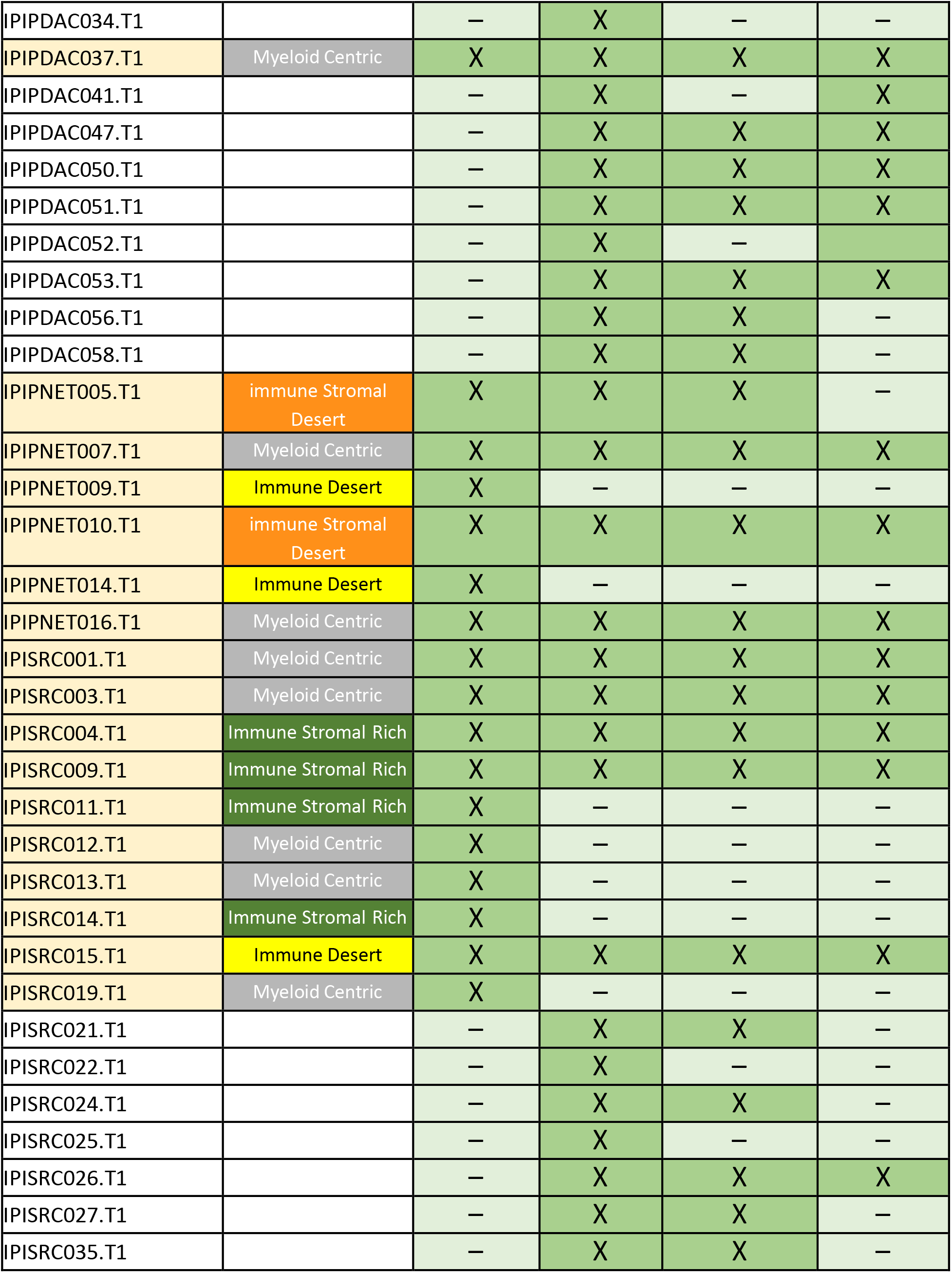
List of UCSF Immunoprofiler Initiative samples and associated flow cytometry data and bulk RNA seq data used for 3 features clustering. List of each tumor biopsy included in Figure 1A and 1C with their associated different flow cytometry data or ‘all live’ bulk RNA sequencing compartment. A cross mean that the given sample have data associated and it has been used in this manuscript. Cluster assignment for the 3 feature clustering is provided.

**Supplementary Table 3:**
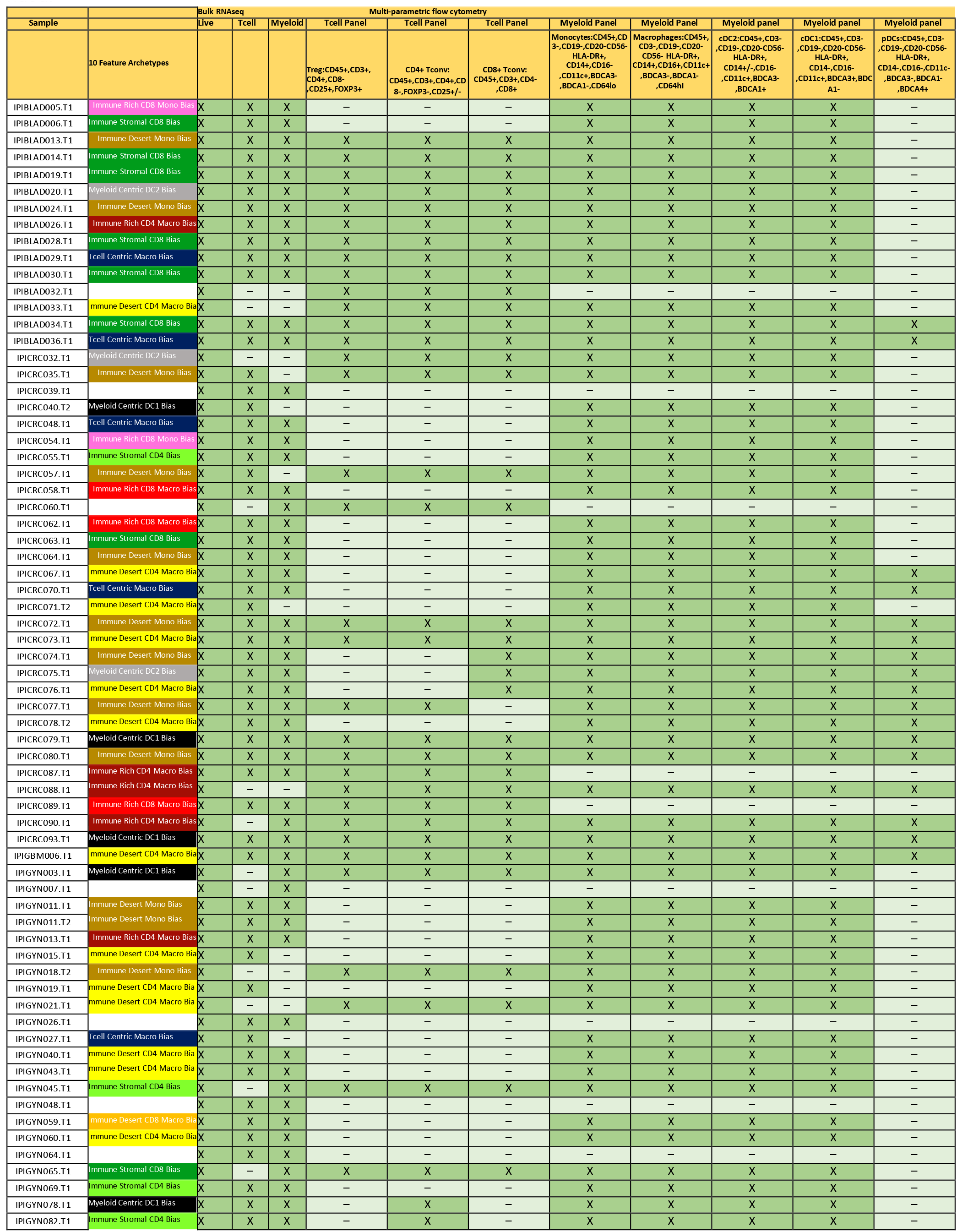
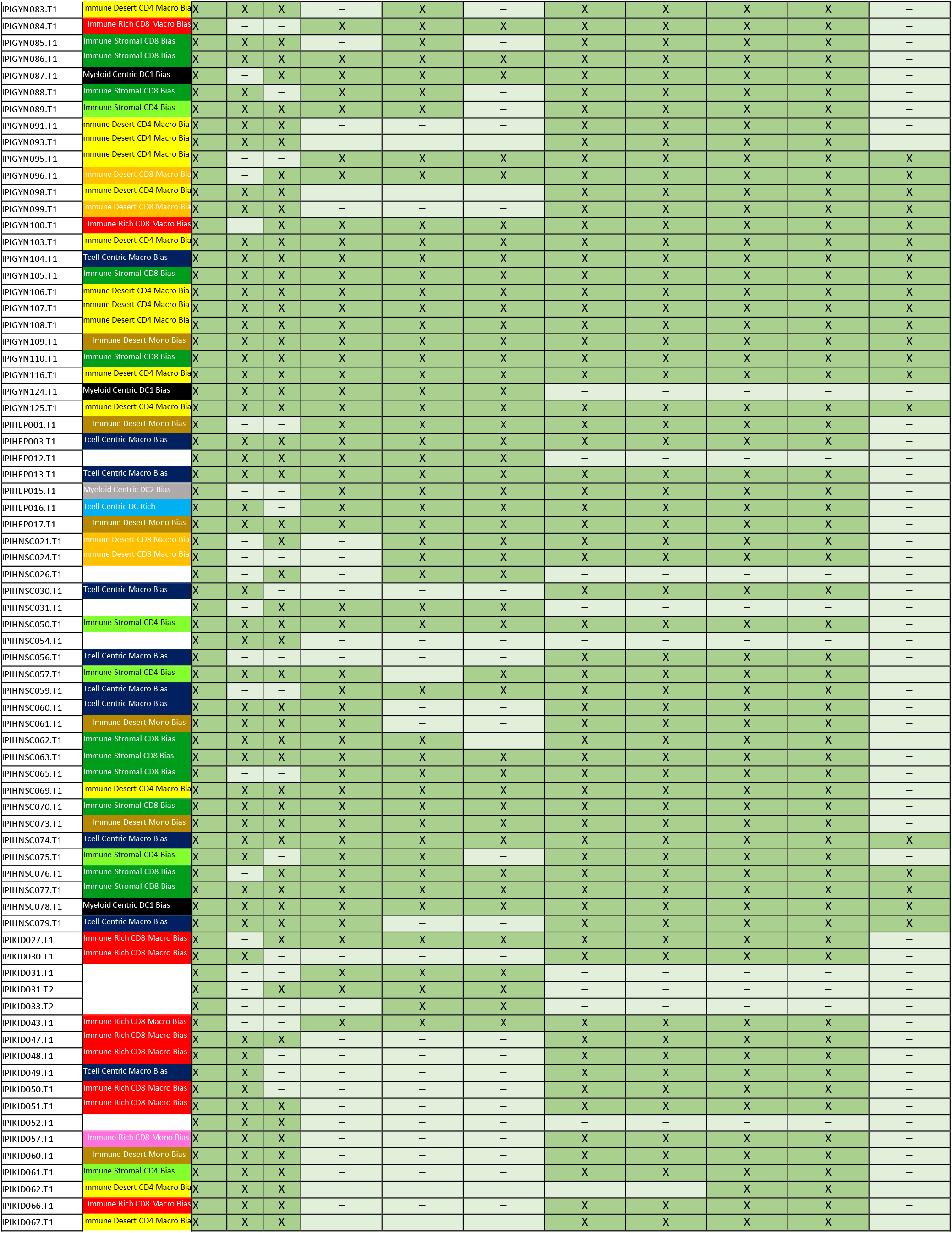
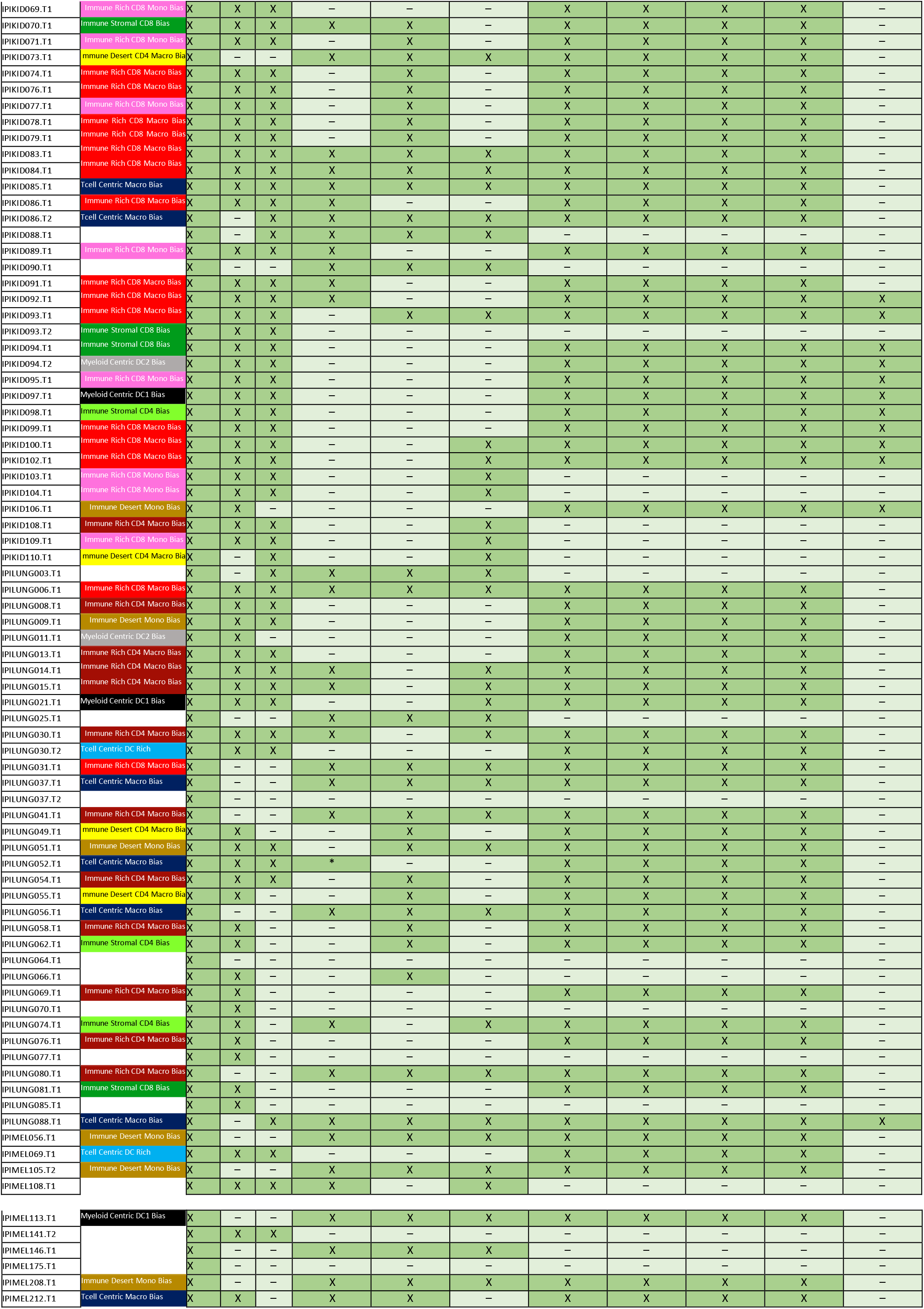
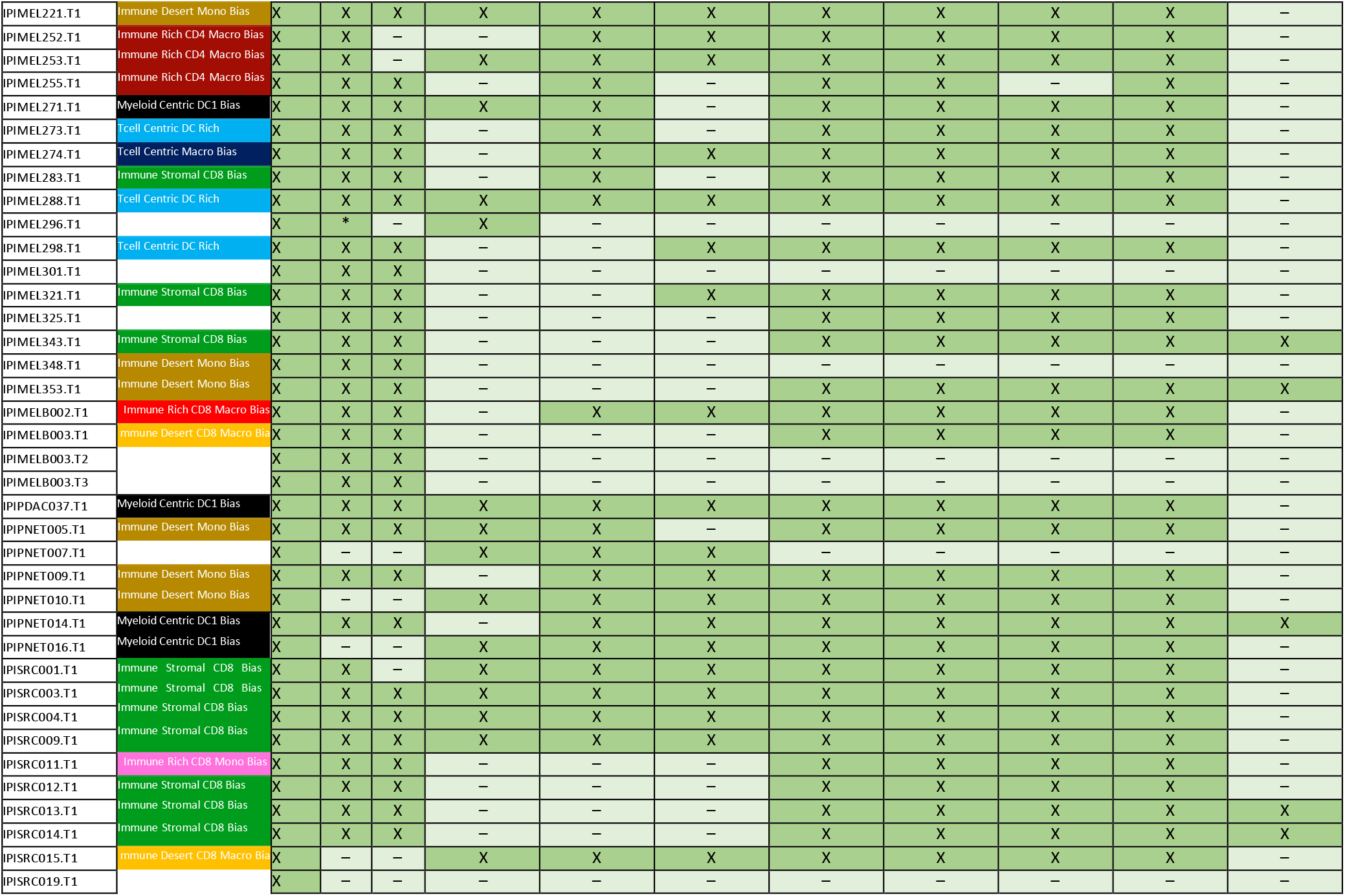
List of UCSF Immunoprofiler Initiative samples and associated flow cytometry data and bulk RNA seq data used for 6 and 10 feature clustering. List of each tumor biopsy included in the 6 and 10 feature clustering and their associated different flow cytometry data and or different bulk RNA sequencing compartment. A cross mean that the given sample have data associated and it has been used in this manuscript. Cluster assignment for the 10-feature clustering is also provided

**Supplementary Table 4:**
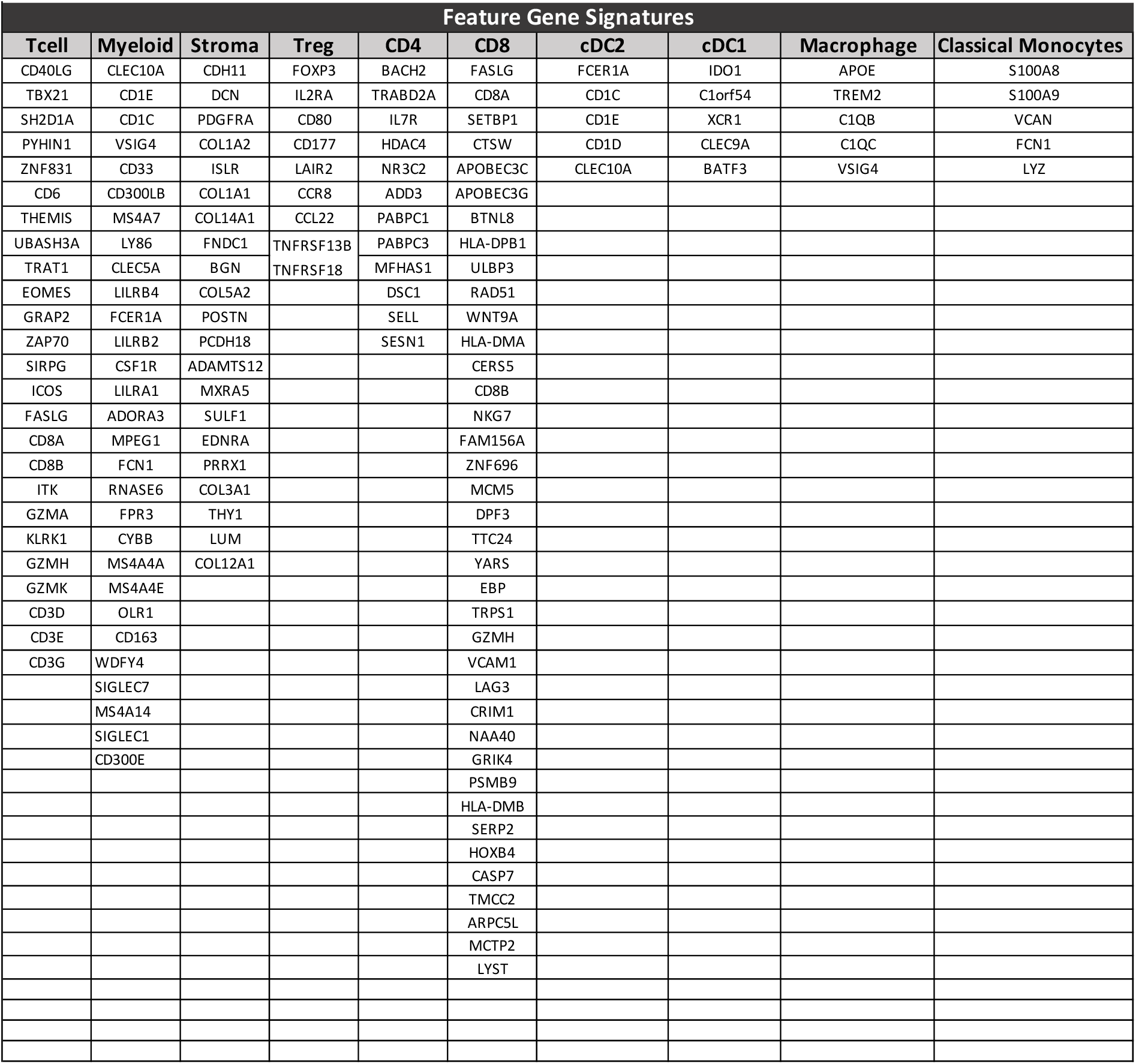
List of genes for the feature gene signatures. List of each gene included in the different feature gene signatures used to perform tumor biopsy Louvain clustering.

**Supplementary Table 5:**
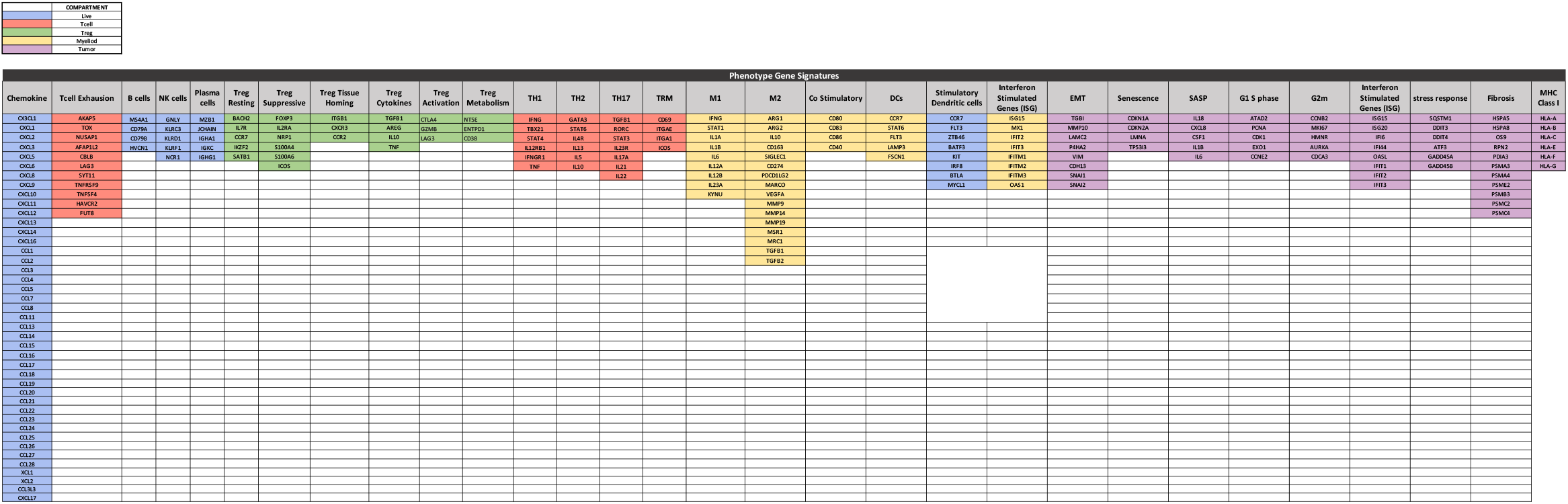
List of genes used for the phenotype gene signatures. List of each gene included in the different phenotype gene signatures used to perform scoring and hierarchical clustering of the different tumor immune archetypes.

### Material and Methods

#### KEY RESOURCES TABLE

| REAGENT or RESOURCE | SOURCE | IDENTIFIER |
| --- | --- | --- |
| Antibodies |  |  |
| Streptavidin BV421 | Biolegend | 405226 |
| anti-human CD45 APC/e780 (clone HI30) | Thermo Fisher | 47-0459-42 |
| anti-human CD3e PerCP/e710 (clone OKT3) | Thermo Fisher | 46-0037-42 |
| anti-human HLA-DR BUV395 (clone G46-6) | BD Biosciences | 564040 |
| anti-human CD56 BUV737 (clone NCAM16.2) | BD Biosciences | 564448 |
| anti-human CD4 PE/Dazzle 594 (clone S3.5) | Biolegend | 100455 |
| anti-human CD8a BV605 (clone RPA-T8) | Biolegend | 301039 |
| anti-human CD127 BV650 (clone HIL-7R-M21) | BD Biosciences | 563225 |
| anti-human CD38 AF700 (clone HIT2) | Biolegend | 303523 |
| anti-human CD25 APC (clone 2A3) | BD Biosciences | 340939 |
| anti-human CD45RO PE (clone UCHL1) | BD Biosciences | 561889 |
| anti-human PD-1 BV786 (clone EH12) | BD Biosciences | 563789 |
| anti-human ICOS BV711 (clone DX29) | BD Biosciences | 563833 |
| anti-human FoxP3 PE/Cy7 (clone 236A/E7) | Thermo Fisher | 25-4777-41 |
| anti-human CTLA-4 BV421 (clone BNI3) | BD Biosciences | 565931 |
| anti-human/mouse/rat Ki67 AF488 (clone SolA15) | Thermo Fisher | 11-5698-82 |
| anti-human CD19 PerCP/e710 (clone H1B19) | Thermo Fisher | 45-0199-42 |
| anti-human CD20 PerCP/e710 (clone 2H7) | Thermo Fisher | 45-0209-42 |
| anti-human CD56 PerCP/e710 (clone CMSSB) | Thermo Fisher | 46-0567-42 |
| anti-human CD64 BUV737 (clone 10.1) | BD Biosciences | 564425 |
| anti-human CD11c AF700 (clone 3.9) | Thermo Fisher | 56-0116-42 |
| anti-human CD16 BV605 (clone 3G8) | Biolegend | 302039 |
| anti-human CD273/PDL2 BV650 (clone MIH18) | BD Biosciences | 563844 |
| anti-human/mouse TREM2 APC (clone 237920) | R&D Systems | FAB17291A |
| anti-human CD304 PE (clone 12C2 ) | Biolegend | 354503 |
| anti-human CD1C/BDCA-1 PE/Cy7 (clone L161) | Biolegend | 331515 |
| anti-human CD197 BV421 (clone G043H7) | Biolegend | 353207 |
| anti-human BDCA-3 FITC (clone AD5-14H12) | Miltenyi | 130-098-843 |
| anti-human PDL1 BV786 (clone MIH1) | BD Biosciences | 563739 |
| anti-human CD14 BV711 (clone M5E2) | Biolegend | 301837 |
| Bacterial and Virus Strains |  |  |
| Biological Samples |  |  |
| Human tumor samples | UC San Francisco | IRB # 20-31740 |
| Chemicals, Peptides, and Recombinant Proteins |  |  |
| N/A |  |  |
| Critical Commercial Assays |  |  |
| N/A |  |  |
| Deposited Data |  |  |
| GEO |  |  |
| Experimental Models: Cell Lines |  |  |
| N/A |  |  |
| Experimental Models: Organisms/Strains |  |  |
| N/A |  |  |
| Oligonucleotides |  |  |
| N/A |  |  |
| Recombinant DNA |  |  |
| N/A |  |  |
| Software and Algorithms |  |  |
| Python (2.7.15 & 3.7.2) | (Rossum, 1995) | https://www.python.org |
| Pandas (0.24.1, 1.0.5) | (McKinney, 2010) | https://pandas.pydata.org/ |
| Seaborn (0.9.0, 0.10.1) | (Waskom et al., 2020) | https://seaborn.pydata.org |
| Matplotlib (2.2.3, 3.2.2) | (Hunter, 2007) | https://matplotlib.org |
| Lifelines | (Davidson-Pilon et al., 2020) | https://lifelines.readthedocs.io/en/latest/ |
| Scanpy 1.5.1 | (Wolf et al., 2018) | https://scanpy.readthedocs.io/en/stable |
| SciPy | (Virtanen et al., 2020) | https://www.scipy.org/ |
| BWA-mem | (Li, 2013) | http://bio-bwa.sourceforge.net/ |
| RSEM | (Li and Dewey, 2011) | http://deweylab.biostat.wisc.edu/rsem/README.html |
| Imaris v9.5 |  | https://imaris.oxinst.com/ |
| STAR | (Dobin et al., 2013) |  |
| limma | (Ritchie et al., 2015) | https://bioconductor.org/packages/release/bioc/html/limma.html |
| edgeR | (Robinson et al., 2010) | https://bioconductor.org/packages/release/bioc/html/edgeR.html |
| voom | (Law et al., 2014) | https://www.rdocumentation.org/packages/limma/versions/3.28.14/topics/voom |
| R | (R Development Core Team, 2010) | https://www.r-project.org/ |
| Other |  |  |

#### METHOD DETAILS

##### Human tumor collection of the UCSF Immunoprofiler Initiative (IPI)

Tumor samples for the Immunoprofiler was transported from various cancer operating rooms (ORs) as well as from outpatient clinics. All patients consented by the UCSF IPI clinical coordinator group for tissue collection under a UCSF IRB approved protocol (UCSF IRB# 20-31740). Samples were obtained after surgical excision with biopsies taken by Pathology Assistants to confirm the presence of tumor cells. Patients were selected without regard to prior treatment. Freshly resected samples were placed in ice-cold PBS or Leibovitz’s L-15 medium in a 50 mL conical tube and immediately transported to the laboratory for sample labeling and prepare either the whole tissue for digestion into single-cell suspension or a part of the tissue was sliced and preserved for imaging analysis.

##### Human tissue digestion and Multiparametric Flow Cytometry staining and sorting

Tumor or metastatic tissue was thoroughly chopped with surgical scissors and transferred to GentleMACs C Tubes (Miltenyi Biotec) containing 20 uL/mL Liberase TL (5 mg/ml, Roche) and 50 U/ml DNAse I (Roche) in RPMI 1640 per 0.3 g tissue. GentleMACs C Tubes were then installed onto the GentleMACs Octo Dissociator (Miltenyi Biotec) and incubated for 45min according to the manufacturer’s instructions. Samples were then quenched with 15 mL of sort buffer (PBS/2% FCS/2mM EDTA), filtered through 100 um filters and spun down. Red blood cell lysis was performed with 175 mM ammonium chloride if needed.

Cells were then incubated with Human FcX (Biolegend) to prevent non-specific antibody binding. Cells were then washed in DPBS and incubated with Zombie Aqua Fixable Viability Dye (Thermo). Following viability dye, cells were washed with sort buffer and incubated with cell surface antibodies mix diluted in the BV stain buffer (BD Biosciences) following manufacturer instruction for 30 minutes on ice in the dark and subsequently fixed in either Fixation Buffer (BD Biosciences) or in Foxp3/Transcription Factor Staining Buffer Set (eBioscience) if intracellular staining was required.

##### Cell sorting for bulk RNA sequencing

| Bulk RNAseq compartment | Gating Strategy | Potential Caveats |
| --- | --- | --- |
| All viable cells 'Live' compartment | All cells Viability dye negative<br>LIVE/DEAD™Fixable Aqua<br>Dead Cell Stain Cat:L34957 | This was performed after tissue enzymatic digestion and allow us to sequence only viable cells after isolation. Therefore it is slightly different than bulk RNAseq from whole tissue prior digestion as it was done in the TCGA dataset. |
| Conventional CD4+ and CD8+ T cells compartment | CD4+ and CD8+ Tconv:<br>CD45+, CD3+<br>, CD4+, CD8-, CD25- CD19/20-,<br>CD56- | Using this gating strategy we aimed to enrich for both Conventional CD4+ and CD8+ T cells but we don't separate CD4+ and CD8+ |
| Regulatory CD4+ T cells compartment | Treg: CD45+, CD3+, CD4+, CD8-<br>, CD25+, CD19/20-, CD56- | Using this gating strategy we aimed to enrich for Regulatory T cells using CD25 expression but as we can't use FOXP3 for sorting unpermeabilised viable cell this compartment may contain some activated CD4+ T cells. |
| Myeloid cells compartment | CD45+, CD3-, CD19-, CD20-<br>CD56- HLA-DR+, Cd11b+/- | Using this gating strategy we aimed to enrich for tumor associated myeloid cell however this compartment may contain plasma cells which would have down-regulate CD19/20 but still express HLA-DR |
| Cd44+ CD90 stromal cells compartment | CD45-, CD44+, CD90+ | Using this gating strategy we aimed to enrich for one type of stromal cells however this gating strategy doesn't encompass all stromal cell subtype present in tumor |
| CD45- Tumor cell compartment | CD45-, CD44-, CD90- | Using this gating strategy we aimed to enrich for tumor cells however this gating strategy is permissive and this compartment may contain other tumor associated non immune cell types |

For cell sorting single cell suspension was stained with cell surface antibodies mix diluted in BV stain buffer (BD Biosciences) following manufacturer instruction for 30 minutes on ice in the dark and subsequently washed three time and resuspended in of sort buffer (PBS/2% FCS/2mM EDTA) after filtering in 100um filter (thermo). When possible, a maximum of 6 different tumor associated cell subsets were sorted with a BD FACSAria Fusion into 1.5mL Eppendorf tubes containing 150ul of lysis buffer used for mRNA extraction (Invitrogen). The 6 different tumor associated cell types between 5000 and 50000 cells were sorted based their protein expression profile as describe here: ‘live cells’ all cells negative for the viability dye ; Tconv: negative for viability dye CD45+ CD3+ CD19− CD56− CD25− ; Treg: negative for viability dye CD45+ CD3+ CD19− CD56− CD4+ CD25+ ; Myeloid: negative for viability dye CD45+ CD3− CD19− CD56− HLADR+ CD11b+/− ; CD90+ CD44+ Stroma: negative for viability dye CD45− CD44+ CD90+ ; Tumor: negative for viability dye CD45− CD44− CD90− EPCAM+/−. Names of the different sorted populations were given based on the presumptive populations and we accepted that both due to slight sorting contamination and due to the limits of antibodies to completely ‘define’ a population, that other populations would occasionally infiltrate these e.g., some ‘activated’ CD4 cells express CD25 and might infiltrate this RNA sample

##### Cell sorting for single-cell RNA sequencing of the tumor associated mononuclear phagocytic cells

For scRNA-seq, live CD3-CD19/20-CD56− SSC-A dim CD16 dim (to exclude neutrophils) cells were sorted from a melanoma involved draining LN on a BD FACSAria Fusion. After sorting, cells were pelleted and resuspended at 1×103 cells/ml in 0.04%BSA/PBA and loaded onto the Chromium Controller (10X Genomics). Samples were processed for single-cell encapsulation and cDNA library generation using the Chromium Single Cell 3’ v2 Reagent Kits (10X Genomics). The library was subsequently sequenced on an Illumina HiSeq 4000 (Illumina).

##### Imaging Staining and Acquisition

###### H&E

H&E slides were prepared in Leica Autostainer XL; slides stained in hematoxylin (Thermo Scientific Shandon Instant Hematoxylin cat. 6765015) for 7 minutes and in Eosin (Thermo Scientific Shandon Instant Eosin-Y Alcoholic cat. 531946) for 20 seconds.

###### Immunofluorescence (IF) Staining

IF staining was performed on the Ventana Discovery Ultra autostainer using Discovery reagents (Ventana Medical Systems) according to the manufacturer’s instructions, except as noted. After deparaffinization, antigen retrieval was performed with Cell Conditioning 1 (CC1) solution (cat. 950-124) for 64 minutes at 95°C. Primary antibody CD45 (D9M8I by CST #13917), was incubated for 20 minutes at 36°C. Secondary antibodies (cat. 760-149) were incubated for 12 minutes. Endogenous peroxidase was inhibited by Discovery Inhibitor (cat. 760-4840) for 8 minutes and non-specific binding blocked with Goat block (cat. 760-6008) for 4 minutes. The primary antibody was visualized with Discovery Rhodamine 6G Kit (cat. 760-244). Finally, slides were counterstained with DAPI (Akoya cat. FP1490) for 8 minutes.

###### Imaging Slides

IF slides were scanned in a whole slide scanner AxioScan.Z1 (Zeiss) with Plan-Apochromat 20x/0.8 M27 objective and images were captured by Orca-Flash 4.0 v2 CMOS camera (Hamamatsu). Filters used for specific fluorophores are: Spectral Gold (Semrock) filter was used for Rhodamine 6G and 87 HE DAPI (Zeiss) for DAPI. H&E slides were scanned in brightfield with the same objective and a HV-F202SCL CCD camera (Hitachi).

##### Bulk RNA-Sequencing

###### Library Preparation

The mRNA was isolated via DinaBead Direct, then converted into amplified cDNA using Tecan Ovation RNA-Seq System V2 kit, following the manufacturer protocol. The dsDNA went through tagmentation, amplification and clean up with AMPure XP beads steps, using the Illumina Nextera XT DNA Library Prep Kit. Quality control analysis was performed on the resulting pooled libraries with the Agilent Bioanalyzer HS DNA chip to assess fragment size distribution and concentration.

###### Sequencing

The pooled libraries were then sequenced via single-read MiSeq/MiniSeq to check if more than 10 percent of the reads aligned to coding regions and contained more than 1,000 unique reads in total. The libraries were then re-pooled based on the percentage of reads in a coding region. These libraries were submitted the UCSF Center for Advanced Technology for 100bp paired end (PE100) sequencing on the HiSeq4000

##### Bioinformatic Data Processing

###### Bulk RNAseq

The RNAseq reads were first aligned to reference of rRNA and mitochondrial rRNA using BWA-mem (Li, 2013) to deplete the dataset of any remaining rRNA. The remaining reads were aligned to the Ensembl GRCh38.85 transcriptome build using STAR (Dobin et al., 2013). Gene expression was computed from the alignments in counts and TPM (transcripts per million) using RSEM (Li and Dewey, 2011).

###### Data pre-processing of 10x Genomics Chromium scRNA-seq data

Feature-barcode matrices were obtained for sample by aligning the raw fastqs to GRCh38 reference genome (annotated with Ensembl v85) using the Cellranger count. Raw feature-barcode matrices were loaded into Seurat 3.1.5 (Stuart et al., 2019) and genes with fewer than 3 UMIs were dropped from the analyses. Matrices were further filtered to remove events with greater than 20% percent mitochondrial content, events with greater than 50% ribosomal content, or events with fewer than 100 total genes. The cell cycle state of each cell was assessed using a published set of genes associated with various stages of human mitosis (Dominguez et al., 2016).

###### Data quality control and Normalization for scRNA-seq data

The filtered count matrices were normalized, and variance stabilized using negative binomial regression via the scTransform method offered by Seurat (Hafemeister and Satija, 2019). The effects of mitochondrial content, ribosomal content, and cell cycle state were regressed out of the normalized data to prevent any confounding signal. The normalized matrices were reduced to a lower dimension using Principal Component Analyses (PCA) and the first 30 principal coordinates per were subjected to a non-linear dimensionality reduction using tSNE reduction. Clusters of cells sharing similar transcriptomic signal were identified using the Louvain algorithm, and clustering resolutions varied between 0. 6 and 1.2 based on the number and variety of cells obtained in the datasets.

###### Single Cell RNAseq intra-sample heterotypic doublet detection

All libraries were further processed to identify heterotypic doublets arising from the 10X sample loading. Processed, annotated Seurat objects were processed using the DoubletFinder package (McGinnis et al., 2019). Briefly, the cells from the object are modified to generate artificial duplicates, and true doublets in the dataset are identified based on similarity to the artificial doublets in the modified gene space. The prior doublet rate per library was approximated using the information provided in the 10x knowledgebase (https://kb.10xgenomics.com/hc/en-us/articles/360001378811) and this was corrected to account for homotypic doublets using the per-cluster numbers in each dataset. Heterotypic doublets were removed from the major analysis (see platelet-first analysis).

###### Imaging bioinformatics

Each image was processed and analyzed in Imaris v9.5 (Bitplane Inc.). We first applied a median filter with size 3×3×1 to every channel of the image. The background of each channel was then subtracted using a filter width of 100um. Next, we used the Spots function to detect cell nuclei in the DAPI channel. We then applied a binary threshold on the max intensity of the CD45 channel to classify each spot as positive or negative for CD45.

We then cropped out a 650um x 650um section containing the tumor border from each histological image. This same section of tissue was found in the immunofluorescence image by looking for tissue features shared between the Immunofluorescence (IF) and histological images. Using the CD45 counts described above, two representative images from each immune class were selected.

##### Assembling Cohorts

###### Immunoprofiler (IPI)

An initial set of 427 bulk RNAseq patient samples in the live compartment were evaluated based on an in-house metric, the EHK score, that serves as a measure of data quality. Each sample is given a score of 0 through 10 depending on the number of EHK genes that are expressed above a precalculated minimum threshold. A score of 10 represents the highest quality data where 10 out of 10 EHK genes are expressed above the minimum threshold. Filtering for samples with an EHK score of EHK8, EHK9 and EHK10 reduced the sample set to 298 patient samples. The sample set was then filtered to remove all adjacent normal samples and all biological replicates, reducing the sample set further to 260 samples. These 260 patient samples are the IPI cohort along with 199 overlapping tcell compartment samples and 189 myeloid compartment samples.

###### TCGA

Tumor RNAseq counts and TPM along with curated clinical data for 13 indications (Bladder urothelial carcinoma (BLCA), Colon adenocarcinoma (COAD), glioblastoma multiforme (GBM), Head and Neck squamous cell carcinoma (HNSC), Kidney renal clear cell carcinoma (KIRC), Liver hepatocellular carcinoma (LIHC), Lung adenocarcinoma (LUAD), Ovarian serous cystadenocarcinoma (OV), Pancreatic adenocarcinoma (PAAD), Sarcoma (SARC), Skin Cutaneous Melanoma (SKCM), Uterine Corpus Endometrial Carcinoma (UCEC), Uterine Carcinosarcoma (UCS)), from the Toil recompute (Vivian et al., 2017) data in the TCGA Pan-Cancer (PANCAN) cohort, were downloaded from the UCSC Xena browser (Goldman et al., 2019). The initial set of 4677 tumor samples was filtered down to include primary solid tumors and metastatic sample types (sample type codes = 01 & 06) only, in order to parallel the IPI cohort sample types as accurately as possible. This reduced the patient sample set to 4341 tumor samples.

#### QUANTIFICATION & STATISTICAL ANALYSIS

##### Feature Gene Signatures

###### Tcell, Myeloid, CD90+ CD44+ Stroma and Treg

Gene signatures for each of the features (Supplementary Table4) were generated by performing differential gene expression (DEG) analysis, between the different sorted RNAseq compartments, using edgeR (Robinson et al., 2010), limma (Ritchie et al., 2015) and Voom (Law et al., 2014), a function of limma that modifies RNAseq data for use with limma. Expression counts for all EHK10 samples in each of the compartments (tcell, myeloid, stroma, treg and tumor) were selected to ensure the DEG analysis was done using the highest quality data. The median adjusted p-value for these DEGs were values much less than 1.00 x 10^−55^ across all comparisons. For the Tcell feature gene signature, DEG analysis was done between the tcell compartment and the stroma, myeloid, and tumor compartments separately. The intersection of the top 50 genes by log fold change (logFC) for each of these three comparisons was designated the Tcell feature gene signature. Similarly, a CD90+ CD44+ Stroma feature gene signature was designated as the intersection between DEG comparisons of the stroma compartment versus the myeloid, tumor and tcell compartments. For the Myeloid feature gene signature, it was necessary to use the top 100 genes by logFC from the DEG between the myeloid compartment and tcell compartment to compensate for the fact that myeloid cells are more auto fluorescent and thus more likely to contaminate the tcell compartment sort. The stroma and tumor compartments are sorted from CD45− and are less likely to experience this issue. The Treg feature gene signature follows all the same steps as the Tcell feature gene signature but with four DEG comparisons instead of three, the treg compartment versus the tcell, myeloid, stroma and tumor compartments. The resulting intersection of these four comparisons was designated the Treg feature gene signature. The process of feature gene signature selection for each of the four features was visualized with volcano plots that showed the DEG results and Venn diagrams that showed the intersection of the top genes from each DEG and the resulting feature gene signature.

###### CD4 and CD8

CD4 high and CD8 high samples were identified based on the flow proportion of (CD4+, CD25-, FOXP3-) / CD3+ and (CD8+, CD4-) / CD3+ respectively. There were nine samples that were both EHK10 in the tcell compartment of RNAseq and had a (CD4+, CD25-, FOXP3-) / CD3+ flow proportion greater than 60%, these were labeled CD4 high. Similarly, there were 12 samples that were EHK10 in the tcell compartment of RNAseq that had a (CD8+, CD4-) / CD3+ flow proportion greater than 60%, these were labeled CD8 high. The CD4 gene signatures were generated by performing DEG analyses using edgeR and voom on the tcell compartment counts of CD4 high samples versus CD8 high samples. The top 50 genes by adjusted P-value were selected and all genes within this gene set that had a positive logFC were designated the CD4 feature gene signature. The same was done for the CD8 feature gene signature except the DEG was CD8 high samples versus CD4 high samples (Supplementary Table 4). A hierarchical clustered heatmap was made using the CD4 and CD8 gene signature logCPM values for all tcell compartment patient samples. The logCPM values were mean-centered and scaled prior to clustering using the Euclidean distance metric and the Ward linkage method.

###### Macrophages, Monocytes, cDC1 and cCD2

Gene signatures were generated from single cell RNAseq. Differential gene expression (DGE) between each clusters identified were performed on log-normalized gene counts using the Poisson test provided by the FindMarkers/FindAllMarkers functions in Seurat. Genes with > 0.4 log-fold changes, an adjusted p value of 0.05 (based on Bonferroni correction), at least 35% expressed in tested groups, were regarded as significantly differentially expressed genes (DEGs). Cluster marker genes were identified by applying the DE tests for upregulated genes between cells in one cluster to all other clusters in the dataset. Genes were curated based on log-fold changes and a minimum expression by the other cluster/cell types. The different subsets of mononuclear phagocytic cells were identified by comparing cluster marker genes with public sources referenced in the text. (Supplementary Table 4)

###### Phenotype Gene Signatures

Phenotype gene signatures (Supplementary Table 5) were used to further characterize the immune archetypes obtained by clustering feature scores generated from the feature gene signatures. The phenotype gene signatures were assembled from standard gene signatures, derived from analysis of IPI samples or curated from gene sets found in literature.

The Tcell Exhaustion gene signature was generated by correlating the expression of all protein coding genes in EHK10 samples from the tcell compartment with the expression of five genes, CTLA4, PDCD1, CD38, HAVCR2 and LAG3, a subset of a published T cell exhaustion gene signature (B et al., 2018). The top 50 genes by Spearman’s rho for each of the five correlations were selected as gene sets and genes that fell in the intersection of at least four of these gene sets were designated as the Tcell Exhaustion feature gene signature. The Spearman’s rho of the CTLA4, PDCD1, CD38, HAVCR2 and LAG3 against the Tcell Exhaustion gene signature genes with additional known exhaustion genes were plotted on heatmap to illustrate the gene signature selection criteria.

The Chemokine gene signature was a selection of 39 chemokines of interest. Hierarchical clustered heatmaps were made to assess chemokine expression in archetypes. The median TPM expression per archetype of each of the 39 chemokines in the Chemokine gene signature were clustered using the Euclidean distance metric and the Ward linkage method. A bubble plot was made to compare the median TPM expression of Chemokine gene signature genes in IPI and TCGA in each archetype. The bubble size represented values of the median TPM in each archetype transformed to a z-score.

The ISG, Senescence, EMT, Fibrosis, Cell Stress DNA damage, Cell Cycle G1-S and Cell Cycle G2-M genes signatures were from literature (Kinker et al., 2019,munoz dp et al, wiley cd et al). A bubble plot was made to visualize the median TPM expression per archetype in the tumor compartment. The bubble size and color represented values of the median TPM in each archetype, transformed to a z-score.

The NK cell gene signature was from literature (Barry et al., 2018), the B cells gene signature was from (Chen et al., 2020; van Galen et al., 2019) and the Plasma cells gene signature was from (Chen et al., 2020; van Galen et al., 2019)A bubble plot was made to visualize the median TPM expression per archetype in the live compartment. The bubble size and color represented values of the median TPM in each archetype, transformed to a z-score.

The Th1, Th2, Th17, Trm and Tex gene signatures were from (B et al., 2018; Kumar et al., 2017; Savas et al., 2018; Zhou et al., 2009). A bubble plot weas made to visualize the median TPM expression per archetype in the tcell compartment. The bubble size and color represented values of the median TPM in each archetype, transformed to a z-score. A hierarchical clustered heatmap was made to visualize the Th1, Th2, Th17, Trm and Tex gene signature genes in the samples of the Immune Rich CD8 Bias archetype. The TPM of the gene signature genes for the archetype samples were clustered using a Euclidian distance metric and the Ward linkage method. The clustered heatmap was visualized using Seaborn with rug for the indications in the archetype samples

The Resting Tregs, Suppressive Tregs, Tissue Homing Tregs, Treg Activation, Treg Cytokines, Treg Metabolism were from (Arce Vargas et al., 2018; Ephrem et al., 2013; Plitas et al., 2016; Zemmour et al., 2018)

The M1, M2 and Costimulatory molecule gene signatures were from (Biswas et al., 2013; Cassetta et al., 2019; Maier et al., 2020; Roberts et al., 2016). A bubble plot was made to visualize the median TPM expression per archetype in the myeloid. The bubble size and color represented values of the median TPM in each archetype, transformed to a z-score.

The MHC Class I gene signature was from HGNC (https://www.genenames.org). The ISG gene signature was from (Combes et al., 2021). A box plot was made to visualize the scores per archetype that were calculated in the live compartment.

A hierarchical clustered heatmap was made to visualize the Th1, Th2, Th17, Trm, Tex, Resting Tregs, Suppressive Tregs, Tissue Homing Tregs, Treg Cytokines, Treg Activation, Treg Metabolism, NK cells, Plasma cells, B cells, M1, M2, Costimulatory molecule, and ISG gene signature gene expression. These genes for each of the phenotype gene signatures were assessed different RNAseq compartments as described above. The median TPM expression per archetype of each of the genes in the gene signatures were clustered using correlation distance metric and the average linkage method.

##### Immune Archetype Gene Signatures

###### Live Archetype Gene Signature

Archetype gene signatures were generated by DEG analysis, between the live compartment counts of samples in each of the 12 archetypes, using limma and Voom. The intersection between the top 3000 genes by logFC of each of 11 DEGs per archetype was assigned as an initial gene signature. If the initial gene signatures had less than 20 genes it was designated as the archetype gene signature. Otherwise, the archetype gene signature was designated as the top 20 genes with both the lowest coefficient of variation (CV) of the log10(TPM+0.001) expression and non-zero expression in at least 80% of the samples in the archetype. A hierarchical clustered heatmap was made to visualize archetype gene signature gene expression. The median TPM expression per archetype of each of the genes in the gene signatures were clustered using correlation distance metric and the average linkage method.

###### Tumor Archetype Gene Signature

Archetype gene signatures were generated by DEG analysis, between the combined tumor and epcam compartment counts of samples in each of the 12 archetypes, using edgeR and Voom. The intersection between the top 100 genes by logFC of each of 11 DEGs per archetype was assigned as an initial gene signature. If the initial gene signatures had less than 10 genes it was designated as the archetype gene signature. Otherwise, the archetype gene signature was designated as the top 10 genes with both the lowest coefficient of variation (CV) of the log10(TPM+0.001) expression and non-zero expression in at least 80% of the samples in the archetype. A hierarchical clustered heatmap using correlation distance metric and the average linkage method was made for the median TPM expression per archetype of each of the genes in the gene signatures.

##### Scores

###### Gene Signature Score

The feature gene signature scores were calculated using an m x n matrix where m represented the TMM (Robinson and Oshlack, 2010) normalized logCPM (log2 counts per million) expression of the feature signature genes and n represented the selected sample set.

The expression of each gene was converted to percentile ranks across the samples using the SciPy (Virtanen et al., 2020) Python module.

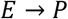

where: E = m x n matrix of gene expression

P = m x n matrix of gene expression percentile ranks across samples

A score was generated for each sample n follows:

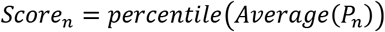

where: P_n_ = column n of P corresponding to sample n

(The second percentile transformation was employed to increase the spread between the scores). Boxplots were generated to visualize the distribution of feature scores by cancer indication with statistical analysis between archetypes performed using the statannot python package. We used the Mann-Whitney test between two archetypes, as we could not assume normal distributions, with the Bonferroni correction for multiple comparisons. The significance levels were annotated on the boxplots using the P-Value annotation legend below

| Annotation | P-Value Range |
| --- | --- |
| ns | $5.00e-02 < p \leq 1.00e+00$ |
| * | $1.00e-02 < p \leq 5.00e-02$ |
| ** | $1.00e-03 < p \leq 1.00e-02$ |
| *** | $1.00e-04 < p \leq 1.00e-03$ |
| **** | $p \leq 1.00e-04$ |

For the IPI cohort the gene signature scores for the Tcell, Myeloid, CD90+ CD44+ Stroma, Treg, Chemokine features were calculated using the live compartment. The CD4, CD8 and T cell Exhaustion were calculated using the tcell compartment. The Macrophage, Classical Monocyte, CDC1, cDC2 were calculated using the myeloid compartment. The MHC Class I was calculated using the tumor compartment.

For the TCGA cohort the gene signature scores for the Tcell, Myeloid and CD90+ CD44+ Stroma features were calculated for all indications. Cross-Whisker plots, that show the median value, the interquartile range (IQR) in both the x and y directions and the identity line, were made to compare the median Tcell, Myeloid and CD90+ CD44+ Stroma feature scores in the IPI cohort with the TCGA cohort. Based on the Cross-Whisker plots only the BLCA, COAD, UCS, UCEC, OV, HNSC, KIRC, SARC and SKCM, indications were used for further analysis as their, median scores per indication appeared to correlate well with the IPI patient samples. The rationale behind including only indications that correlated well with IPI was to ensure that further analysis in TCGA would have extrapolated relevance in the IPI patient samples. The excluded indications’ lack of correlation with IPI patient samples could be attributed to variety of reasons that cannot be easily controlled, e.g. technical variability in upstream processing steps and cohort race differences due to local demographic biases.

###### Gene Signature Score Validation

Based on the assumption that the flow cytometry population ratios represent the true feature abundance in the patient sample, correlation between flow cytometry population ratios and feature gene signatures were used to validate the feature gene signatures. On average 9 to 10 samples per indication, that had both flow cytometry data and RNAseq data in the relevant compartments, were selected (Supplementary Table 3). There were fewer samples selected in PDAC(Pancreatic), PNET(Neuroendocrine) and GBM(Glioblastoma) indications as there were not many representative samples for theses indications in the IPI cohort. Correlation plots were generated using the scores and their corresponding flow population ratios and the Spearman’s rho was calculated for each of the correlations.

Additionally, the Tcell and Myeloid feature samples were submitted to CIBERSORT (Newman et al., 2015). The TPMs of all protein coding genes of the samples were input as the mixture file, the built in LM22 gene signatures were used and quantile normalization was disabled as this was the recommended setting for RNAseq data. The CIBERSORT score used for the Tcell feature was the summation of the T cells CD8, T cells CD4 naïve, T cells CD4 memory resting, T cells CD4 memory activated, T cells follicular helper, T cells regulatory (Tregs) and T cells gamma delta cell type relative fractions. The CIBERSORT score used for the Myeloid feature was the summation of the Macrophages M0, Macrophages M1, Macrophages M2, Dendritic cells resting, Dendritic cells activated, Mast cells resting, Mast cells activated, Eosinophils and Neutrophils cell type relative fractions. Correlation plots were generated using these CIBERSORT scores and their corresponding flow population ratios and the Spearman’s rho was calculated for each of the correlations.

###### Flow Score

The flow scores for the flow cytometry population ratios were calculated by converting the ratios to percentiles and then further transforming the percentile to percentiles to increase the spread in the scores. Boxplots were generated to visualize the scores by cancer indication using Seaborn.

###### Calculated Score for Missing Data

The IPI cohort consists of 260 patient samples that all have a live compartment sequenced. However, there are only 199 overlapping patient samples with the tcell compartment sequenced and 189 overlapping patient samples with the myeloid compartment sequenced. As some feature gene signature scores, noted above, were evaluated in the tcell and myeloid compartment a need for a method to calculate scores for missing data was warranted. Gene signature scores were calculated for missing patient samples by leveraging the fact that there was very high correlation between gene signatures scores and their corresponding flow population ratios. For example, the correlation between the Tcell gene signature score and the CD3+ / live population ratio from flow cytometry had a Spearman’s rho of 0.91. The missing scores were calculated by modeling the correlation using linear regression and using the regression equation and the flow population ratios to calculate the score. Of the 61 missing tcell compartment samples 5 did not have flow data and of the 71 missing myeloid compartment samples 15 did not have flow data.

###### Live Archetype Gene Signature Score

Using TCGA data, a gene signature score for each archetype was calculated using the gene signature score method above. Each sample has 12 scores for each archetype. Each sample was assigned to the archetype for which it had the highest score and if the highest score was tied between archetypes, the sample was excluded from the analysis. Survival analysis was performed per indication and across all indications using lifelines (Davidson-Pilon et al., 2020), a survival analysis Python library. If an archetype had less than fifteen representative samples, it was excluded from analysis. Pie charts were made to compare the archetype abundance per indication between IPI and TCGA.

##### Feature Clustering

The feature gene signature scores in the IPI cohort were clustered using SCANPY (Wolf et al., 2018) a Python-based toolkit. The initial clustering was done using three features (IPI-3f), Tcell, Myeloid and CD90+ CD44+ Stroma; subsequent clustering was done with additional features: six features (IPI-6f), which was IPI− 3f + Treg, CD4 and CD8 and ten features (IPI-10f), which was IPI-6f + Macrophage, Classical Monocytes, cDC1 and cDC2. The feature clustering was done in three steps. In the first step, a neighborhood graph was constructed using the k-nearest neighbors (KNN) algorithm with Euclidean distances. In the second step, the neighborhood graph was clustered using the Louvain method (Blondel et al., 2008), a community detection algorithm that maximizes network modularity. In the third step, the resulting clustering was evaluated using the Davies Bouldin Index (DBI) (Davies and Bouldin, 1979), a metric that assesses the ratio of the intra-cluster distance to inter-cluster distance. The lower the DBI the better the separation between clusters, the more compact the samples within clusters and hence the better the clustering. The clustering steps were repeated incrementally over values of n between 3 to 300 for the nearest neighbors for KNN. At each iteration of n, the values for the resolution parameter for Louvain clustering were varied between 0.3 to 2 in 0.1 increments. The formula below was used to optimize the nearest neighbor and resolution parameters for final cluster selection.

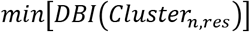

Where: n = [3,4,5…,300] (KNN nearest neighbor)

res = [03,0.4,0.5…,2] (resolution)

In the TCGA cohort only the 3f gene feature signature score clustering (TCGA-3f) was done as above with the exception of the values for the range of n for KNN which was 3-1,500 as the number of samples in the TCGA cohort were greater.

The final clustering selections for the IPI-3f, IPI-6f, IPI-10f and TCGA-3f were given archetype labels based on the prevalence and distributions of the features within the cluster which were visualized as violin plots of all clusters in each feature.

The IPI-3f and TCGA-3f final clustering selection had six clusters and hence six archetypes. Survival analysis was performed on TCGA-3f per indication and across all indications using lifelines. Pie charts were made to compare the archetype abundance per indication in IPI-3f and TCGA-3f. Additionally, 3D plots of IPI-3f score and TCGA-3f score visualized the distribution of the unclustered scores.

The IPI-6f and IPI-10f final clustering selection had eight and twelve clusters respectively resulting in eight archetypes for IPI-6f and twelve archetypes for IPI-10f. All the final clustering selections (IPI-3f, IPI-6f, IPI-10f,TCGA-3f) were visualized as UMAPs (McInnes et al., 2018), a dimensionality reduced projection of the clustering. The min-dist UMAP parameter, which is the minimum distance between points, controls how close together points are placed in the low-dimensional space of the UMAP. For all the UMAPs we used a value of 0.25 for min-dist. Population flow proportions and phenotype gene signatures were overlayed on the UMAP clustering. Alluvial plots were made using RAWgraphs (Mauri et al., 2017) to visualize the stability of clusters as the clustering progressed from IPI-3f to IPI-6f to IPI-10f.

